# Dysregulation of M segment gene expression contributes to influenza A virus host restriction

**DOI:** 10.1101/599886

**Authors:** Brenda M. Calderon, Shamika Danzy, Gabrielle K. Delima, Nathan T. Jacobs, Ketaki Ganti, Megan R. Hockman, Graeme L. Conn, Anice C. Lowen, John Steel

**Affiliations:** 1 Departments of Microbiology and Immunology, Emory University School of Medicine, Atlanta, GA, USA; 2 Departments of Microbiology and Biochemistry, Emory University School of Medicine, Atlanta, GA, USA

## Abstract

The M segment of the 2009 pandemic influenza A virus (IAV) has been implicated in its emergence into human populations. To elucidate the genetic contributions of the M segment to host adaptation, and the underlying mechanisms, we examined a panel of isogenic viruses that carry avian- or human-derived M segments. Avian, but not human, M segments restricted viral growth and transmission in mammalian model systems, and the restricted growth correlated with increased expression of M2 relative to M1. M2 overexpression was associated with intracellular accumulation of autophagosomes, which was alleviated by interference of the viral proton channel activity by amantadine treatment. As M1 and M2 are expressed from the M mRNA through alternative splicing, we separated synonymous and non-synonymous changes that differentiate human and avian M segments and found that dysregulation of gene expression leading to M2 overexpression diminished replication, irrespective of amino acid composition of M1 or M2. Moreover, in spite of efficient replication, virus possessing a human M segment that expressed avian M2 protein at low level did not transmit efficiently. We conclude that (i) determinants of transmission reside in the IAV M2 protein, and that (ii) control of M segment gene expression is a critical aspect of IAV host adaptation needed to prevent M2-mediated dysregulation of vesicular homeostasis.

**Author summary:** Influenza A virus (IAV) pandemics arise when a virus adapted to a non-human host overcomes species barriers to successfully infect humans and sustain human-to-human transmission. To gauge the adaptive potential and therefore pandemic risk posed by a particular IAV, it is critical to understand the mechanisms underlying viral adaptation to human hosts. Here, we focused on the role of one of IAV’s eight gene segments, the M segment, in host adaptation. Comparing the growth of IAVs with avian- and human-derived M segments in avian and mammalian systems revealed that the avian M segment restricts viral growth specifically in mammalian cells. We show that the mechanism underlying this host range phenotype is a dysregulation of viral gene expression when the avian IAV M segment is transcribed in mammalian cells. In particular, excess production of the M2 protein results in viral interference with cellular functions on which the virus relies. Our results therefore reveal that the use of cellular machinery to control viral gene expression leaves the virus vulnerable to over- or under-production of critical viral gene products in a new host species.

## Introduction

Influenza A virus (IAV) epidemics and pandemics result in widespread and often severe disease [1–3], as well as considerable societal economic costs [4]. IAV lineages endemic in humans are ultimately derived from those circulating in wild waterfowl. Sporadic transmission of IAV from avian reservoirs to humans occurs mainly through domesticated intermediate hosts such as chickens and pigs [5]. For example, infection of humans with H7N9 and H5N1 subtype IAVs occurs through direct exposure to infected poultry. Although they often cause severe disease, these zoonotic cases have not led to sustained onward transmission to date and have therefore not caused a pandemic. Despite abundant circulation of IAVs at the animal-human interface, pandemics occur only rarely owing to the host specificity of IAV infection. As with all viruses, IAV must exploit host cell functions and overcome antiviral barriers to execute its life cycle. Viral replication in a new host species therefore requires adaptation to re-establish virus-host interactions broken by species-specific features of host cellular processes. IAV can undergo such adaptation through reassortment of intact gene segments between avian IAVs and those already adapted to mammals or through more incremental changes brought about by polymerase error [6]. With the goal of anticipating and mitigating IAV emergence in humans, it is essential to understand the species-specific barriers to infection and the mechanisms by which IAV evolution allows these impediments to be overcome.

Changes to viral receptor specificity and polymerase function necessary to overcome host species barriers have been well documented [7–11], but it is clear that additional features of avian IAVs restrict their growth in mammalian systems [12–14]. Seminal studies, performed in the 1970s, indicated a potential role for the M segment. Avian IAVs were shown to express low levels of M1 protein during abortive infection of mammalian cells, but not avian cells [15–17], and the limited quantity of viral particles formed during such abortive infections correlated with the amount of M1 protein expressed. Significantly, these studies were conducted prior to the identification of a second gene product expressed from a spliced form of the M segment mRNA: the M2 protein [18,19]. Subsequent studies demonstrated that low M1 expression during abortive avian IAV infection in mammalian cells was accompanied by increased M2 mRNA and protein expression, relative to productive infection in avian cells, leading to new hypotheses regarding the mechanisms leading to abortive outcomes [20–22].

Additional evidence for a role of the M segment in host range comes from the *in silico* identification of positive selection acting on the M segment of the Eurasian avian-like swine lineage, following transfer of this IAV from birds to swine [23]. Moreover, others and we have shown that the M segment of the 2009 pandemic H1N1 (pH1N1) strain carries determinants of transmissibility, suggesting that adaptation of this segment to humans contributed to the emergence of the pH1N1 lineage [24–28]. The mechanistic basis for this contribution; the specific roles, if any, of M1 and M2 proteins to host range; and whether the M segment plays a broader role in IAV host adaptation, remain unclear.

It is now recognized that the M segment of IAV encodes the M1 matrix and the M2 proton channel proteins via alternative splicing (**Suppl. Figure 1A**) [29,30]. M1 forms a structural layer under the viral envelope and is a major determinant of virion morphology [24,26,31,32]. Upon endocytosis of virus particles into cells, low pH and high [K+] trigger a conformational change in M1 required for release of viral ribonucleoprotein (vRNP) complexes into the cytoplasm [33]. M1 is also critical for the transport of newly replicated vRNPs from the nucleus to the cytoplasm [34–37]. At late stages of infection, cytoplasmic M1 is thought to recruit vRNP complexes to the budding site [38–40].

M2 is a transmembrane protein that tetramerizes to form a proton channel. Within the envelope of incoming virions, this channel allows diffusion of protons into the virion core, thereby triggering the release of vRNPs from M1, as well as facilitating the intracellular transport of HA protein [41–44]. M2 also plays an important role in the final stages of virion morphogenesis, mediating ESCRT-independent membrane scission [45]. Importantly for the findings reported herein, multiple groups have reported that M2 blocks the fusion of autophagosomes with lysosomes [46–48]. Under normal conditions, this fusion event allows degradation and recycling of organelles and protein aggregates contained within autophagosomes [49], and is thus important for cellular homeostasis. M2-mediated interference with autolysosome formation therefore stalls autophagic flux and leads to the accumulation of autophagosomes within the cell [46–48].

Many elegant molecular virology studies have revealed that viral strategies for regulating gene expression are central to orchestration of the viral life cycle [50–53]. Here, we report that regulation of M2 expression of IAV is an important determinant of host range. In particular, our data show that the expression of M1 and M2 from alternatively spliced M segment mRNAs is dysregulated when an avian IAV replicates in mammalian cells, and that the resultant overexpression of the M2 proton channel attenuates viral growth by disrupting the cellular autophagy pathway. The observed block in autophagy is consistent with that reported previously for lab-adapted strains of IAV [46–48]. Importantly, however, we see that lab-adapted IAVs express M2 at high levels, comparable to those seen for avian IAV M segments in mammalian cells. In contrast, IAVs carrying M segments well adapted to mammalian systems express relatively little M2 and do not trigger the accumulation of autophagosomes to the same extent as avian-origin and lab-adapted strains. These findings suggest that IAV-induced stalling of autophagic turnover is a result of aberrantly high M2 expression upon infection of mammalian cells with avian IAVs, and which is overcome in human-adapted influenza A lineages. Critically, the mechanism leading to this host-restricted phenotype occurs at the level of gene expression and is driven by host-specific differences in viral nucleic acid sequence, not amino acid changes. This mechanism constitutes a novel paradigm in RNA virus host adaptation, and reveals a new species barrier for IAV, which may be highly relevant for the emergence of avian IAVs into humans.

## Results

### Generation of isogenic IAVs differing only in the M segment

To assess the contribution of the M segment to host adaptation, we generated a panel of IAVs possessing human- or avian-derived M segments in the A/Puerto Rico/8/34 (H1N1) [PR8] background (**Table 1; Figure 1A; Suppl. Figure 1B, C, D**). The M segments used were selected with the aims of representing circulating human IAVs and capturing the breadth of diversity of M segment sequences present in wild waterfowl. In addition to three wild-type avian M segments, we included in our panel a mutant of the dk/Alb/76 M segment that carries a uniquely identified mutation in M2 (S89). Reversion of this unique amino acid residue to glycine yields an M2 protein matching the consensus amino acid sequence of IAVs in wild bird reservoirs. The human-adapted M segment analyzed most extensively herein is that of the pandemic A/NL/602/09 (H1N1) strain [NL09].

**Table 1.**
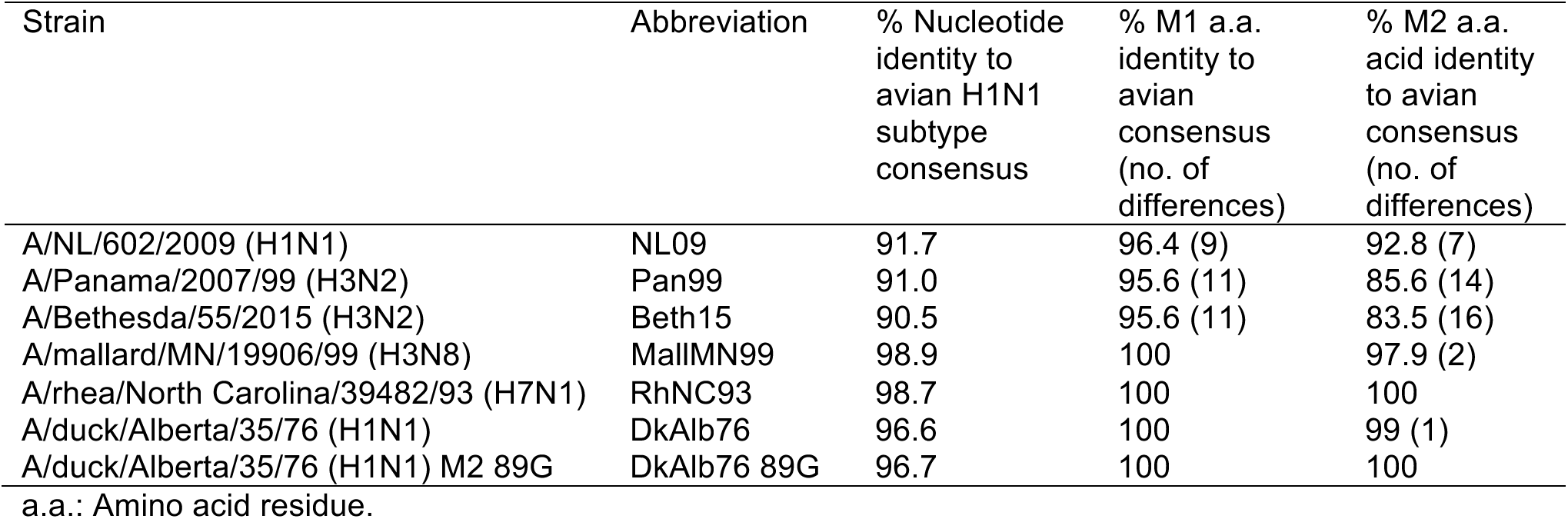
Origin of IAV M segments used in study.

**Figure 1.**
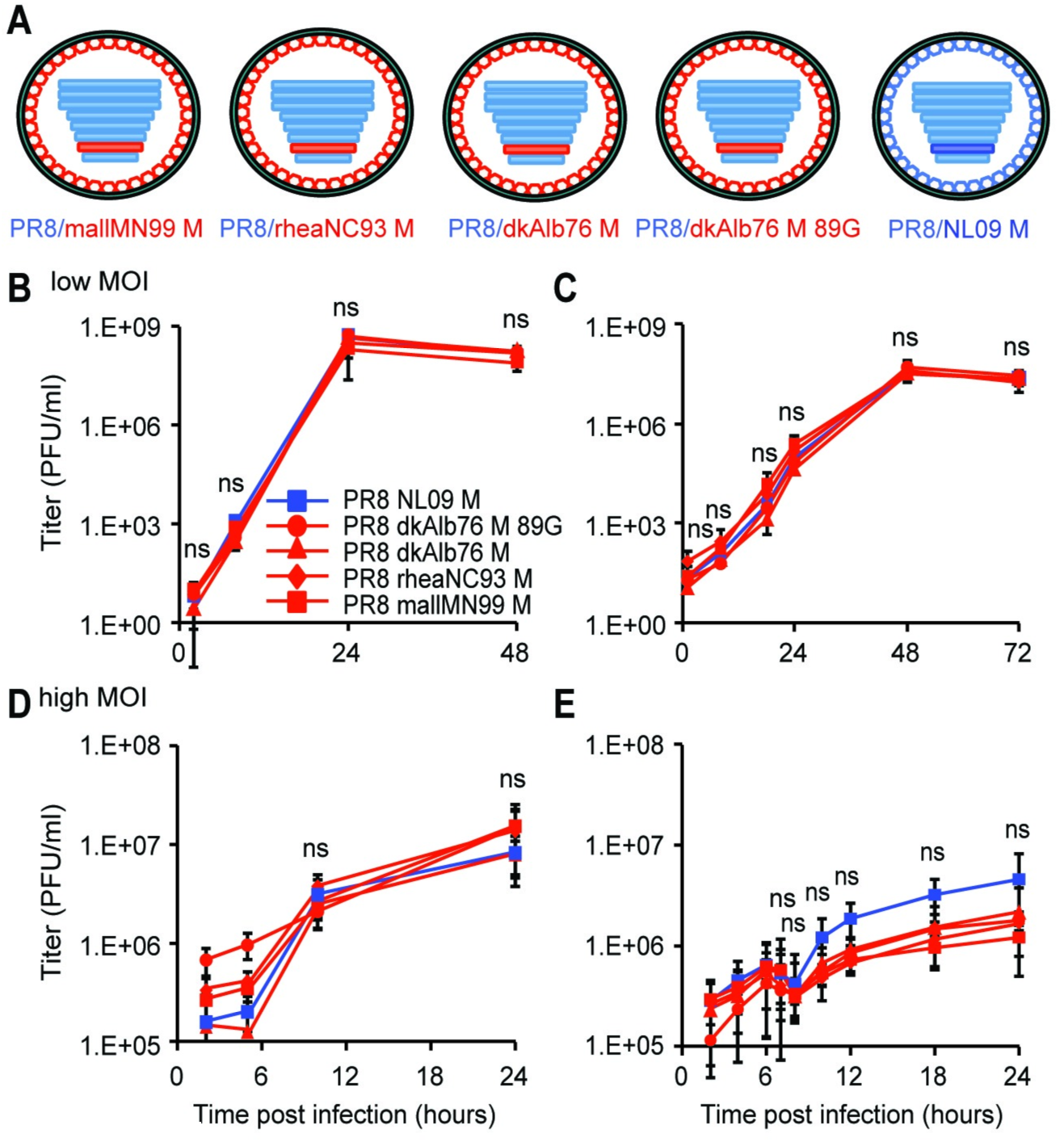
pH1N1 M segment does not confer improved growth relative to avian M segment in avian cells at 37°C. **(A)** A panel of PR8-based viruses was rescued by reverse genetics. The viruses differ only in the identity of the M segment, which were derived from A/mallard/MN/99 (H3N8), A/rhea/NC/93 (H7N1), A/duck/Alberta/76 (H1N1), or A/NL/602/09 (H1N1) viruses. **(B)** 11-day-old embryonated chicken eggs (ECE) or (**C**) chicken DF-1 cells were infected at low MOI with the indicated viruses. **(D)** Quail QT-6 cells or **(E)** DF-1 cells were infected at a high MOI of 5 PFU/cell with the indicated viruses. In all avian systems assessed, the pH1N1 M segment did not confer an increase in magnitude or kinetics of growth, relative to any avian-origin M segment. Multi-cycle and single-cycle growth were assessed in three independent experiments, with three technical replicates per experiment. Graphs show the means with SD for the three experiments. Statistical significance was determined using repeated measures, two-way, multiple ANOVA on log-transformed values, with Bonferroni correction applied to account for comparison of a limited number of means.

### IAV M segments of avian origin confer a host-restricted phenotype

The relative fitness of PR8-based viruses carrying M segments from the human-adapted NL09 virus or from avian IAVs was evaluated in three avian substrates: embryonated chicken eggs (ECE), chicken origin DF-1 cells, and quail origin QT-6 cells. When grown from low multiplicities of infection (MOI) in either ECE or DF-1 cell substrates, all viruses exhibited similar kinetics and reached comparable titers (**Figure 1B and 1C**). Also, under high MOI growth conditions in DF-1 and QT-6 cells, the growth phenotypes of the PR8 NL09 M and PR8 avian M viruses were similar, although the NL09 M-containing virus trended towards increased growth relative to the avian M-containing viruses in DF-1 cells (**Figure 1D and 1E**). Nonetheless, at no time did the NL09 M-encoding virus grow to significantly higher titers than the avian M-encoding viruses in any avian host substrate. In contrast, in mammalian cells, human-adapted IAV M segments were found to support improved growth relative to avian-adapted M segments. Viral growth was monitored from low and high MOI in human A549 cells and canine MDCK cells (**Figure 2**), and additionally from high MOI in human 293T cells (**Suppl. Figure 2A**). In each case, PR8-based viruses carrying the NL09 M segment grew with more rapid kinetics and to higher titers than isogenic viruses carrying avian M segments, although the differences in growth from low MOI did not reach statistical significance. To extend our findings to an independent human lineage of IAV, we evaluated the growth from high MOI in A549 cells of PR8 Pan99 M and PR8 Beth15 M viruses, each of which have an M segment derived from the human seasonal H3N2 lineage (**Suppl. Figure 1C, D**). We found that both of these human-adapted M segments supported significantly faster kinetics and higher magnitude of growth than the dkAlb76 89G M segment (**Suppl. Figure 2B**). Thus, results from multiple different avian and mammalian culture systems reveal defects in the kinetics and magnitude of viral growth associated with avian IAV M segments specifically in mammalian host cells and provide evidence of the acquisition of adaptive changes in the M segment of pH1N1 and H3N2 human influenza A lineages.

**Figure 2.**
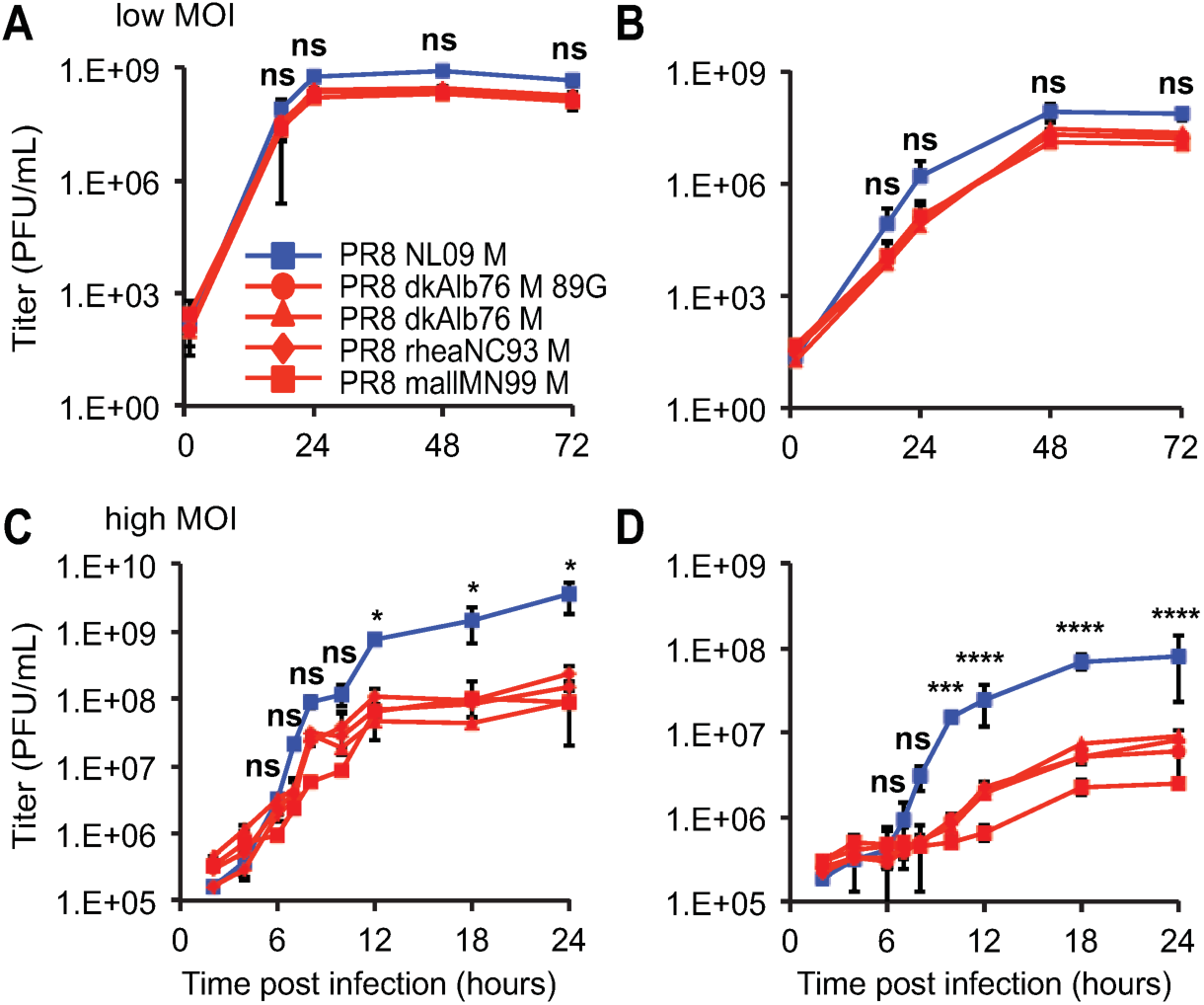
pH1N1 M segment confers higher growth relative to avian M segment in mammalian cells at 37°C. **(A)** Canine MDCK cells or **(B)** human A549 cells were inoculated with the indicated viruses at a low MOI of 0.01 PFU/cell. **(C)** MDCK cells or **(D)** A549 cells were inoculated with the indicated viruses at a high MOI of 5 PFU/cell. In mammalian cells, the pH1N1 M segment conferred more rapid kinetics and higher peak titers than any avian-origin M segment. Multi-cycle and single-cycle growth were assessed in three independent experiments, with three technical sample replicates per experiment. Graphs show the means with SD for the three experiments. Statistical significance was determined using repeated measures, two-way, multiple ANOVA on log-transformed values, with Bonferroni correction applied to account for comparison of a limited number of means. At each time point, pairwise comparisons between viruses were performed, and asterisks corresponding to the most significant difference observed is shown.

### Avian IAV M segments confer lower viral infectivity, growth and transmission than NL09 M segment in a guinea pig model

To evaluate the impact of M segment host adaptation on viral phenotypes in an intact host, we evaluated the PR8 NL09 M virus and the four PR8-based viruses carrying avian M segments in a guinea pig model [54,55]. A low inoculation dose of 10 PFU per animal was used to maximize the sensitivity of the system. Each inoculated animal was co-caged with a naïve guinea pig at 24 h post-inoculation and viral infectivity, magnitude and kinetics of growth, and transmission were monitored by collecting nasal lavage samples every other day. Among those animals that were productively infected, total virus sampled in nasal washes over the course of infection was evaluated by calculating the area under the shedding curves shown in **Figure 3A**. Area under the curve values did not differ significantly among the four viruses with avian M segments, but were significantly higher for PR8 NL09 M virus (**Figure 3A**). As shown in **Figure 3B**, fewer guinea pigs were productively infected with 10 PFU when the inoculating virus carried an avian M segment. Differences in the kinetics of viral growth were determined by assessing changes in growth over time using repeated measures ANOVA (**Suppl. Figure 3**). Each virus encoding an avian host-derived M segment replicated with significantly slower kinetics than the PR8 NL09 M virus.

**Figure 3.**
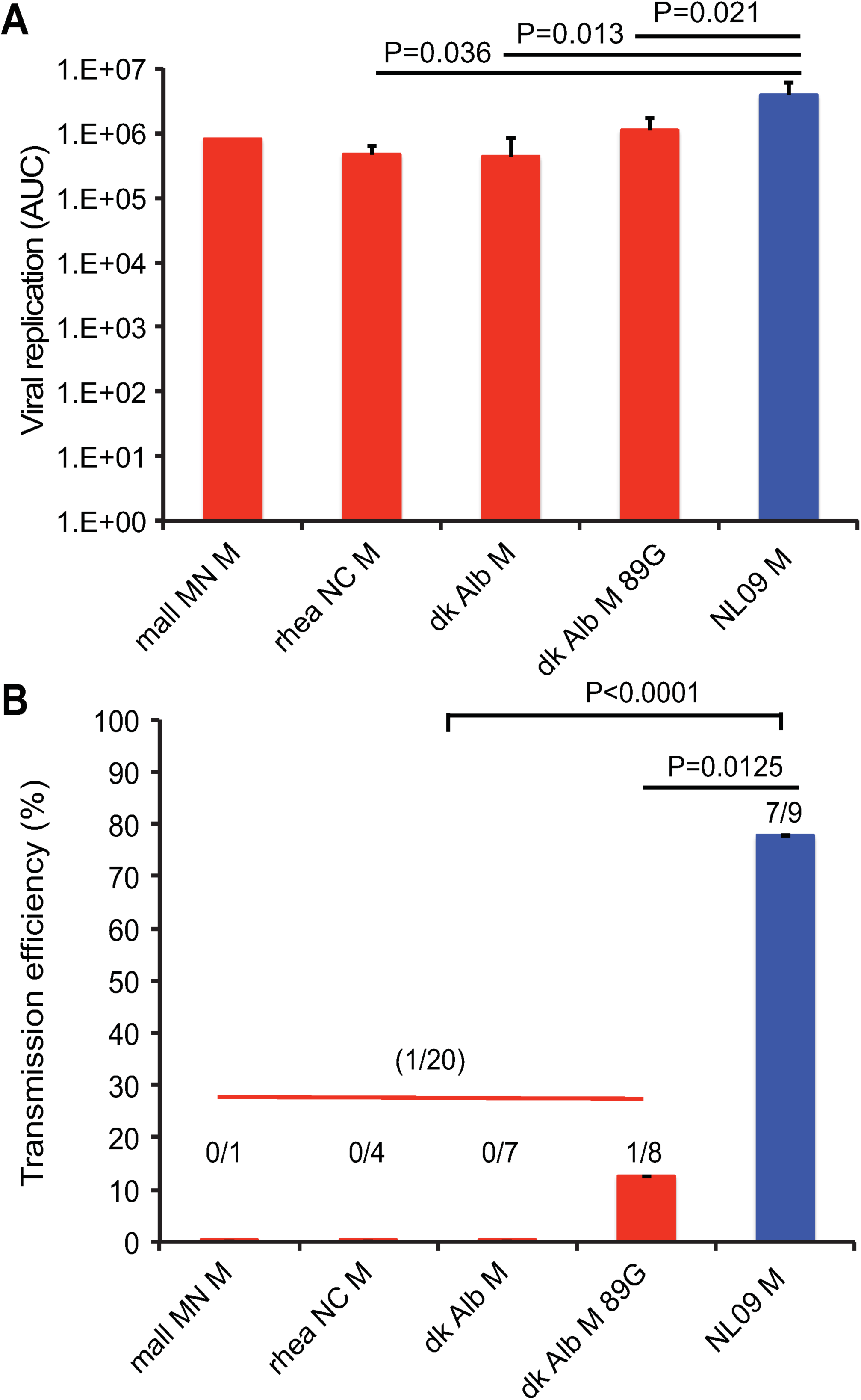
pH1N1 influenza A virus M segment confers efficient replication and transmission of PR8-based virus among guinea pigs. Groups of four guinea pigs were inoculated with 10 PFU of each avian M-encoding virus or NL09 M-encoding virus, as indicated. **(A)** Virus replication in nasal wash was measured by plaque titration at days 2, 4, 6, and 8 post-infection and the area under the curve was calculated. Graphs show mean AUC with SD for three experiments. All avian M-encoding viruses exhibited lower levels of growth *in vivo* than the isogenic virus possessing the NL09 M segment. The replication differences between avian M and NL09 M-based viruses were statistically significant, whereas those among avian M-based viruses were not. Statistical significance was evaluated using an unpaired, two-tailed Student’s t-test. **(B)** To assess transmission, each inoculated guinea pig was treated as an independent biological sample. The pH1N1 M segment conferred efficient transmission to naïve animals that were contact exposed (7/9 transmissions; 78%). In contrast, significantly poorer (12%), or no transmission was observed from guinea pigs infected with PR8 avian M viruses to naïve cage mates. Statistical significance of transmission differences were evaluated using an unpaired, two-tailed Student’s t-test.

The efficiency of transmission was assessed by calculating the proportion of contact animals that contracted infection, considering only cages in which donor guinea pigs were productively infected through intranasal inoculation. While transmission occurred in 7/9 instances for PR8 NL09 M virus, a significantly lower proportion of animals (1/20; p<0.0001) were infected with a virus carrying an avian M segment transmitted from her cage mate (**Figure 3B**). Thus, data from a mammalian animal model reveal defects in infectivity, growth and transmission associated with avian IAV M segments. Our results suggest the acquisition of adaptive changes in the M segment of pH1N1 human IAV since its introduction from the avian reservoir into mammals.

### The M2 proton channel is aberrantly overexpressed from avian IAV M segments in mammalian cells

To investigate the mechanism underlying the host-restricted phenotypes conferred by avian-adapted M segments, the steady state levels of the M1 and M2 proteins were evaluated in human A549 and avian DF-1 cells infected at high MOI. Western immunoblotting with the monoclonal antibody E10, which recognizes the shared N-terminus of M1 and M2, was used to monitor levels of both proteins concurrently.

In the avian cell line, the PR8 NL09 M virus expressed lower levels of M2 than M1 (**Figure 4A**). When M1 and M2 levels were normalized to those of vinculin and the relative levels of M1 and M2 calculated, the NL09 M2 protein comprised <10% of the total M segment-derived protein expression. Very similar results were seen for four PR8-based viruses carrying avian M segments grown in DF-1 cells. In each case, M2 comprised <20% of the total M segment-derived protein, and in the case of three avian M segment-containing viruses (PR8 dkAlb76 M; PR8 dkAlb76 M 89G; and PR8 rheaNC93 M), the levels of M1 and M2 proteins were not significantly different from those expressed by the PR8 NL09 M virus (**Figure 4B, 4C, 4D**).

**Figure 4.**
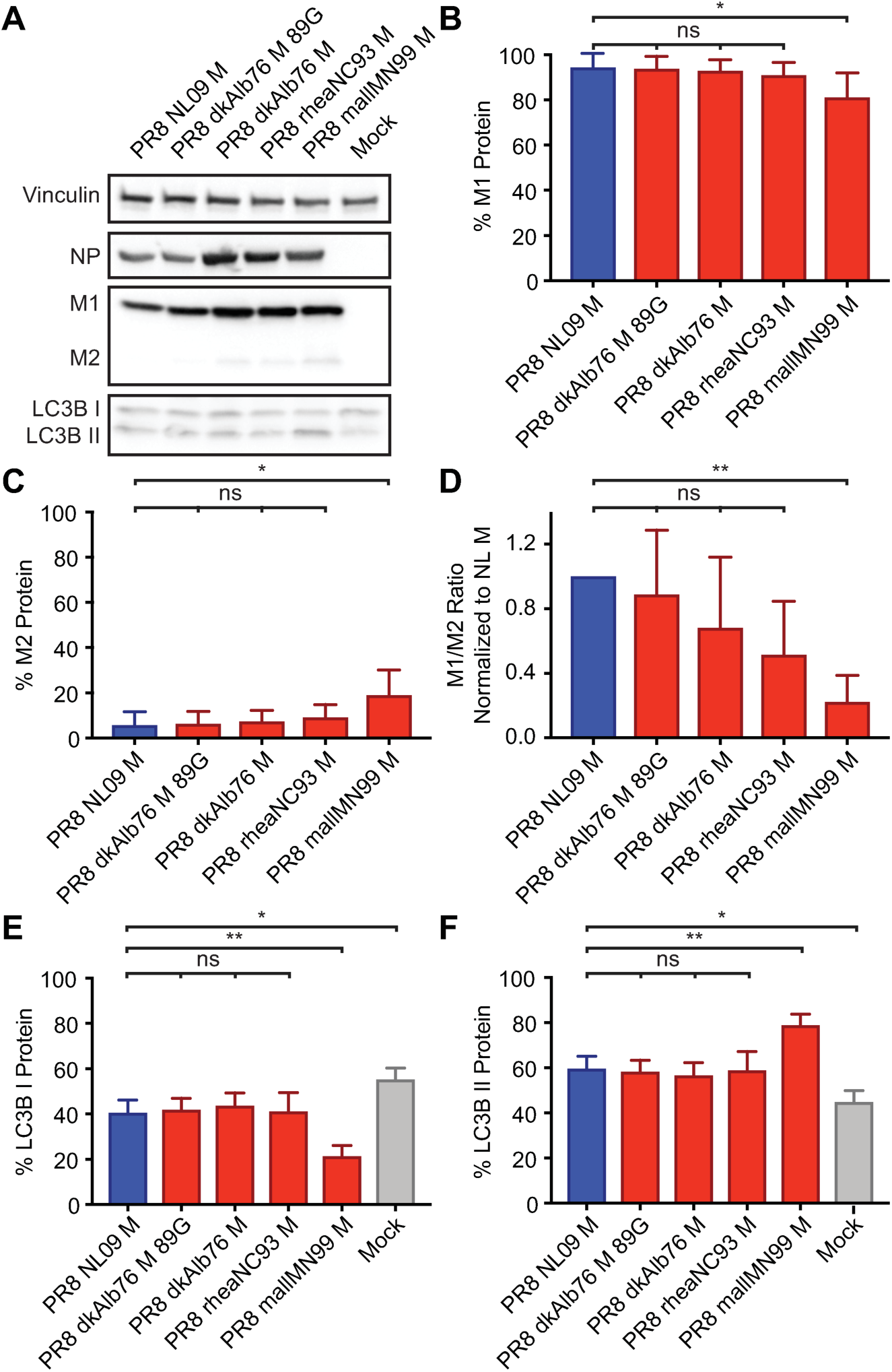
Ratio of M1 to M2 protein expression is high in DF-1 Cells irrespective of viral M segment host origin. DF-1 cells were inoculated at a MOI of 5 PFU/cell, with PR8 viruses encoding avian or human-derived M segments, and incubated at 37°C for 8 h, then lysed. Western immunoblot analysis of virus-infected DF-1 cells: **(A)** Vinculin expression was measured to allow normalization of viral protein levels. NP expression was measured to assess viral replication. Levels of M1 and M2 protein expression were assessed using an antibody (Mab E10) to a common epitope at the amino terminus of M1 and M2 proteins, allowing relative expression to be assessed. Levels of LC3B I and II were assessed using an antibody that detects both the precursor and activated forms of LC3B protein. **(B)** M1 protein and **(C)** M2 protein were normalized to vinculin, quantitated and displayed as a percentage of total protein expressed from the M gene. **(D)** The ratio of M1:M2 protein expression**. (E)** LC3B I protein and **(F)** LC3B II protein were normalized, quantitated and displayed as a percentage of total LC3B protein. Graphs in **B-F** show the means with SD from three independent experiments. For each experiment, two replicate Western immunoblots were performed and quantitated. Statistical significance was assessed using ordinary one-way ANOVA.

In A549 cells, the PR8 NL09 M virus expressed M2 at ∼20% of the total M protein, similar to the expression levels observed in infected DF-1 cells (**Figure 5A**). In contrast, avian M segments yielded markedly higher levels of the M2 protein than were observed for the same viruses grown in avian cells. Here, M2 levels were approximately equal to those of M1 (**Figure 5B, 5C, 5D**). Moreover, when compared to the NL09 M segment in human cells, levels of M2 protein were significantly higher for avian M segments (P<0.0001) (**Figure 5C, 5D**). Thus, compared to matched virus-host pairings in both human and avian systems, M2 was markedly overexpressed when avian-adapted M segments were introduced into human cells.

**Figure 5.**
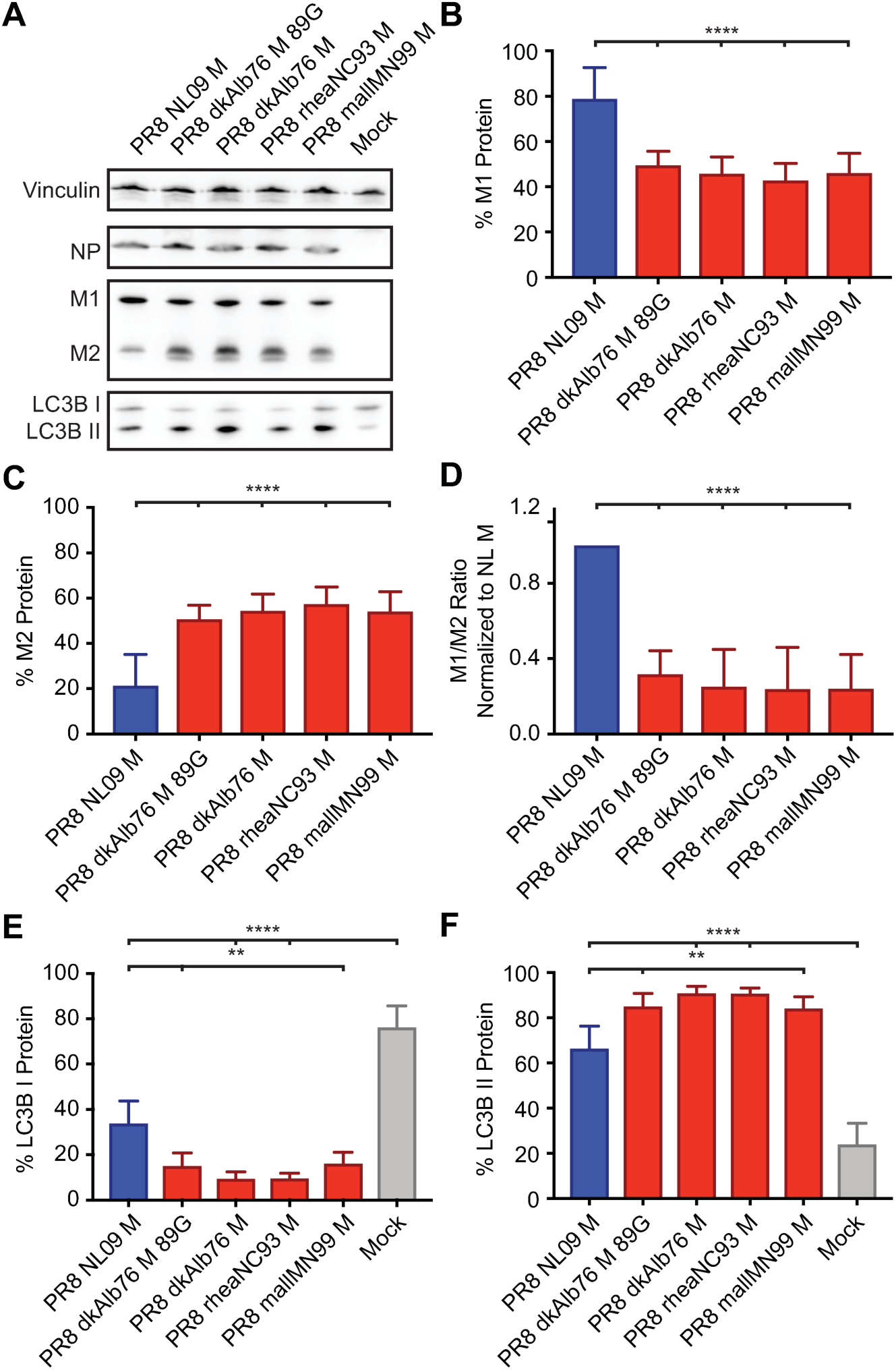
High expression ratio of M1 to M2 protein in A549 cells is dependent upon viral M segment host origin. A549 cells were inoculated at a MOI of 5 PFU/cell with PR8 viruses encoding avian or human-derived M segments and incubated at 37°C for 8 h, then lysed. Western immunoblot analysis of virus-infected A549 cells: **(A)** Vinculin expression was measured to allow normalization of viral protein levels. NP expression was measured to assess viral replication. Levels of M1 and M2 protein expression were assessed using an antibody (Mab E10) to a common epitope at the amino terminus of M1 and M2 proteins, allowing relative expression to be assessed. Levels of LC3B I and II were assessed using an antibody that detects both the precursor and activated forms of LC3B protein. **(B)** M1 protein and **(C)** M2 protein were normalized to vinculin, quantitated and displayed as a percentage of total protein expressed from the M gene. **(D)** The ratio of M1:M2 protein expression. **(E)** LC3B I protein and **(F)** LC3B II protein were normalized, quantitated and displayed as a percentage of total LC3B protein. Graphs in **B-F** show the means with SD from three independent experiments. For each experiment, two replicate Western immunoblots were performed and quantitated. Statistical significance was assessed using ordinary one-way ANOVA.

### Overexpression of M2 protein in mammalian cells corresponds to increased levels of the M2 mRNA (mRNA 10)

The M segment vRNA is transcribed by the viral polymerase to give rise to a co-linear mRNA (mRNA_7_), which can be spliced by cellular splicing factors to yield mRNA_10_ or mRNA_11_[18,56]. The M1 protein is translated from the unspliced mRNA_7_, while M2 is expressed from the spliced mRNA_10_. To date, no polypeptide corresponding to the short mRNA_11_ ORF has been identified. Aberrantly high levels of IAV mRNA_10_ in mammalian cells have been previously reported for the avian IAV, fowl plague virus [20,21]. We therefore hypothesized that the high expression levels of M2 protein, observed for avian-adapted M segments in human cells, was due to increased splicing of mRNA_7_ to yield excessive mRNA_10_. We used a RT primer extension assay to quantify the levels of each M segment-derived mRNA species in DF-1 and A549 cells infected with either PR8 NL09 M virus or one of four viruses carrying an avian-adapted M segment.

As expected, based on the levels of the encoded proteins, mRNA_10_ was of low abundance compared to mRNA_7_ for all viruses in DF-1 cells (**Figure 6A**). Quantification of band intensities revealed that, in DF-1 cells infected with PR8 NL09 M virus, mRNA_7_ comprised ∼75%, while mRNA_10_ comprised ∼5% of the total M segment-derived mRNA (**Figure 6B**). For three avian M viruses (PR8 dkAlb76 M; PR8 dkAlb76 M 89G; and PR8 rheaNC93 M), relative levels of mRNA_7_ were ∼70-75% and were not significantly different from those of PR8 NL09 M virus (**Figure 6B**). For all viruses, mRNA_10_ was expressed at low levels (10-40% of total segment 7 mRNA), and in the case of two avian M segment-containing viruses (PR8 dkAlb76 M; PR8 dkAlb76 M 89G), the levels of mRNA_10_ were not significantly different from those expressed by the PR8 NL09 M virus (**Figure 6C**). mRNA_11_ was expressed in each infection at approximately 20% of the total M segment-derived mRNA, and the relative level did not differ significantly with any virus (**Figure 6D**). Overall, the relative levels of each of the three mRNA species were similar when synthesized from avian- or human-origin encoded M segments in DF-1 cells, correlating with Western immunoblot data on M segment protein expression in these cells.

**Figure 6.**
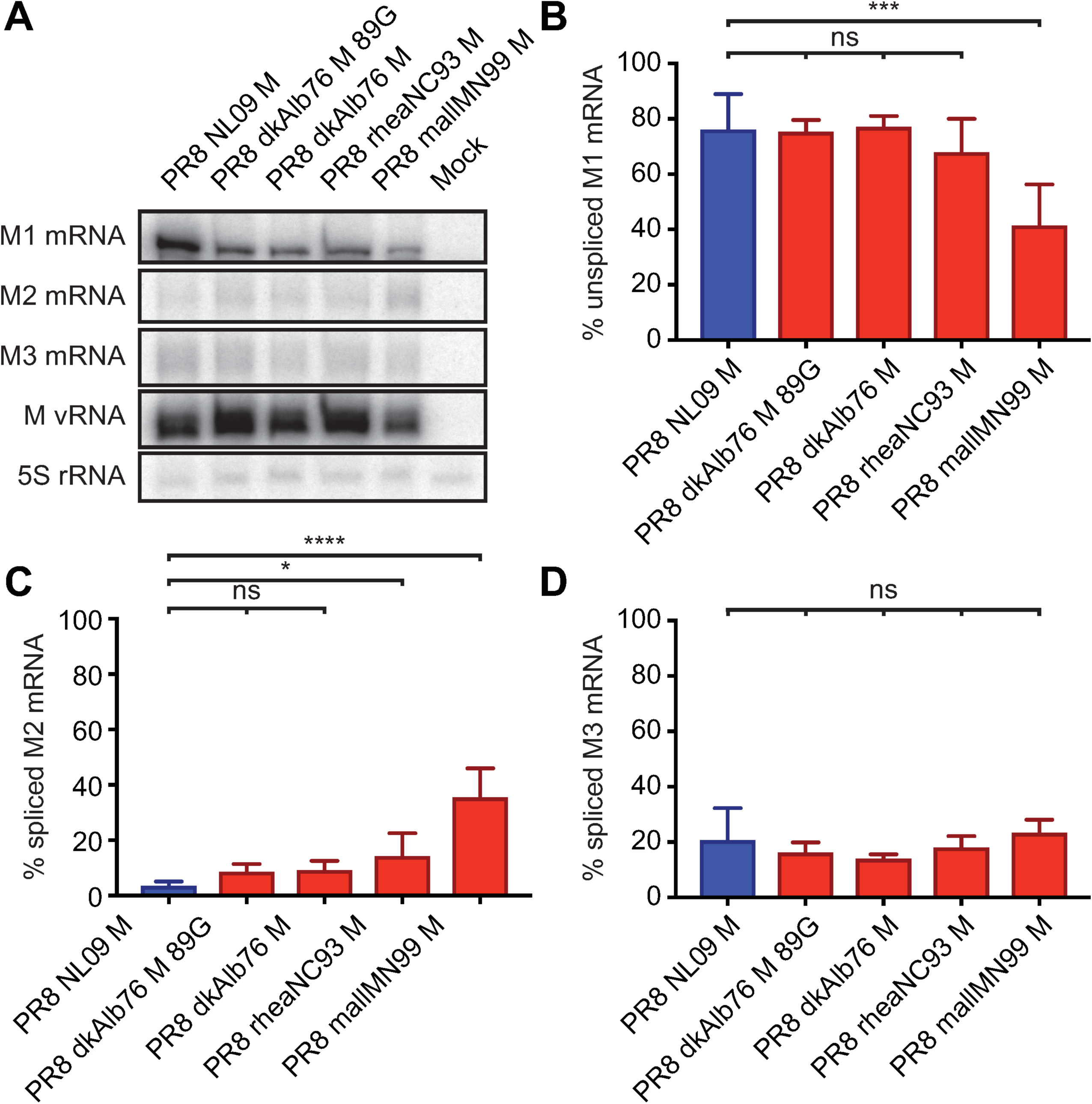
Ratio of mRNA_7_ (encoding M1) to mRNA_10_ (encoding M2) expression is high in DF-1 cells irrespective of viral M Segment host origin. DF-1 cells were inoculated at a MOI of 5 PFU/cell with PR8 viruses encoding avian or human-derived M segments and incubated at 37°C for 8 h prior to RNA extraction. RT primer extension radiogram of virus-infected DF-1 cells: **(A)** 5S rRNA levels were measured to allow normalization of viral RNA. Segment 7 vRNA expression was measured to assess viral replication. Levels of mRNA_7_, mRNA_10_, and mRNA_11_ mRNA expression were assessed using radiolabeled probes. **(B)** mRNA_7_, **(C)** mRNA_10_, and **(D)** mRNA_11_ was quantitated and displayed as a percentage of total M gene-expressed mRNA. Graphs in **B-D** show the means with SD from three independent experiments. For each experiment, two replicate radiograms were quantitated. Statistical significance was assessed using ordinary one-way ANOVA.

In A549 cells, the PR8 NL09 M virus showed relatively high abundance of mRNA_7_ and low abundance of mRNA_10_, as was seen in DF-1 cells (**Figure 7A**). By contrast, for viruses carrying avian-adapted M segments, mRNA_10_ levels exceeded those of mRNA_7_ in A549 cells (**Figure 7B, 7C**). Both decreases in mRNA_10_ levels and increases in mRNA_7_ levels contributed to the altered relative abundance. Again, no significant differences in relative mRNA_11_ levels among the viruses were noted (**Figure 7D**) suggesting that changes in splicing to produce mRNA_11_ do not contribute to changes in M segment mRNA and protein expression observed in A549 cells. Overall, our data point to increased splicing to produce mRNA_10_ as the underlying mechanism leading to heightened M2 expression from avian M segments in A549 cells, and suggest that cis-acting signals present on the M segment are responsible for driving the disparate M2 expression levels.

**Figure 7.**
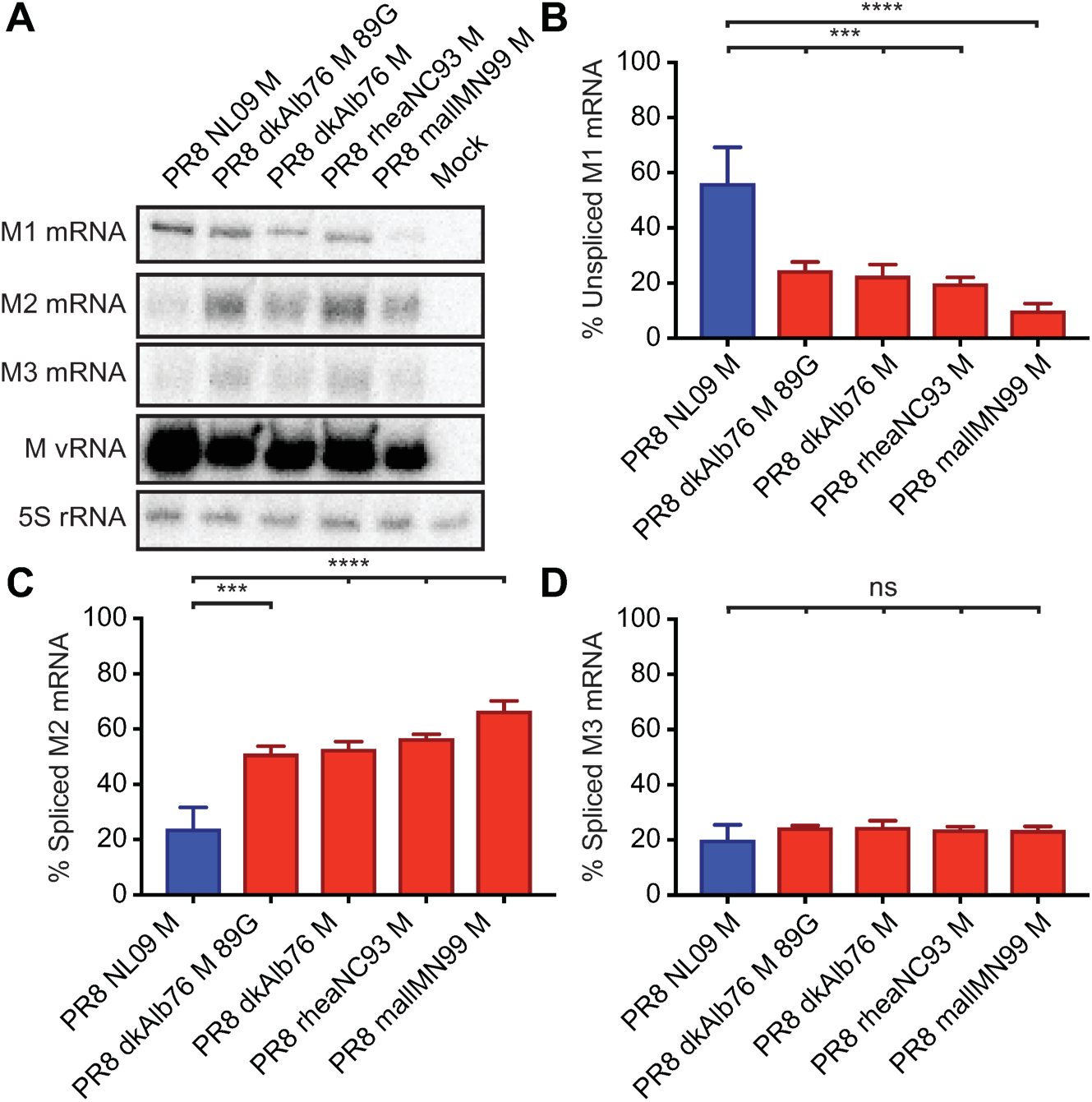
Ratio of mRNA_7_ (encoding M1) to mRNA_10_ (encoding M2) expression in A549 cells is dependent upon viral M segment host origin. A549 cells were inoculated at a MOI of 5 PFU/cell with PR8 viruses encoding avian or human-derived M segments and incubated at 37°C for 8 h prior to RNA extraction. RT primer extension radiogram of virus-infected A549 cells: **(A)** 5S rRNA levels were measured to allow normalization of viral RNA. Segment 7 vRNA expression was measured to assess viral replication. Levels of mRNA_7_, mRNA_10_, and mRNA_11_ mRNA expression were assessed using radiolabeled probes. **(B)** mRNA_7_, **(C)** mRNA_10_, and **(D)** mRNA_11_ was quantitated and displayed as a percentage of total M gene expressed mRNA. Graphs in **B-D** show the means with SD from three independent experiments. For each experiment, two replicate radiograms were quantitated. Statistical significance was assessed using ordinary one-way ANOVA.

### M2 expressed from avian M segments localizes in perinuclear puncta

To gain initial insight into the phenotypic consequences of M2 overexpression, we examined the subcellular localization of M2 in A549 cells infected with PR8 NL09 M virus or one of four PR8-based viruses with avian M segments. At 8 hours-post-infection (hpi) in PR8 NL09 M virus-infected cells, M2 showed mainly dispersed cytoplasmic staining, along with weak cytoplasmic membrane localization and some perinuclear accumulation. By contrast, in cells infected with viruses that encoded any of the four avian origin M segments, we noted more abundant M2 protein staining localized at the plasma membrane, and in large puncta near the cell nucleus (**Figure 8A**). Similar data were obtained from virus-infected 293T cells at 8 hpi (**Figure 8B**) and from both human cell lines at 12 hpi (**Suppl. Figure 6**). This staining pattern, along with data in previously published studies [46–48], prompted us to investigate whether highly expressed M2 protein was interacting with the cellular autophagy pathway.

**Figure 8.**
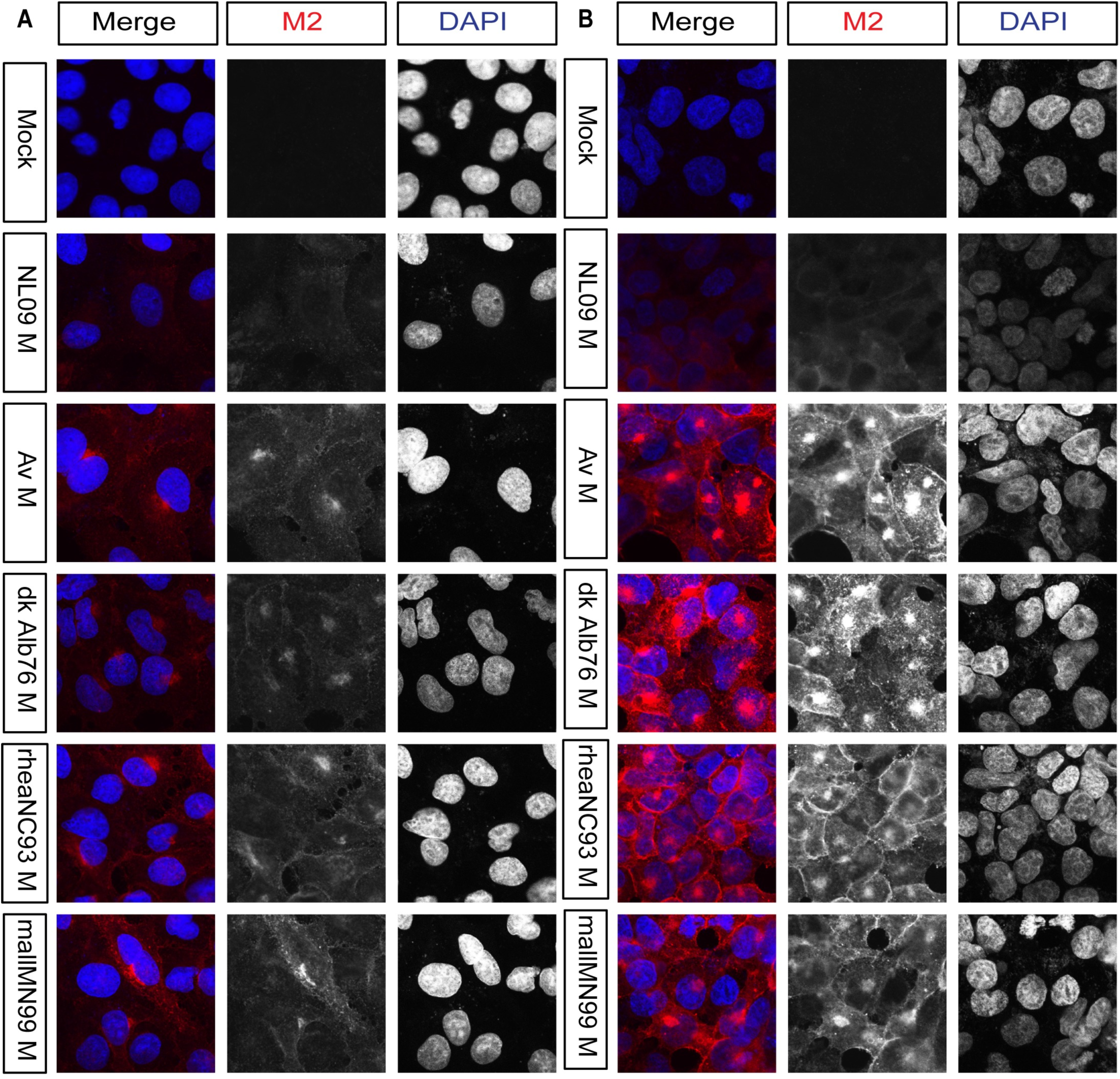
In human cells, avian M2 was expressed at higher levels and localized more strongly to perinuclear bodies than human M2. A549 and 293T cells were inoculated with the indicated viruses, encoding avian or human M segments, at a MOI of 5 PFU/cell. At 8 hpi cells were stained with anti-M2 (Mab E10; red) and DAPI (blue) and imaged by confocal microscopy. Examples of optical sections are shown, either as merged 2-color images or the red and blue channels alone (shown in grey scale). **(A)** A549 cells with 63x optical and 3x digital magnification (189x total magnification). **(B)** 293T cells with 63x optical and 3x digital magnification (189x total magnification). Brightness was adjusted for optimal clarity, with all images treated equally.

### Human cells exhibit increased LC3B II levels when infected with IAV carrying avian M segments

An interaction between the IAV M2 protein and the cellular autophagy pathway has been reported previously [46–48]. To test whether M2 expression from avian vs. human M segments has differential effects on autophagy, we first evaluated the level of LC3B lipidation in PR8 NL09 M- and PR8 avian M-infected cells. Upon activation of the autophagy pathway, LC3B is modified through the addition of phosphotidylethanolamine [57,58]. This lipidated form, called LC3B II, remains associated with the membranes of maturing and mature autophagosomes. Thus, increased levels of LC3B II in the cell are indicative of autophagosome formation; however, under normal conditions LC3B I and II are cycled through an autophagic flux which involves autophagosome maturation and degradation via fusion with lysosomes [59]. A block in this fusion event can therefore lead to accumulation of autophagosomes in the cytoplasm and, consequently, accumulation of LC3B II in the cell.

Western immunoblot analysis of LC3B revealed decreased LC3B I and increased LC3B II protein levels in virus-infected cells compared to mock, providing evidence of autophagy activation and/or block during IAV infection of either avian or human cells (**Figure 4A, 4E, 4F; 5A, 5E, 5F**). In avian DF-1 cells, levels of LC3B I were slightly decreased and levels of LC3B II were slightly increased over mock, but LC3B I and LC3B II levels were generally comparable among human M- and avian M-containing viruses (**Figure 4E, 4F**). Importantly, human A549 cells infected with PR8 avian M-containing viruses consistently showed lower levels of LC3B I, and higher levels of LC3B II than A549 cells infected with PR8 NL09 M as well as other human M-containing viruses (**Figure 5E, 5F**). Additionally, the same phenotypes were obtained from the panel of viruses upon infection of human origin 293T cells (**Suppl. Figure 4A-4F**), or canine origin MDCK cells (**Suppl. Figure 4G-4L**).

Taken together, these data reveal a correlation between levels of M2 expression and LC3B lipidation (accumulation of LC3B II), suggesting that high levels of M2 present in PR8 avian M-infected cells may be inducing an over-activation of autophagy, or a block in the turnover of autophagosomes.

### Infection of human cells with IAV carrying avian M segments triggers accumulation of autophagosomes

To confirm that M2 protein was co-localizing with LC3-positive autophagic vesicles in mammalian cells, we monitored sub-cellular localization of an overexpressed LC3B-GFP fusion protein in infected cells. Human 293T cells were primarily used for this purpose as they transduce well with GFP-LC3. Under normal conditions, LC3B-GFP can be seen throughout the cell with a diffuse distribution (**Figure 9, row 1: mock infected cells**). Treatment with chloroquine, an inhibitor of autophagy that decreases autophagosome-lysosome fusion, results in the concentration of LC3B-GFP into punctate cytoplasmic structures (**Figure 9, row 7: CQ-treated cells**). This relocalization is indicative of the accumulation of autophagosomes [60]. In LC3B-GFP expressing 293T cells infected with PR8 NL09 M virus, we observed little to no increase in GFP-positive cytoplasmic puncta (**Figure 9, row 2: NL09 M infected cells**). In cells infected with PR8 avian M-encoding viruses, by contrast, extensive accumulation was apparent, with both the size and number of intracellular vesicles increased relative to PR8 NL09 M infection (**Figure 9, rows 3-6: Avian M infected cells**). In addition to these cytoplasmic puncta, viruses with avian M segments were found to redirect a small proportion of LC3B-GFP to the plasma membrane. Notably, the M2 protein co-localized with LC3B-GFP in PR8 avian M-infected cells, both in cytoplasmic puncta and at the plasma membrane. Similar phenotypes were observed in virus-infected A549 cells, at 8 hpi (**Suppl. Figure 7**). Overall, these results suggest that expression of avian M2 protein at high levels interferes with the turnover of autophagosomes and with the localization of LC3B-GFP. In addition, the results reveal a reduction in the tight perinuclear localization of M2 upon LC3B overexpression.

**Figure 9.**
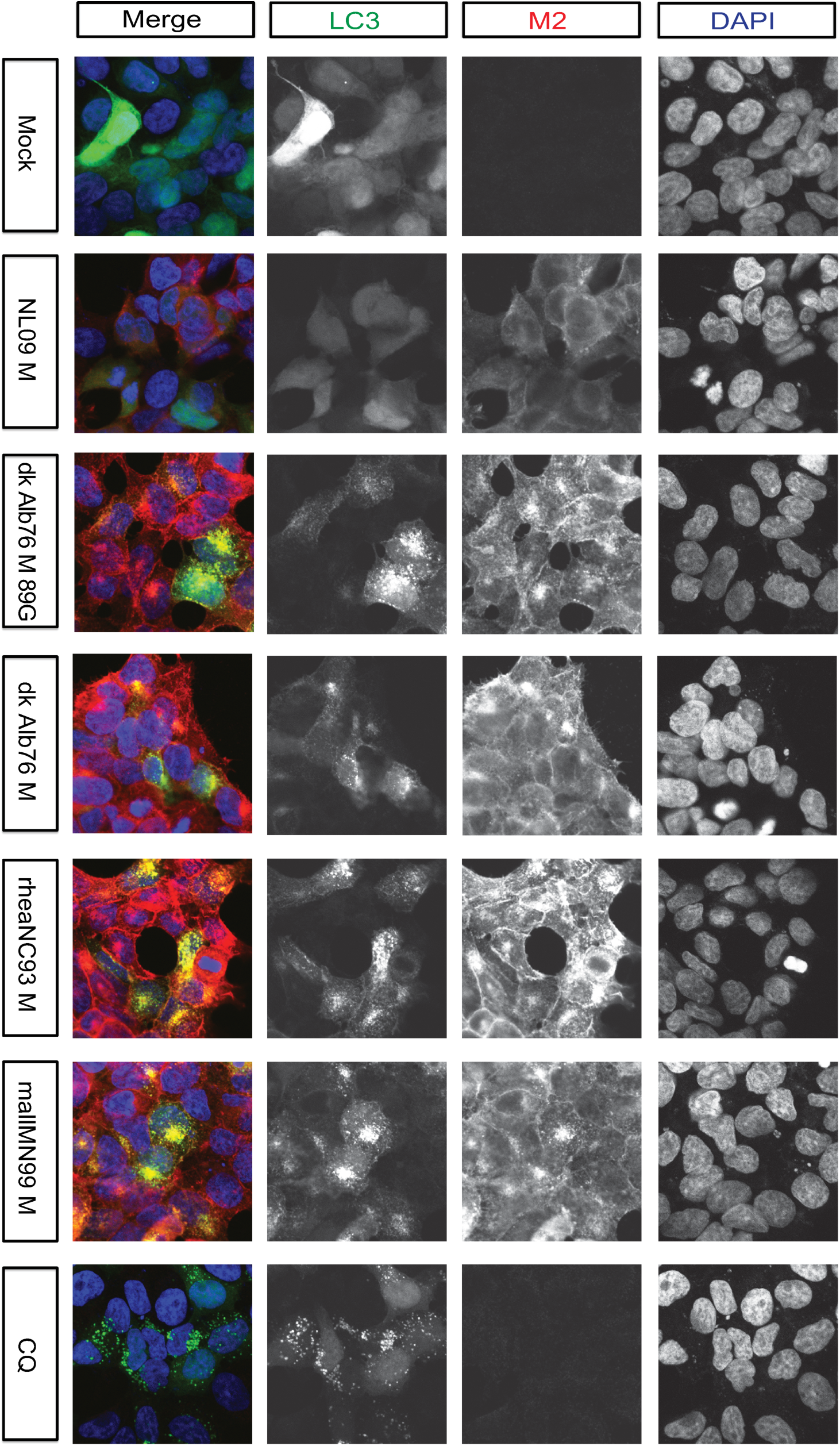
Visualization of LC3 and M2 co-localization. 293T cells were transduced with GFP-LC3 protein and inoculated 24 h later with the indicated IAVs, encoding avian or human-derived M segments, at a MOI of 5 PFU/cell. Cells were fixed 8 h later and stained with anti-M2 (Mab E10; red) and DAPI (blue) followed by imaging with confocal microscopy. CQ: chloroquine. Examples of optical sections are shown, either as merged 3-color images, or the red, green, and blue channels alone (in grey scale). Images are at 63x optical and 3x digital magnification (189x total magnification). Brightness was adjusted for optimal clarity, with all images treated equally.

As an additional test designed to differentiate autophagy induction from autophagy block in cells infected with IAV carrying avian M segments, we used chloroquine treatment followed by Western immunoblotting for LC3B I and LC3B II. We hypothesized that, if autophagic flux is blocked in cells infected with PR8 avian M viruses, then treatment with chloroquine would not lead to a further increase in LC3B II levels. Conversely, if virus infection of these cells induces autophagy, then blocking their turnover with chloroquine would result in heightened LC3B II over mock-infected, chloroquine-treated cells. As shown in **Suppl. Figure 8A and 8C**, we observed a significant difference in the activation of LC3B II following infection with human- or avian M-encoding viruses. This difference in LC3B II activation is lost in the presence of chloroquine. Additionally, as we observed no change in LC3B II levels upon treatment of virus-infected cells with chloroquine, relative to chloroquine-treated, mock-infected cells, we conclude that the high levels of IAV M2 protein expressed from avian M segments, but not human M segments, precipitate a block in the autophagy pathway such that chloroquine treatment cannot bring about further accumulation of autophagosomes.

### Ion channel activity of IAV M2 contributes to LC3B II accumulation in A549 cells

To evaluate the contribution of IAV M2 ion channel activity to the observed stalling of autophagy in PR8 avian M virus-infected cells, we inhibited channel activity with amantadine [61]. Amantadine was added at 1 hpi and the level of LC3B lipidation was assessed at 8 hpi in PR8 NL09 M and PR8 avian M virus-infected A549 cells. The timing of amantadine addition was designed to avoid disruption of the early function of M2 in delivery of the viral genome to the cytoplasm, but allow inhibition of M2 activity in autophagosomes.

As noted above (**Figure 5; Suppl. Figure 4**), Western immunoblot analysis of LC3B revealed decreased LC3B I and increased LC3B II upon infection of A549, 293T or MDCK cells, with avian or human M-encoding IAV, with the effect being stronger for viruses possessing avian M segments. Interestingly, addition of amantadine reversed the intracellular accumulation of LC3B II in both human and avian M-infected A549 cells (**Figure 10**). Quantitation of M segment-encoded proteins by Western immunoblot analysis showed no difference in the expression levels of M1 or M2 protein upon treatment with amantadine (**Figure 10 A, 10B, 10C, 10D**), but revealed a statistically significant increase in LC3B I (**Figure 10E**) and decrease in LC3B II (**Figure 10F**) to levels comparable to mock-infected cells.

**Figure 10.**
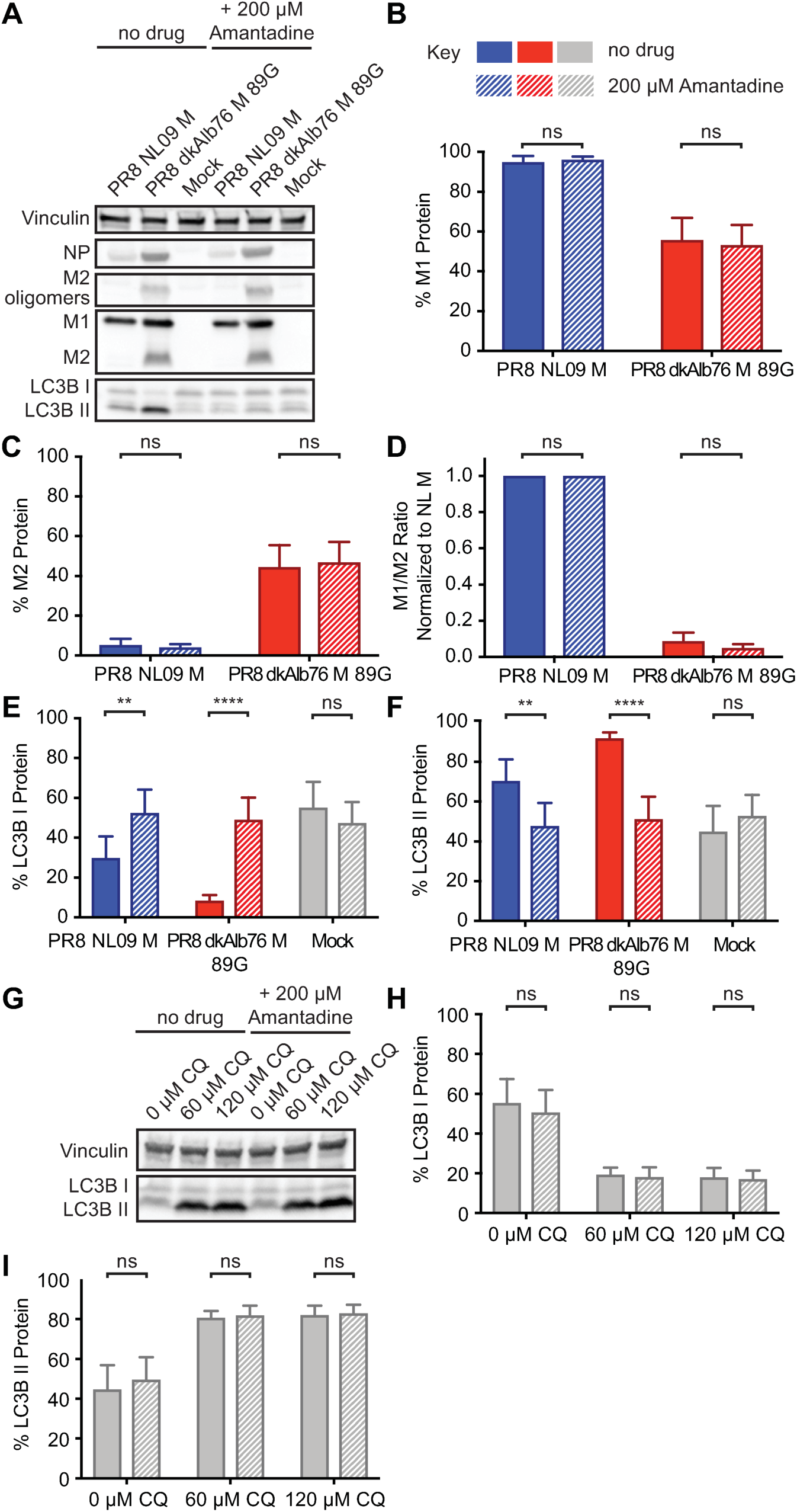
Inhibition of M2 ion channel results in loss of activation of LC3B. A549 cells were inoculated at an MOI of 5 PFU/cell with PR8 viruses encoding avian or human-derived M segments and incubated in the presence or absence of 200 μM amantadine from 1 hpi. Cells were lysed at 8 hpi. **(A)** Representative Western immunoblot. Vinculin expression was measured to allow normalization of viral protein levels. NP expression was measured to assess viral replication. Levels of M1 and M2 protein were assessed using an antibody (Mab E10) to a common epitope of M1 and M2 proteins, allowing relative expression to be assessed. Levels of LC3B I and II were assessed using an antibody that detects both the precursor and activated forms of LC3B protein. **(B)** M1 protein and **(C)** M2 protein were normalized to vinculin and displayed as a percentage of total M gene-expressed protein. **(D)** The ratio of M1:M2 protein expression. **(E)** LC3B I and **(F)** LC3B II were normalized to vinculin and displayed as a percentage of total LC3B protein. **(G)** As a control to ensure the specificity of amantadine, we incubated chloroquine-treated A549 cells in the presence or absence of 200 μM amantadine for 8 h. **(H)** LC3B I and **(I)** LC3B II were normalized to vinculin and displayed as a percentage of total LC3B protein. Graphs in **B-F, H-I** show the means with SD from three independent experiments. For each experiment, two replicate Western immunoblots were performed and quantitated. Statistical significance was assessed using ordinary one-way ANOVA.

To ensure that the amantadine was working to prevent LC3B II accumulation through inhibition of M2 ion channel activity, we assessed the impact of amantadine treatment on the inhibition of phagosome-lysosome fusion by chloroquine. Amantadine had no impact on LC3B II accumulation in the absence of M2 protein (**Figure 10G-I**), indicating that the mechanism of amantadine block of autophagosome accumulation was specifically mediated through interference with M2 function.

As a complementary approach to monitor the effects of amantadine, we again used overexpressed LC3B-GFP fusion protein in infected 293T cells at 12 hpi. Using confocal microscopy, we tracked LC3B-GFP localization in the presence and absence of the ion channel blocker. In line with the Western immunoblot analyses obtained in A549 cells, we saw that amantadine reduced the accumulation of LC3B-GFP within cytoplasmic puncta and at the plasma membrane of infected cells (**Figure 11**). Of note, amantadine treatment did not alter levels or localization of M2 protein and, moreover, had no detectable effect on LC3B-GFP appearance in chloroquine-treated cells (**Figure 11**).

**Figure 11.**
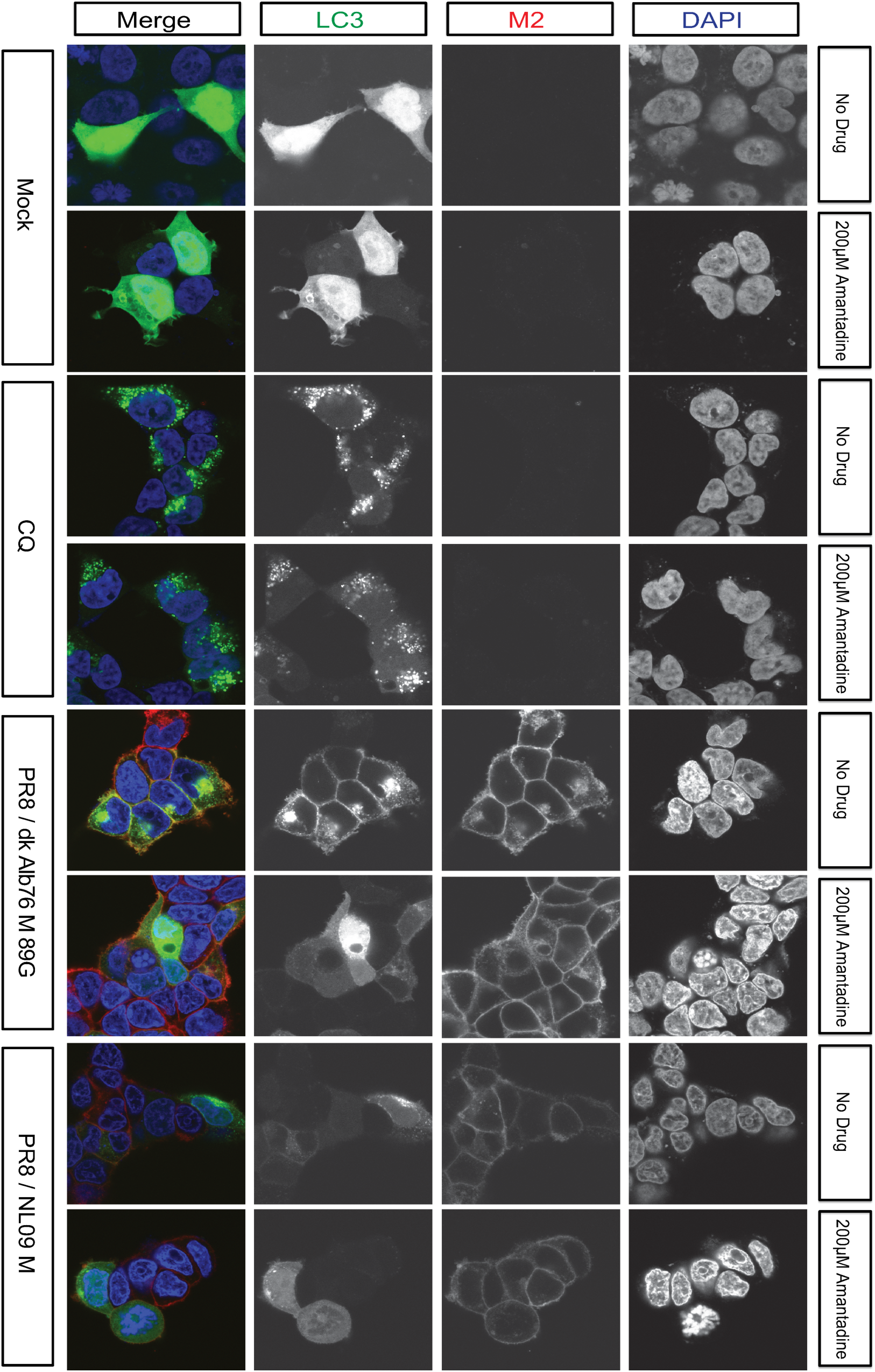
Inhibition of proton channel activity relieves M2-induced accumulation of LC3B positive vesicles. 293T cells transduced with LC3-GFP were inoculated at a MOI of 5 PFU/cell with PR8 viruses possessing M segments from a human or an avian strain and incubated in the presence or absence of 200 μM amantadine from 1 hpi. Cells were fixed at 12 hpi and stained with anti-M2 (Mab E10; red) and DAPI (blue) followed by imaging with confocal microscopy. Examples of optical sections are shown, either as merged 3-color images or the red, green, and blue channels alone (in grey scale). 3x zoom of a 63x magnification is shown. Brightness was adjusted for optimal clarity, with all images treated equally. CQ: chloroquine.

Overall, our results indicate that the intracellular ion channel activity of IAV M2 protein is necessary to mediate the observed block in autophagy, and that addition of amantadine can reverse the autophagic phenotypes induced by both human and avian M-encoding IAVs.

### Improved viral growth in the presence of amantadine implicates autophagy block in viral growth restriction

To test for a causal link between the observed block in autophagy and the restriction of growth seen for PR8 avian M-encoding viruses in mammalian cells, we monitored the effect on viral growth of relieving the autophagy block with amantadine treatment. Specifically, we evaluated single-cycle growth in A549 cells using PR8 NL09 M and PR8 dkAlb76 M 89G viruses. Amantadine was again added at 1 hpi to avoid disruption of the earliest steps of the viral life cycle. While amantadine had no effect on growth of the PR8 NL09 M virus (P=0.271), a modest but consistent increase in the titer of PR8 dkAlb76 M 89G virus was noted throughout the time-course (P=0.0007) (**Figure 12**). This result indicates that M2 ion channel-induced disruption of vesicular homeostasis contributes to the host restriction conferred by avian IAV M segments.

**Figure 12.**
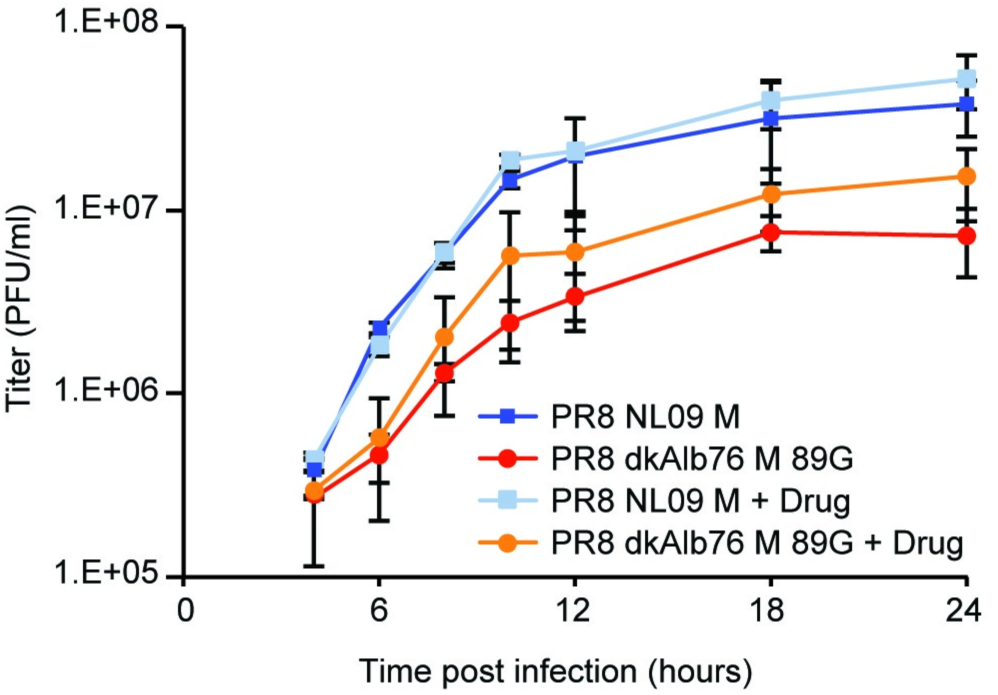
Amantadine improves growth of avian M segment-encoding virus in A549 cells. PR8-based NL09 M or avian M-encoding viruses were inoculated at an MOI of 5 PFU/cell onto A549 cells. At 1hpi, cells were treated with 200μM amantadine, and incubated at 37°C for 24 h in the continuous presence of amantadine. Virus released into supernatant was collected at the indicated time points, and virus growth was measured by plaque titration. The pH1N1 M segment conferred more rapid kinetics and higher peak titers of growth than the avian-origin M segment, however amantadine had no impact on the growth of the pH1N1 M encoding virus. In contrast, growth of the avian M-encoding virus was improved in the presence of drug (P= 0.0007). Single-cycle growth was assessed in four independent experiments, with three technical sample replicates per experiment. Graphs show the means with SD for the four experiments. Statistical significance was determined using repeated measures, two-way, multiple comparisons ANOVA on log-transformed values, with Sidak’s correction applied.

### Viral growth restriction and autophagy block are linked to M2 expression levels, not amino acid sequence

To determine whether the observed effects of avian-origin M segments on the viral life cycle and the autophagy pathway are attributable to avian-adapted amino acid sequences or rather to the overexpression of the viral proton channel, we generated a series of chimeric M segments in which the coding changes found in the NL09 M segment were transferred to an avian-adapted RNA background, and vice versa. We introduced all 16 coding changes into M1 and M2 simultaneously, or introduced only the nine or seven changes into M1 or M2, respectively. The genotypes of this panel of six chimeric viruses are outlined in **Table 2** and Suppl. Figure 9.

**Table 2.** Genotypes of avian-human chimeric M segments.

|  | Nucleic acid background | M1 amino acid sequence | M2 amino acid sequence |
| --- | --- | --- | --- |
| NL M 16mut | NL09 M | Avian consensus M1 | Avian consensus M2 |
| NL M 9mut | NL09 M | Avian consensus M1 | NL09 M2 |
| NL M 7mut | NL09 M | NL09 M1 | Avian consensus M2 |
| Av M 16mut | Dk Alb 76 M 89G | NL09 M1 | NL09 M2 |
| Av M 9mut | Dk Alb 76 M 89G | NL09 M1 | Avian consensus M2 |
| Av M 7mut | Dk Alb 76 M 89G | Avian consensus M1 | NL09 M2 |

Western immunoblot analysis of M1 and M2 proteins expressed from these six chimeric M segments in infected A549 cells is shown in **Figure 13A**. Among the viruses with the NL09 M nucleic acid background, the PR8 NL M 16mut and PR8 NL M 9mut viruses both showed increased levels of M2 relative to the PR8 NL09 M virus control, indicating that all or some of the coding changes introduced into the M1 ORF impact regulation of M2 gene expression. The PR8 NL M 7mut virus showed similar M1 and M2 expression levels as the PR8 NL09 M control, however. This virus therefore expresses the avian consensus M2 protein at low levels. All viruses with the nucleic acid background of the avian M segment showed relatively high levels of M2, similar to those seen for the PR8 dkAlb76 M 89G virus control. Thus, in the avian nucleic acid background, we were able to fully separate M2 expression level from M2 and M1 amino acid sequence.

**Figure 13.**
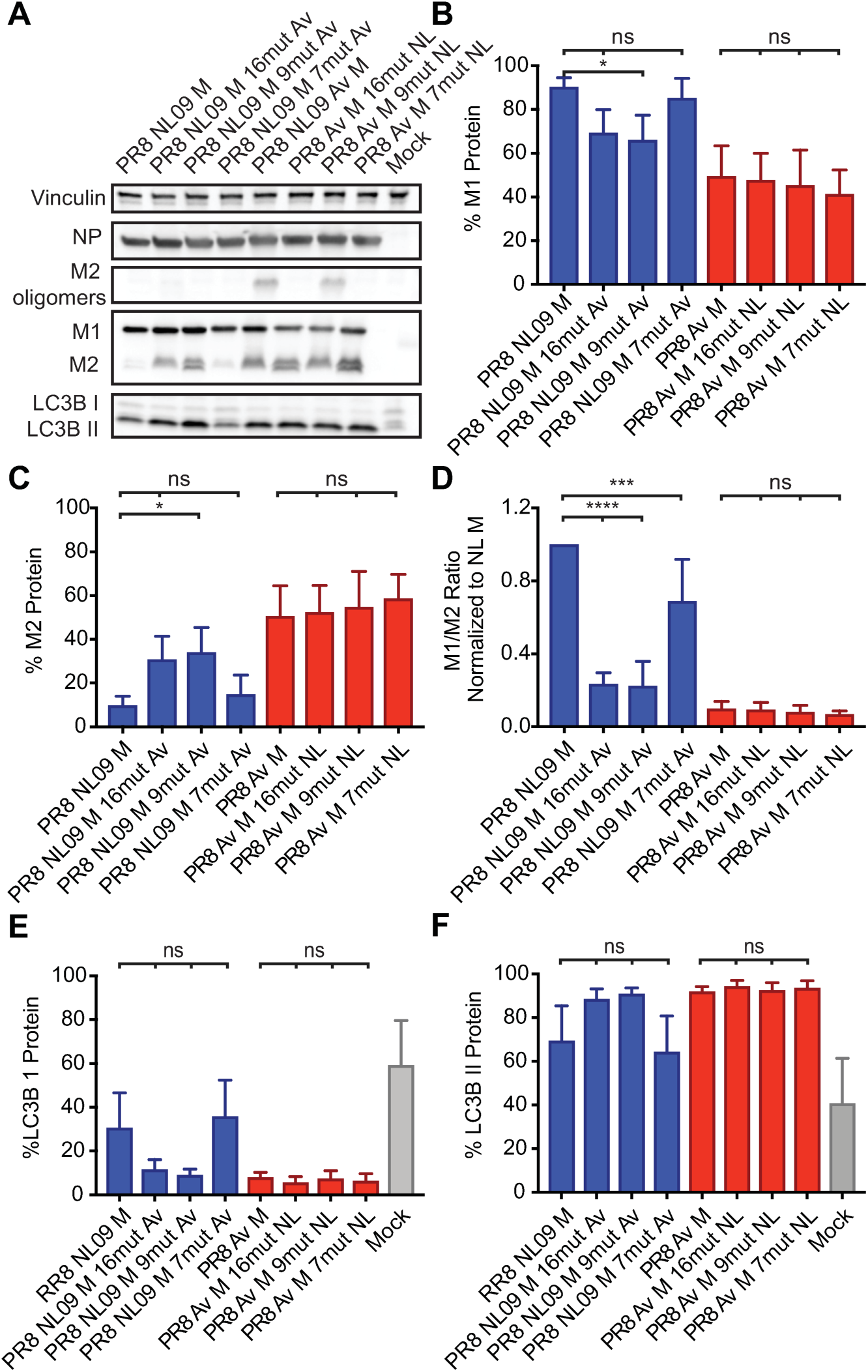
Activation of LC3B II correlates with high expression of M2 protein in A549 cells regardless of M2 amino acid composition. PR8 NL09 M virus and PR8 avian M virus, along with six chimeric variant PR8-based viruses, were inoculated at a MOI of 5 PFU/cell onto A549 cells. Cells were incubated for 8 h, then lysed. **(A)** Western immunoblot analysis of virus-infected A549 cells. Vinculin was measured to allow normalization of viral protein levels. NP was measured to assess viral replication. Levels of M1 and M2 protein were assessed using an antibody (Mab E10) to a common epitope at the amino terminus of M1 and M2 proteins, allowing their direct comparison. Levels of LC3B I and II were assessed using an antibody that detects both the precursor and activated forms of LC3B protein. **(B)** M1 protein and **(C)** M2 protein were normalized to vinculin and displayed as a percentage of total protein expressed from the M gene. **(D)** The ratio of M1:M2 protein expression. **(E)** LC3B I protein and **(F)** LC3B II protein were normalized to vinculin and displayed as a percentage of total LC3B protein. Graphs in **B-F** show the means with SD from three independent experiments. For each experiment, two replicate Western immunoblots were performed and quantitated. Statistical significance was assessed using ordinary one-way ANOVA.

Single-cycle growth analysis of the chimeric viruses in A549 cells revealed a strong correlation between M2 expression level and viral yields (**Figure 14A, B and C**). Notably, all viruses with the dkAlb76 M 89G-derived nucleic acid background, which express high levels of M2 protein, grew to approximately 10-fold lower titers than the PR8 NL09 M virus, despite expression of NL09 M1 and/or M2 proteins. In addition, the PR8 NL M 7mut virus, which expresses the human M1 and avian M2 proteins at levels typical of human-adapted strains, showed equivalent growth to that seen with the PR8 NL09 M virus, indicating that amino acid changes in M2 are not required for efficient replication in human cells. Notably, the PR8 Av M 9mut virus, which encodes identical M1 and M2 proteins as PR8 NL M 7mut virus (**Suppl. Figure 9**), exhibits approximately 10-fold reduced viral replication relative to PR8 NL M 7mut virus (P=0.0003) (**Figure 14B and C**). This result confirms that overexpression of M2 protein is a dominant negative trait that is deleterious to virus growth.

**Figure 14.**
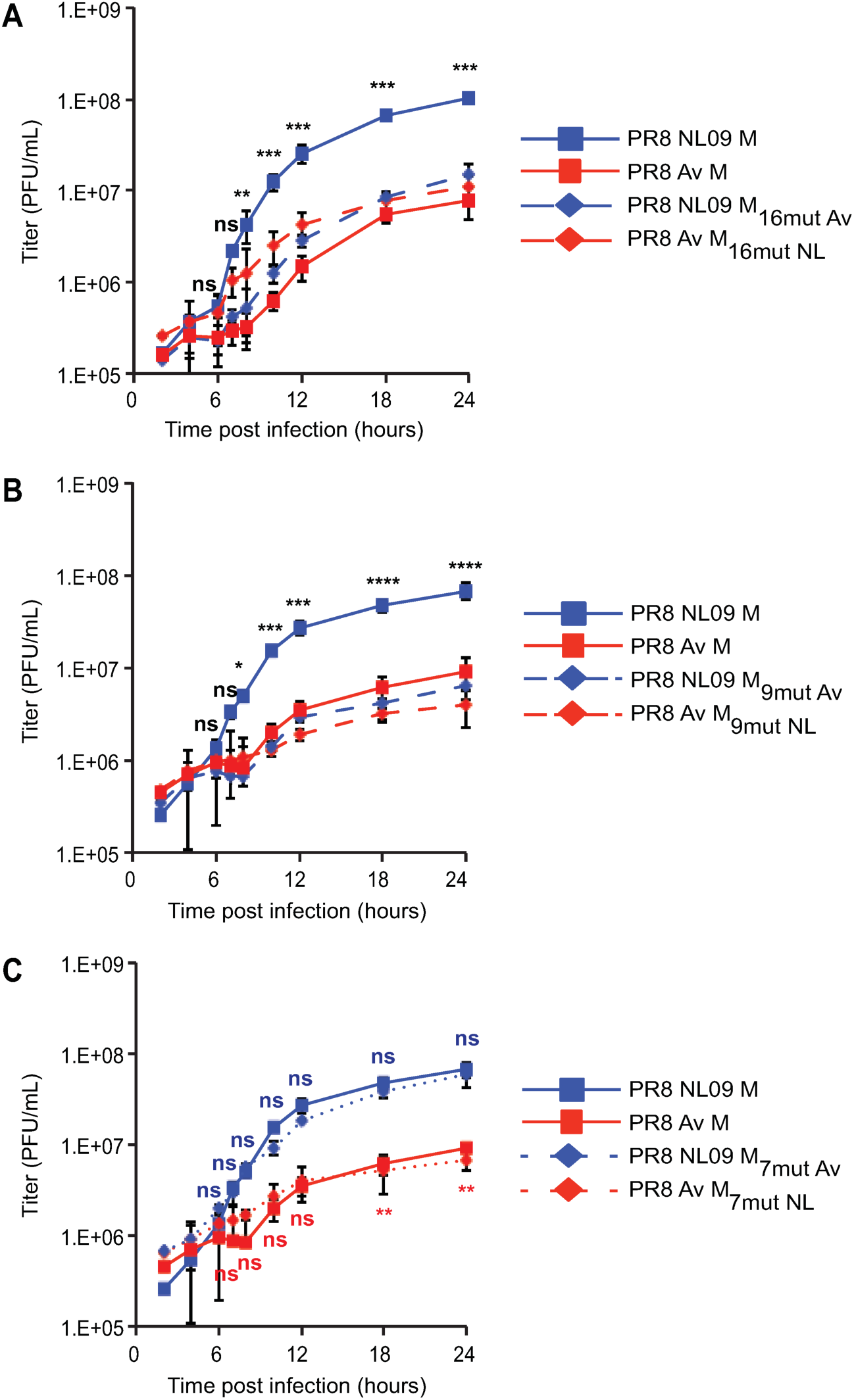
pH1N1 M amino acid sequences are not sufficient to confer high growth phenotype in A549 cells. PR8 NL09 M virus and PR8 avian M virus, along with six PR8-based viruses carrying avian-human chimeric M segments, were inoculated at a MOI of 5 PFU/cell onto A549 cells. Virus growth was measured by plaque titration. The effects on viral growth of amino acid changes in **(A)** both M1 and M2, **(B)** M1 only and **(C)** M2 only are shown. Single-cycle growth was assessed in at least three independent experiments, with three technical sample replicates per experiment. Graphed data show the means with SD for at least three experiments. Statistical significance was determined using repeated measures, two-way, multiple ANOVA on log-transformed values, with Bonferroni correction applied to account for comparison of a limited no of means. Experiments involving PR8 NL09 9mut or 7mut viruses as well as PR8 Av M 9mut or 7mut viruses were performed at the same time, but are displayed on two separate graphs with the same PR8 NL09 M and PR8 avian M control virus data, to improve clarity of presentation.

Examination of LC3B lipidation by Western immunoblot analysis also revealed a clear correlation between M2 expression levels and those of LC3B II within infected cells (**Figure 13A**). Importantly, overexpression of either human or avian M2 during infection boosted LC3B II levels (compare lanes 2, 3, 5, 6, 7 and 8 in **Figure 13A**), while expression of avian M2 at levels comparable to those of PR8 NL09 M virus gave low LC3B II levels (compare lanes 1 and 4 in **Figure 13A**). Confocal images of 293T cells transduced with LC3B-GFP and infected with the chimeric viruses were consistent with these findings. As seen for wild-type avian M segments, high M2 expression again led to the accumulation of LC3B-GFP in perinuclear puncta and at the plasma membrane (**Suppl. Figure 10**). Thus, as with viral growth, the accumulation of autophagosomes within infected cells appears to be triggered by high levels of M2, regardless of amino acid sequence.

To verify that the levels of M1 and M2 protein exhibited by the chimeric M segment-encoding viruses are modulated at the level of mRNA expression, we conducted RT primer extension assays. These data confirmed that differences in the levels of segment 7 mRNA expression (**Suppl. Figure 11**) mirrored the observed differences in M1 and M2 protein expression levels. Overall, these data support the premise that splicing signals are at least partly encoded in non-synonymous nucleotide changes residing in the M1 ORF.

### Chimeric M segment expressing NL09-like levels of NL09 M1 and avian M2 supports robust growth *in vivo*

To evaluate whether correcting expression of the avian M2 protein to levels seen for viruses with human-adapted M segments yielded a well-adapted phenotype in a mammalian animal model, we evaluated growth and transmission of PR8 NL09 M 7mut virus in guinea pigs. Titration of nasal wash samples revealed that the PR8 NL09 M 7mut virus (which expresses avian M2 at low levels) had a similar growth phenotype to that of PR8 NL09 M virus: growth kinetics and area under the curve values were not significantly different (**Figure 15; Suppl. Figure 12**). In contrast, analysis of nasal wash samples collected from contact animals indicated that only one of eight inoculated guinea pigs transmitted the PR8 NL09 M 7mut virus to her cage mate. This level of transmission is significantly decreased relative to the PR8 NL09 M virus (P=0.0339) (**Figure 3B**). Taken together, these results indicate that, when expressed at levels typical for a human-adapted M2 protein, an avian M2 supports efficient replication but inhibits transmission in a mammalian host.

**Figure 15.**
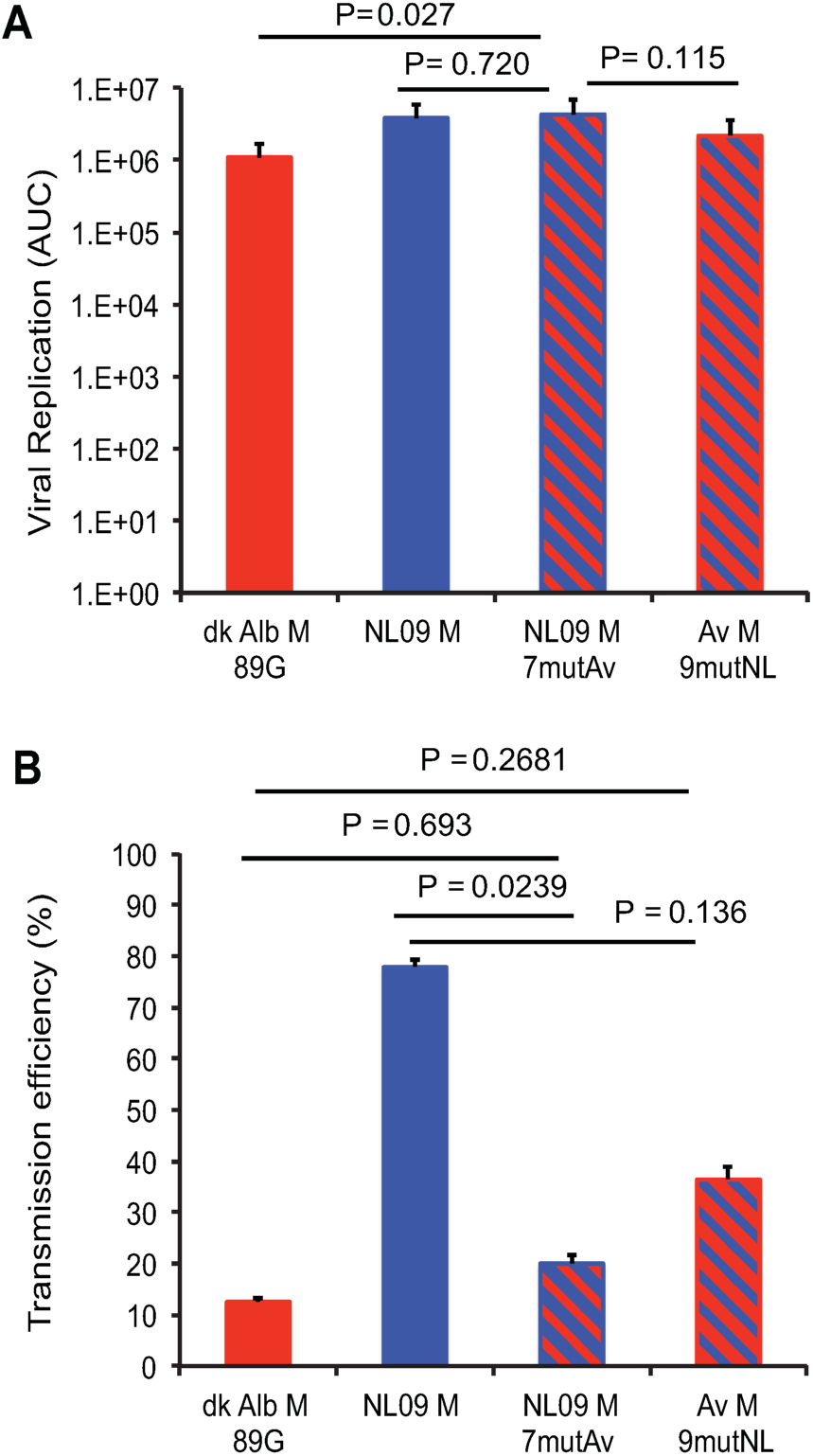
Reduced expression of avian M2 improves replication efficiency *in vivo* but is not sufficient for efficient transmission in guinea pigs. Groups of twelve guinea pigs were inoculated with 10 PFU of PR8-based viruses carrying avian M, NL09 M, chimeric NL09 M 7mut M, or chimeric avian M 9mut segments, as indicated. **(A)** Virus replication in nasal washings was measured by plaque titration at days 2, 4, 6, and 8 post-infection and the total area under the curve was calculated. The replication differences between viruses with NL09 M and NL09 M 7mutAv segments were not significant (P=0.720). Differences between viruses encoding the NL09 M 7mut segment and the avian M 9mut segment trended towards, but were not statistically significant (P=0.115). (**B**) Transmission to naïve cage mates that were exposed at 24 hpi. Neither chimeric M segment conferred efficient transmission (7/9 transmissions; 78%). Statistical significance of replication differences was determined using a two-tailed Student’s t-test. Statistical significance of transmission differences was determined using an unpaired two-tailed Student’s t-test. NL M and avian M data are identical to data presented in Figure 3 and are reproduced here for clarity.

### Lab-adapted IAV, but not recent human isolates, express abundant M2 and block autophagy in human cells

The M2-induced block in autophagic flux reported previously was based on multiple IAVs that were originally isolated from human hosts but passaged extensively in laboratory substrates [46–48]. To assess whether these prior data were consistent with our own, we compared M2 expression levels exhibited by one of the lab-adapted strains used, PR8. Our results show that the relative expression levels of M1 and M2 in PR8-infected 293T cells are comparable to those seen in cells infected with avian M-encoding viruses (**Suppl. Figure 13**). These results suggest that prior reports of M2-induced block in autophagy reflect a feature of mal-adaptation to the host cell.

## Discussion

Our data indicate that dysregulation of gene expression from the IAV M segment contributes to the limited replicative capacity of avian IAVs in mammalian hosts. When transcribed within mammalian cells, avian IAV M segments gave rise to abundant mRNA_10_ and correspondingly high levels of the encoded M2 proton channel. Comparison to matched virus-host systems indicated that this M2 expression profile was an aberrant feature of avian IAV replication in mammalian cells. When present at high levels, M2 had marked effects on the infected cell, blocking the turnover of autophagic vesicles and redirecting LC3B II, the activated form of a critical autophagy mediator, to the plasma membrane. These effects were found to rely on the proton channel activity of M2 and occurred when either the avian or human M2 protein was overexpressed in infected cells. Importantly, the reduction in viral growth conferred by avian M segments in mammalian systems could be attributed at least in part to disruption of vesicular homeostasis and was fully attributed to the altered expression levels of M1 and M2. Thus, our data identify the regulation of viral gene expression as a novel host-dependent feature of the IAV lifecycle that contributes to the host range restriction of this virus (**Figure 16**).

**Figure 16.**
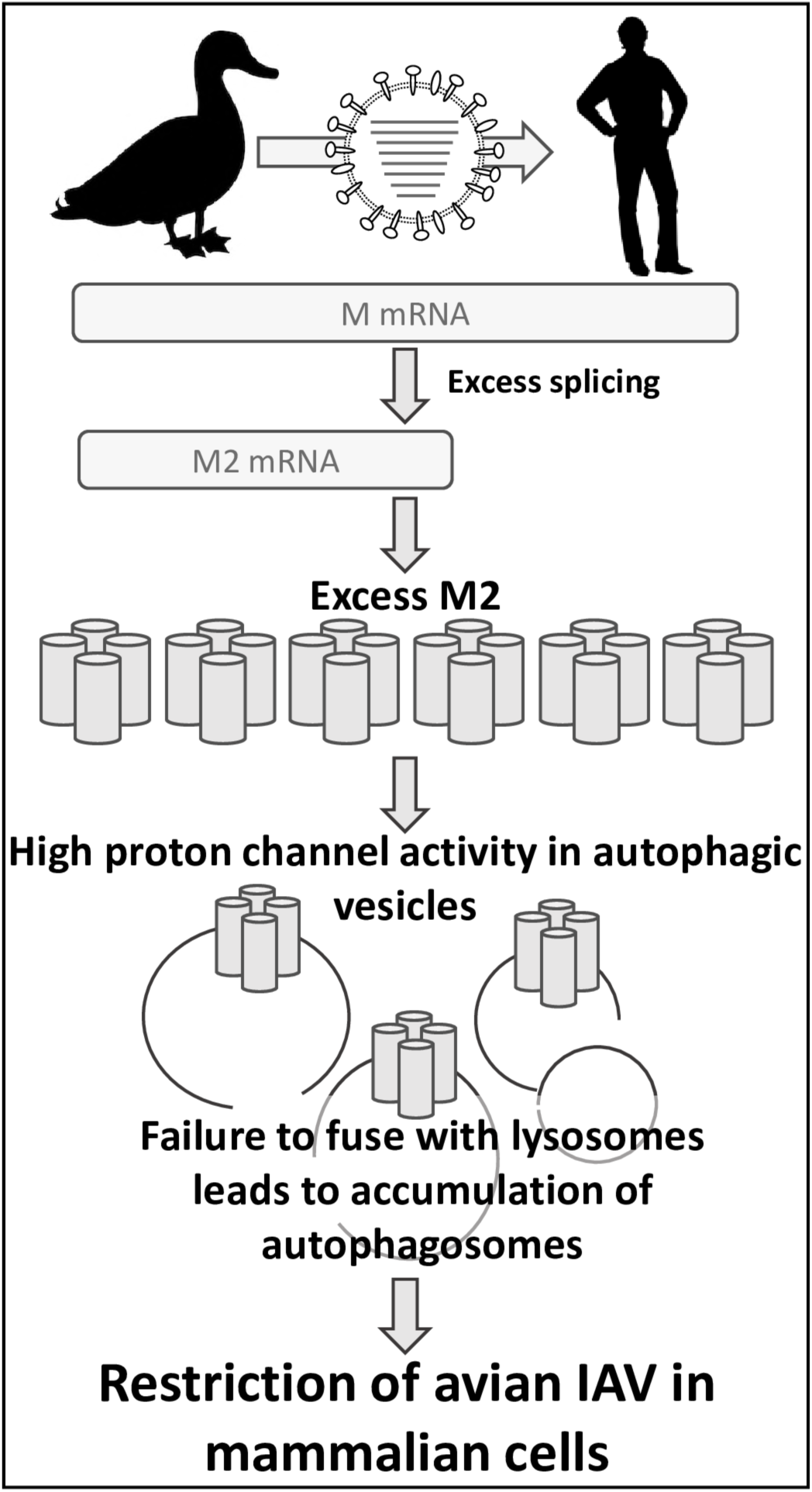
Regulation of M segment gene expression contributes to host range restriction of IAV. When an avian IAV M segment is transcribed within mammalian cells, excessive splicing of mRNA_7_ results in aberrant overexpression of the M2 proton channel. When highly expressed, this channel accumulates in autophagic vesicles. Abundant proton channel activity in the membranes of autophagosomes blocks their turnover by preventing fusion with lysosomes. This stalling of a critical cellular housekeeping function in turn reduces the efficiency of viral replication, contributing to the species barrier that limits the zoonotic potential of avian IAVs.

Quantification of IAV encoded mRNAs within infected cells indicated that overexpression of M2 from avian IAV M segments stems from over-production of the corresponding mRNA. M2 is encoded by the mRNA_10_ transcript, which is generated through splicing of the M1 message, mRNA_7_[18,56]. Thus, regulation of M2 gene expression is intimately linked to the cellular splicing machinery. Our data indicate that this is a finely balanced interaction between IAV and its host cell, which is susceptible to disruption upon transfer to new species. In this regard, it is notable that the M segment splice donor and acceptor sites are identical between the human and avian IAVs examined. M segment sequences outside of these canonical splice sites have, however, been shown to modulate splicing efficiency [62–65]. Interestingly, mRNA_7_ association with mammalian SF2/ASF [64], and with mammalian hnRNPK and NS1-binding protein [65], have been shown to alter the efficiency of splicing. Nucleotide differences in these binding regions exist between human- and avian-adapted M segments. An interesting area of further investigation will be to test whether there are differences in the binding of human and avian mRNA_7_ molecules to mammalian SF2/ASF, hnRNPK, and NS1-binding proteins.

The gene segments of IAVs circulating in humans are ultimately derived from the avian IAV gene pool. Thus, the contrasting phenotypes conferred by avian- and human-adapted M segments in mammalian cells suggest that a resetting of M segment gene expression to reduce M2 levels has been positively selected following emergence of avian IAVs into mammalian populations. Indeed, our data reveal that this adaptation has occurred in at least two independent incidences: i) within the Eurasian avian-like swine lineage M, that emerged from birds in the late 1970s and contributed the M segment to the pH1N1 virus prior to 2009; and ii) within the human seasonal lineage derived from the 1918 pandemic, which contributed its M segment to the H3N2 lineage represented by Pan99 virus. The timing of the adaptive changes is unclear in each case and cannot be inferred from our data. Importantly, M2 expression in human IAV mimics that of avian IAV in avian cells. Thus, restriction of splicing efficiency to achieve low M2 levels appears to be an important feature of IAV host adaptation. Our data furthermore suggest that the negative impact of an autophagy block on viral growth constitutes an important pressure driving this selection.

To date, most publications related to autophagy in IAV-infected cells have used the PR8 or WSN lab-adapted strains, the X31 variant of PR8, or avian IAVs [46,47,66,67]. We found that PR8 virus expresses levels of M2 in mammalian cells that are comparable with avian IAVs. This phenotype may be a result of adaptation to growth in chicken eggs. Owing to the use of viruses carrying lab-adapted M segments, the current literature suggests that IAV infection routinely induces a block in autophagy [46,47,66,67]. We also see evidence of such a block when using viruses that express high levels of M2, but we propose that this phenotype is a symptom of poor adaptation to the mammalian host cell. Analysis of viruses carrying human M segments in mammalian cells suggests that low levels of M2 and activation, but not blocking, of autophagy are the norm in IAV-infected cells.

While the previously reported block in autophagy induced by IAV has generally been interpreted as a viral defense against an antiviral mechanism, a pro-viral role for autophagy is not without precedent. In Dengue virus infection, autophagy has been reported to support viral replication. Specifically, autophagic turnover of lipid droplets in infected cells is thought to provide energy needed for viral growth and to supply phospholipids important during virion assembly [68–70]. For IAV, the availability of activated LC3B protein was reported to be important for viral morphogenesis [46]. Specifically, M2 was implicated in recruitment of LC3B II to the plasma membrane, and disruption of this recruitment was found to affect budding and reduce stability of released virions [46]. We also noted strong redistribution of LC3B-GFP to the plasma membrane during infection with viruses that express high levels of M2. This relocalization occurred to a lesser extent in a well-matched IAV-host system (NL09 M in human cells), and may play a functionally important role in this context. A role for LC3B II in morphogenesis is consistent with the activation of autophagy having a pro-viral effect. Conversely, our data indicate that a block in autophagic turnover reduces viral growth, potentially owing to disruption of vesicular trafficking that is normally exploited by the virus [71–73], or due to disruption of cellular metabolic functions on which the virus relies.

By inhibiting the proton channel activity of the avian M2 with amantadine, we found that a functional channel is needed to bring about the observed effects on autophagy. Indeed, M2 has been shown previously to block the fusion of autophagosomes with lysosomes in a proton channel-dependent fashion [66]. Surprisingly, however, amantadine treatment was also found to reduce LC3B II accumulation in PR8 NL09 M-infected cells. The pH1N1 M2 protein carries the S31N mutation, which confers resistance to amantadine. Our data suggest that this resistance is not effective at a relatively high concentration of 200 μM amantadine, at least in the context of intracellular viral infection.

The mutagenesis of the M segment that we carried out with the goal of separating the effects of M2 expression level from M2 amino acid sequence revealed an important constraint on M1 coding capacity. Namely, the human / avian amino acid polymorphisms introduced into the M1 ORF were found to impact regulation of M2 gene expression. Some or all of these non-synonymous changes to M1 likely modulate the efficiency of mRNA_7_ splicing. With this new insight in mind, a fresh look at viral phenotypes previously attributed to M1 amino acid variations is warranted. Effects of M1 amino acid sequence on virion morphology [25,31,32,74,75], for example, may in fact be directly or indirectly mediated through changes in intracellular M2 expression level.

Notably, while an avian M2 protein was sufficient to support robust viral replication in mammalian systems when expressed at low levels, this sufficiency did not extend to transmission. Virus expressing the avian M2 from a human M segment transmitted poorly, indicating that the IAV M2 carries viral determinants of transmission and that these determinants are not well conserved between avian and human strains. This novel observation also supports the notion that high replicative capacity of a virus in a given host species is not necessarily sufficient for transmission between individuals. In this context, it will be interesting to evaluate the impact of avian M2 amino acid signatures on virion stability in the environment.

In summary, our data suggest that adaptive change in the M segment is needed to maintain low expression of M2 in mammalian cells and that, in the absence of such adaptive changes, excess M2 limits viral growth by blocking the autophagy pathway. These findings establish regulation of viral gene expression as a feature of the virus-host interface critical for IAV host range determination and therefore emergence into novel host populations.

## Materials and Methods

### Ethics Statement

The Institutional Animal Care and Use Committee (IACUC) of Emory University approved the study protocol under approval number PROTO201700595.

### Cells

Madin-Darby Canine Kidney (MDCK) cells (a kind gift of Peter Palese, Icahn School of Medicine at Mount Sinai) were maintained in minimal essential medium (Gibco) supplemented with 10% fetal bovine serum (FBS) and penicillin-streptomycin. A549 and 293T cells were obtained from the ATCC and maintained in Dulbecco’s minimal essential medium (DMEM; Gibco) supplemented with 10% FBS and penicillin-streptomycin. DF-1 cells were obtained from ATCC and maintained in DMEM plus 5% FBS and penicillin-streptomycin. QT-6 cells were obtained from ATCC and maintained in F-12K medium plus 5% FBS, 10% tryptose phosphate and penicillin-streptomycin. All cells were incubated at 37°C with 5% CO_2_.

### Viruses

All viruses used in this work were generated using reverse genetics techniques [76]. The rescue system for PR8 was a gift of Peter Palese (Icahn School of Medicine at Mount Sinai). The rescue system for A/Panama/2007/99 (H3N2) [Pan/99] virus was initially described in [7]; and that for A/Netherlands/602/2009 (H1N1) [NL09] virus was a gift of Ron Fouchier (Erasmus Medical Center) [77]. The NL09 virus was propagated in MDCK cells for two passages. All other virus stocks were generated in 10-11 day old embryonated chicken eggs.

### Generation of chimeric M segments

Non-synonymous nucleotide changes differentiating NL09 and avian consensus M segments were introduced in a reciprocal fashion into i) M1 and M2 to generate 16 mut viruses, ii) M1-only to generate 9 mut viruses, and iii) M2-only to generate 7 mut viruses. The modified viral cDNAs were synthesized by Genewiz and subcloned into the pDZ reverse genetic vector [78].

### Sequence alignments and generation of consensus sequences

Alignment of the FASTA sequences downloaded from Genbank (or GISAID) databases was carried out using DNASTAR Lasergene 13 MegAlign software running the Clustal W phylogenetic alignment algorithm.

To obtain a consensus nucleotide sequence and to assess the degree of variability in the M segment of H1N1 subtype avian IAVs, we searched the Genbank nucleotide database for full-length M segment sequences derived from avian H1N1 subtype IAV collected between 1976 and 2008 from any temperate Northern hemisphere location. We obtained 262 sequences, which included sequences with partial 3’ and 5’ UTRs, and manually curated the sequences to remove swine and human lineage derived H1N1 M segments, retaining 247 sequences, and trim UTR sequences from the retained segments. Alignment of the sequences by Clustal W provided a working consensus sequence and revealed up to 7% variability at the nucleotide level among the selected avian host-derived sequences.

To assess the degree of amino acid variability in the M segment of avian IAV of any subtype, we searched Genbank for full-length M1 or M2 protein sequences derived from avian H1Nx to H16Nx subtype IAV, collected between 1970 and 2000, from any geographical location. These dates were chosen to avoid selecting sequences representing isolates originating from the repeated H5 subtype HPAI incursions into poultry populations which dominate the Genbank database from the early 2000s onwards, and which would likely bias the subsequent analysis.

For each hemagglutinin subtype (H1Nx to H16Nx), we aligned the available sequences to generate individual consensus M1 and M2 amino acid sequences (**Supplemental Table 1**), and aligned those consensus sequences to the avian H1N1 subtype consensus M1 and M2 sequences, in order to assess amino acid diversity. The matrix proteins of each subtype, with the exception of H9Nx, were 100% conserved. The H9Nx subtype matrix protein shared 97.6% identity to the avian H1N1 sequence. The M2 proteins of each subtype were 100% conserved, excepting H9Nx, H13Nx and H16Nx subtypes, which shared 97.2%, 94.8% and 93.8% identity respectively, to the avian H1N1 consensus sequence. Thus, the amino acid composition of M1 and M2 proteins are highly conserved within wild waterfowl, particularly within isolates that circulate among dabbling ducks and geese. In these host species, both M1 and M2 proteins are at, or close to, 100% identity at the consensus amino acid level.

To identify a consensus nucleotide sequence for the 2009 pH1N1 IAV, and compare it to the avian consensus sequence, we searched Genbank for full-length M segments derived from pH1N1 subtype IAV isolated in North America, collected between the 7^th^ June and the 20^th^ June 2009, which was shortly after emergence of the pH1N1 lineage into the human population. 189 individual nucleotide sequences were obtained, curated, and aligned, to provide a working consensus sequence for the pH1N1 M segment. The pH1N1 M segments possessed >2% variability at the nucleotide level. The A/NL/602/09 (H1N1) (NL09) M segment is a 100% match to the consensus sequence of the 2009 pH1N1 lineage at the nucleotide level, and the M segment from this strain was adopted for use throughout the current study. Interestingly, comparison of the NL09 M segment to the avian consensus sequence revealed 8.3% difference at the nucleotide level, and 9 and 7 amino acid changes in M1 and M2 proteins, respectively (**Table 2**). These data suggest that pH1N1 M segment possesses a nucleotide sequence that is highly divergent from M segments obtained from viruses circulating within the wildfowl reservoir, and which may encode host adaptive changes.

### Evaluation of viral growth in cell culture

To assess growth, subconfluent cell monolayers in 6-well dishes were inoculated at an MOI of 5 PFU/cell (for single-cycle growth assays) or 0.01 PFU/cell (for multi-cycle growth assays) in a 200 μl volume. Following incubation for 1 h at 37°C, inocula were removed, cells washed 3x with PBS and 2 ml viral growth medium (MEM (for MDCK cells), DMEM (for 293T, and DF1 cells), or F-12K (for A549, and QT-6 cells) medium plus 3% bovine serum albumin, penicillin streptomycin, and TPCK-treated trypsin) was added. Infected cells were incubated at 37°C for the remainder of the time-course. At the indicated time points, 120 μl medium was sampled from each dish and 120 μl fresh medium was added to maintain a 2 ml volume. Samples were stored at −80°C and later titered by plaque assay on MDCK cells. Each infection was performed in triplicate and at least three biological replicate infections were carried out on different days. Data are plotted as mean with SD and analyzed by repeated measures ANOVA.

### Evaluation of viral growth in guinea pigs

Female, Hartley strain, guinea pigs weighing 300-350 g were obtained from Charles River Laboratories. Prior to inoculation or nasal lavage, animals were sedated with a mixture of ketamine (30 mg/kg) and xylazine (4 mg/kg). Virus used for inoculation was diluted in PBS to allow intranasal inoculation of guinea pigs with 1×10^1^ to 1×10^3^ PFU in a 300 μl volume. Nasal wash samples were collected as described previously [54], with PBS as the collection fluid. Animals were housed in a Caron 6040 environmental chamber set to 10°C and 20% RH throughout the seven-day period and lids were left off of the cages during this time to ensure environmental control within the cages [79].

### Immunoblotting

A549 or DF-1 cells in 6-well plates were infected with the indicated IAVs and cells were lysed with 2X Laemmli sample buffer (Bio-rad) plus 2% beta-mercaptoethanol at 8 hpi. Samples were boiled at 95 °C for 10 min and resolved on 4-20% gradient SDS-PAGE gels, followed by transfer to nitrocellulose membranes (Bio-rad), and incubation of membranes with 5% non-fat dry milk/TBST blocking buffer. Antibodies used for immunoblotting were: anti-vinculin monoclonal antibody (catalog no. V9131; Sigma-Aldrich) at 1:5000 dilution, IAV nucleoprotein monoclonal antibody (HT-103; catalog no. EMS010, Kerafast) at 1:1000 dilution, IAV matrix protein monoclonal antibody (E10; catalog no. EMS009, Kerafast) at 1:1000 dilution, and LC3B polyclonal antibody (Thermo Fisher Scientific) at 1:1000 dilution. Bands were quantified using ImageLab software (Bio-rad) after normalization to vinculin. At least three biological replicate viral infections were carried out, with two technical replicates of the immunoblotting for each infection. Data shown with error bars representing SD and statistical analysis was carried out using one-way ANOVA with Tukey’s multiple comparisons in GraphPad Prism software.

### RT primer extension assay

A549 or DF-1 cells in 6-well plates were infected with the specified IAVs and total RNA was extracted 8 hpi using a RNeasy Kit (Qiagen). Total RNA was eluted in RNase-free water and stored at −80 °C until needed.

Radiolabeling of primers: 1 μmol of oligo DNA primer was incubated with 10 μCuries of [γ-^32^P]ATP and 10 U of T4 PNK (New England Biolabs) in 70 mM Tris-HCl (pH 7.6) containing 10 mM MgCl_2_ and 5 mM DTT. Radiolabeling reactions were carried out at 37 °C for 1 h and diluted to 30 μL with RNase-free water after termination of the reaction. Radiolabeled primers were stored at –20 °C until needed. 500 ng of total RNA was incubated with 0.45 μL unlabeled 5S rRNA primer, 0.05 μL radiolabeled 5S rRNA primer, 0.25 μL radiolabeled IAV segment 7 vRNA primer, and 0.25 μL radiolabeled IAV segment 7 mRNA primer in a final volume of 5 μL. Primers were annealed to RNA by sequential incubations at 95 °C for 3 min and ice for 3 min. Reactions were pre-warmed at 50 °C for 5 min. Transcription was initiated by the addition of 50 mM Tris-HCl (pH 8.3) buffer containing 75 mM KCl, 3 mM MgCL2, 10 mM DTT, 2mM dNTP, and 50 units of SuperScript III Reverse Transcriptase (Thermo Fisher Scientific). Reactions were incubated at 50 °C for 1 h and terminated by the addition of Gel Loading Buffer II (Thermo Fisher Scientific) and incubation at 95 °C for 10 min. Reactions were resolved on denaturing sequencing polyacrylamide gels (7M urea, 6% acrylamide). Gels were subsequently dried and exposed to a phosphor storage screen overnight. The intensity of bands was analyzed using a Typhoon Trio Imager (GE Healthcare) and quantified with ImageQuant software (GE Healthcare) after normalization to 5S rRNA. At least three biological replicate viral infections were carried out, with two technical replicates of the RT primer extension assay for each infection. Data are shown with error bars representing SD and statistical analysis was carried out using one-way ANOVA with Tukey’s multiple comparisons in GraphPad Prism software.

### Fluorescence microscopy

To allow tracking of LC3B II and viral protein, cells grown on collagen-coated coverslips were transduced with lentivirus expressing LC3B-GFP (BacMamm 2.0; catalog no. P36235, ThermoFisher), as follows.

18h post-transduction, 293T or A549 cells were infected with the indicated IAVs at an MOI of 5 PFU/cell and incubated at 37°C. At 8h or 12 h post-IAV infection, cells were washed once with PBS and fixed with 4% paraformaldehyde. Following permeabilization with 0.1% Triton X-100, cells were incubated overnight at 4°C with anti-M2 monoclonal antibody (E10; catalog no. EMS009, Kerafast). Three washes were performed prior to addition of goat anti-mouse Alexa Fluor-647 secondary antibody (catalog no. A21241, Life Technologies) and incubation for 2 h at room temperature. Antibody dilutions and washes were performed with PBS plus 1% Tween-20. Cells were treated with DAPI, and coverslips were mounted on slides using Vectashield (Vector Labs) mounting medium. Images were obtained with on a Nikon FV1000 confocal microscope at the Emory Integrated Cellular Imaging core facility.

### Drug treatments

Where indicated, chloroquine (Invitrogen) was added to cell culture medium at a final concentration of 60 or 120 μM. Amantadine HCl (catalog no. 1018505, Sigma) was added to cell culture medium at a final concentration of 200 μM. Both drugs were added to infected cells 1 h after IAV inoculation.

### Statistical analyses

#### Statistical analyses were performed in GraphPad Prism

Multi-cycle and single-cycle growth in cell culture was assessed in three independent experiments, with three technical sample replicates per experiment. Overall, statistical significance was determined using repeated measures, two-way, multiple comparisons ANOVA on log-transformed mean values, with Bonferroni correction applied to account for comparison of a limited number of means, as well as by assessing pairwise comparisons of viruses at each time point, except for single-cycle growth in cell culture in the presence of amantadine, which was assessed in four independent experiments, with three technical sample replicates per experiment. Overall statistical significance was determined using repeated measures, two-way, multiple comparisons ANOVA on log-transformed values, with Sidak’s correction applied, as well as by assessing pairwise comparisons of viruses at each time point.

Growth of each virus in the guinea pig model was assessed in three independent experiments with four guinea pigs per experiment. For transmission analysis, each guinea pig was treated as an independent biological sample. Statistical significance in the magnitude of replication was determined by comparing AUC of growth curves using two-tailed Student’s t-test. Statistical significance in transmission efficiency was determined using unpaired two-tailed Student’s t-test. Significance of differences in the kinetics of replication was determined by assessing the interaction of time and virus using repeated measures, two-way ANOVA on mean log-transformed values, with Bonferroni correction applied to account for comparison of a limited number of means.

For experiments presented throughout the manuscript, unless specific values are provided, P values are represented as follows: *< 0.05; **<0.01; ***< 0.001; ****< 0.0001.

## Acknowledgements

We thank Paul Digard for his excellent advice relating to RT Primer Extension assays, and Emory Integrated Cellular Imaging Core Facility for assistance with confocal imaging techniques.

## Supporting Information Legends

**Supplementary Figure 1. Schematic of M segment mRNAs and gene products.**

**(A)** The M segment of influenza virus is template for synthesis of mRNA_7_ (encoding M1), mRNA M_10_ (encoding M2), and mRNA_11_ (which encodes a putative but unconfirmed 10 amino acid peptide from a short open reading frame). (**B)** Pandemic H1N1 influenza virus M1 and M2 proteins differ from the avian consensus sequences by 9 residues in M1 and 7 residues in M2. The M segments differ by 8.3% at the nucleotide level. (**C)** Seasonal H3N2 influenza virus strain A/Panama/2007/99 M1 and M2 proteins differ from the avian consensus sequences by 11 residues in M1 and 14 residues in M2. The M segments differ by 9% at the nucleotide level. (**D)** Seasonal H3N2 influenza virus strain A/Bethesda/55/15 M1 and M2 proteins differ from the avian consensus sequences by 11 residues in M1 and 16 residues in M2. The M segments differ by 9.5% at the nucleotide level.

**Supplementary Figure 2. Human host-derived M segments confer higher growth to PR8-based viruses than avian host-derived M segments in mammalian cells at 37°C.**

PR8-based viruses were inoculated at a MOI of 5 PFU/cell onto human-derived 293T cells **(A)**, or A549 cells **(B)**. Cells were incubated at 37°C for up to 24 h. Virus released into supernatant was collected at the indicated time points, and virus growth was measured by plaque titration. Data obtained from viruses possessing human M segments are represented with blue lines: A/NL/602/09 M **(A,B)**, A/Panama/2007/99 M **(B)**, and A/Bethesda/15 M **(B)**, while data from viruses encoding avian M segments are represented with red lines. In each cell type, the human M segments conferred more rapid kinetics and higher peak titers of growth than any avian-origin M segment. Single-cycle growth was assessed in three independent experiments, with three technical sample replicates per experiment. Graphs show the means with SD for the three experiments. Statistical significance was determined using repeated measures, two-way, multiple ANOVA on log-transformed data, with Bonferroni correction applied as there were a limited no of means to compare.

**Supplementary Figure 3. pH1N1 influenza virus M segment increases kinetics of replication of PR8-based viruses among guinea pigs.**

Groups of four guinea pigs were inoculated with 10 PFU of each avian M-encoding virus, or NL09 M-encoding virus, as indicated. Graphs show individual titers obtained from animals used in three independent experiments. **(A-E)** Virus replication in nasal wash of inoculated animals was measured by plaque titration at days 2, 4, 6, and 8 post-infection and the titers at each time point were plotted (dotted lines). The differences between PR8 NL09 M and each avian M-encoding virus were considered significant. Statistical significance in kinetics of growth was determined by assessing the interaction of time and virus using repeated measures, two-way, multiple comparisons ANOVA on mean values, with Bonferroni correction applied to account for comparison of a limited no of means.

**Supplementary Figure 4. High expression ratio of M1 to M2 protein in human cells is dependent upon viral M segment host origin.**

293T and MDCK cells were inoculated at a MOI of 5 PFU/cell with PR8 viruses encoding avian or human-derived M segments and incubated at 37°C for 8 h, then cells were lysed. Western immunoblot analysis of virus-infected 293T cells **(A)** and MDCK cells **(G)**. Vinculin expression was measured to allow normalization of viral protein levels. NP expression was measured to assess viral replication. Levels of M1 and M2 protein expression were assessed using an antibody (Mab E10) to a common epitope at the amino terminus of M1 and M2 proteins, allowing relative expression to be assessed. Levels of LC3B I and II were assessed using an antibody that detects both the precursor and activated forms of LC3B protein. **(B, H)** M1 protein and **(C, I)** M2 protein were normalized to vinculin, quantitated and displayed as a percentage of total protein expressed from the M gene. **(D, J)** The ratio of M1:M2 protein expression. **(E, K)** LC3B I protein and **(F, L)** LC3BII protein were normalized, quantitated and displayed as a percentage of total LC3B protein. Graphs in **B-F**, and **H-K** show the means with SD from three independent experiments. For each experiment, two replicate Western immunoblots were performed and quantitated. Statistical significance was assessed using ordinary one-way ANOVA.

**Supplementary Figure 5. High expression ratio of M1 to M2 protein in human cells is dependent upon viral M segment host origin.**

A549 cells were inoculated at a MOI of 5 PFU/cell with PR8 viruses encoding avian or human-derived M segments and incubated at 37°C for 8 h, then cells were lysed. Human M segments were derived from the following viruses: A/NL/602/09 (H1N1) (NL09); A/Panama/2007/99 (H3N2) (Pan99); and A/Bethesda/15 (H3N2) (Beth15). Vinculin expression was measured to allow normalization of viral protein levels. NP expression was measured to assess viral replication. Levels of M1 and M2 protein expression were assessed using an antibody (Mab E10) to a common epitope at the amino terminus of M1 and M2 proteins, allowing relative expression to be assessed. Levels of LC3B I and II were assessed using an antibody that detects both the precursor and the activated forms of LC3B protein. Data presented are representative of Western immunoblots from three independent experiments.

**Supplementary Figure 6. Visualization of M2 localization by immunofluorescence microscopy at 12 h post-infection in A549 and 293T cells.**

A549 **(A, B)**, or 293T **(C, D)** cells were inoculated with the indicated viruses, encoding avian or human M segments, at a MOI of 5 PFU/cell. Cells were fixed at 12 hpi, permeabilised, and stained with anti-M2 (Mab E10; red) and DAPI (blue) followed by imaging with confocal microscopy. Examples of optical sections are shown, either as merged 2-color images or the red and blue channels alone (in grey scale). **(A)** A549 cells with 63x magnification. **(B)** 3x magnification of the same images shown in **A. (C)** 293T cells with 63x magnification. **(D)** 3x magnification of the same images shown in **C.** Brightness was adjusted for optimal clarity, with all images treated equally.

**Supplementary Figure 7. Visualization of LC3 and M2 co-localization by immunofluorescence microscopy.**

A549 cells were transduced with GFP-LC3 protein and inoculated 24 h later with the indicated IAVs, encoding avian- or human-derived M segments, at a MOI of 5 PFU/cell. Cells were fixed 8 h later and stained with anti-M2 (Mab E10; red) and DAPI (blue) followed by imaging with confocal microscopy. Examples of optical sections are shown, either as merged 3-color images or the red, green, and blue channels alone (in grey scale). **(A)** 63x magnification. **(B)** 3x magnification of the same images shown in **A.** Brightness was adjusted for optimal clarity, with all images treated equally. CQ: chloroquine.

**Supplementary Figure 8. Chloroquine treatment results in loss of activation of LC3B.**

A549 cells were inoculated at a MOI of 5 PFU/cell with PR8 viruses encoding avian or human-derived M segments and incubated in the presence or absence of 60 μM chloroquine from 1 hpi. Cells were lysed with whole cell lysis buffer following 8 h incubation at 37°C. Western immunoblot of virus-infected A549 cells: **(A)** Vinculin expression was measured to allow normalization of viral protein levels. NP expression was measured to assess viral replication. Levels of M1 and M2 protein expression were assessed using an antibody (Mab E10) to a common epitope at the amino terminus of M1 and M2 proteins, allowing relative expression to be assessed. Levels of LC3B I and II were assessed using an antibody that detects both the precursor and the activated forms of LC3B protein. Data presented are representative Western immunoblots from three independent experiments. LC3B I protein and LC3B II protein **(B, C)** were normalized, quantitated and displayed as a percentage of total LC3B protein in the absence **(B)** or presence **(C)** of 60 μM chloroquine. Data presented in **B-C**, show the means with SD from three independent experiments. For each experiment, two replicate radiograms were quantitated. Statistical significance was assessed using ordinary two-way ANOVA. CQ: chloroquine.

**Supplementary Figure 9. Schematic depicting chimeric M Segment RNAs**

**(A)** Pandemic H1N1 influenza virus M1 and M2 proteins differ from the avian consensus sequences by 9 residues in M1 and 7 residues in M2. The pH1N1 amino acid identities are indicated in blue at their approximate positions within linear representations of M1 and M2 proteins. The pH1N1 M segment differs from the avian consensus by 8.3% at the nucleotide level. Here, pH1N1 RNA sequence is indicated by blue coloring. **(B-D)** Chimeric human-avian M segments were constructed in which only the non-synonymous changes in the avian consensus were introduced into the A/NL/602/09 (H1N1) M segment. **(E)** Avian consensus M1 and M2 proteins differ from the NL09 sequences by 9 residues in M1 and 7 residues in M2. The avian amino acid identities are indicated in red and avian RNA sequence is indicated by red coloring. **(F-H)** A second set of chimeric M segments was constructed in which the NL09 amino acid identities were introduced into the M segment of A/duck/Alberta/76 (H1N1) virus, yielding segments that encode avian consensus protein(s) but retain much of the nucleotide sequence of the NL09 M segment.

**Supplementary Figure 10. Visualization of LC3 and M2 co-localization in 293T cells infected with viruses carrying avian-human chimeric M segments.**

293T cells were transduced with GFP-LC3 protein and inoculated 24 h later with the indicated IAVs, encoding avian, human or chimeric M segments, at a MOI of 5 PFU/cell. Cells were fixed 12 h later and stained with anti-M2 (Mab E10; red) and DAPI (blue) followed by imaging with confocal microscopy. Examples of optical sections are shown, either as merged 3-color images or the red, green, and blue channels alone (in grey scale). 3x magnification of 63x images are shown. Brightness was adjusted for optimal clarity, with all images treated equally.

**Supplementary Figure 11. Levels of mRNA_7_ (encoding M1) and mRNA_10_ (encoding M2) transcripts in A549 cells infected with viruses carrying human, avian or chimeric M segments.**

A549 cells were inoculated at a MOI of 5 PFU/cell with PR8 NL09 M virus and PR8 Av M virus, along with six PR8-based viruses with chimeric M segments. Cells were incubated at 37°C for 8 h and then lysed with a RNeasy Kit. **(A)** RT primer extension radiogram of virus-infected A549 cells. 5S rRNA levels were measured to allow normalization of viral RNA. Segment 7 vRNA expression was measured to assess viral replication. **(B)** Levels of mRNA_7_, **(C)** mRNA_10_, and **(D)** mRNA_11_ were quantitated and displayed as a percentage of total M gene expressed mRNA. Graphs in **B-D** show the means with SD from three independent experiments. For each experiment, two replicate radiograms were quantitated. Statistical significance was assessed using ordinary one-way ANOVA.

**Supplementary Figure 12. High expression of M2 protein reduces kinetics of replication of PR8-based viruses in guinea pigs.**

Groups of four guinea pigs were inoculated with 10 PFU of each chimeric M-encoding virus, as indicated. Graphs show individual titers obtained from animals used in three independent experiments. Virus replication in nasal wash of inoculated animals was measured by plaque titration at days 2, 4, 6, and 8 post-infection and the titers at each time point were plotted (dotted lines). The differences between PR8 NL09 M 7 mut Av and avian M 9 mut NL encoding viruses were considered significant. Statistical significance in kinetics of growth was determined by assessing the interaction of time and virus using repeated measures, two-way, multiple comparisons ANOVA on mean values, with Bonferroni correction applied to account for comparison of a limited no of means.

**Supplementary Figure 13. PR8 M2 protein is overexpressed and partly localized in perinuclear vesicles in 293T cells.**

293T cells were inoculated with the indicated IAVs, encoding avian-, human- or PR8-derived M segments, at a MOI of 5 PFU/cell, or treated with 60 μM chloroquine. Cells were fixed 8 h later and stained with anti-M2 (Mab E10; red) and DAPI (blue) followed by imaging with confocal microscopy. Examples of optical sections are shown, either as merged 2-color images or the red and blue channels alone (in grey scale). 63x magnification with 3x optical zoom. Brightness was adjusted for optimal clarity, with all images treated equally. CQ: chloroquine.

