## Supplemental figures for "Dysregulation of M segment gene expression contributes to influenza A virus host restriction"

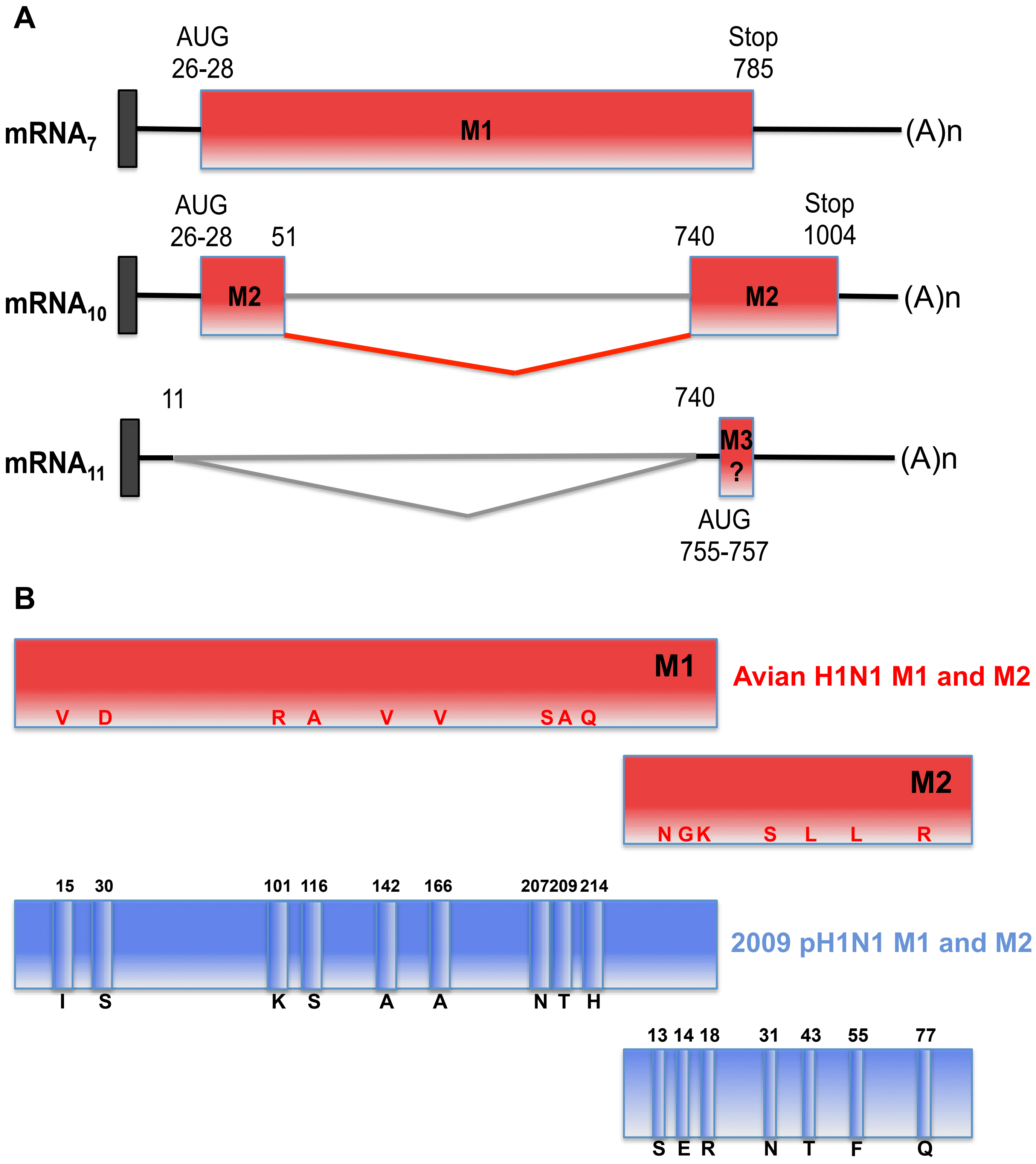

A
AUG
26-28
Stop
785
mRNA7
M1
(A)n
AUG
26-28
51
740
Stop
1004
mRNA10
M2
M2
(A)n
11
740
mRNA11
M3
?
(A)n
AUG
755-757
B
M1
V D R A V V S A Q
Avian H1N1 M1 and M2
M2
N G K S L L R
15 30 101 116 142 166 207 209 214
I S K S A A N T H
2009 pH1N1 M1 and M2
13 14 18 31 43 55 77
S E R N T F Q

C

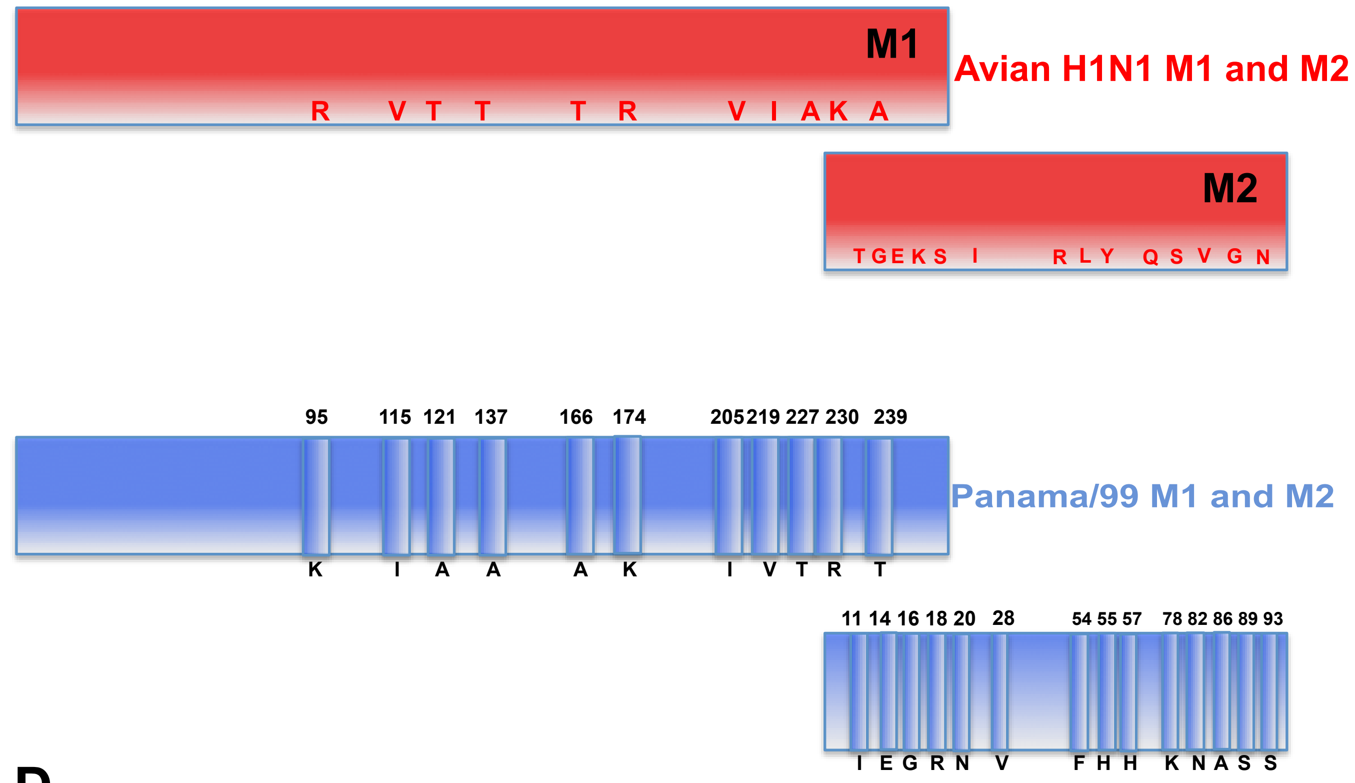


D

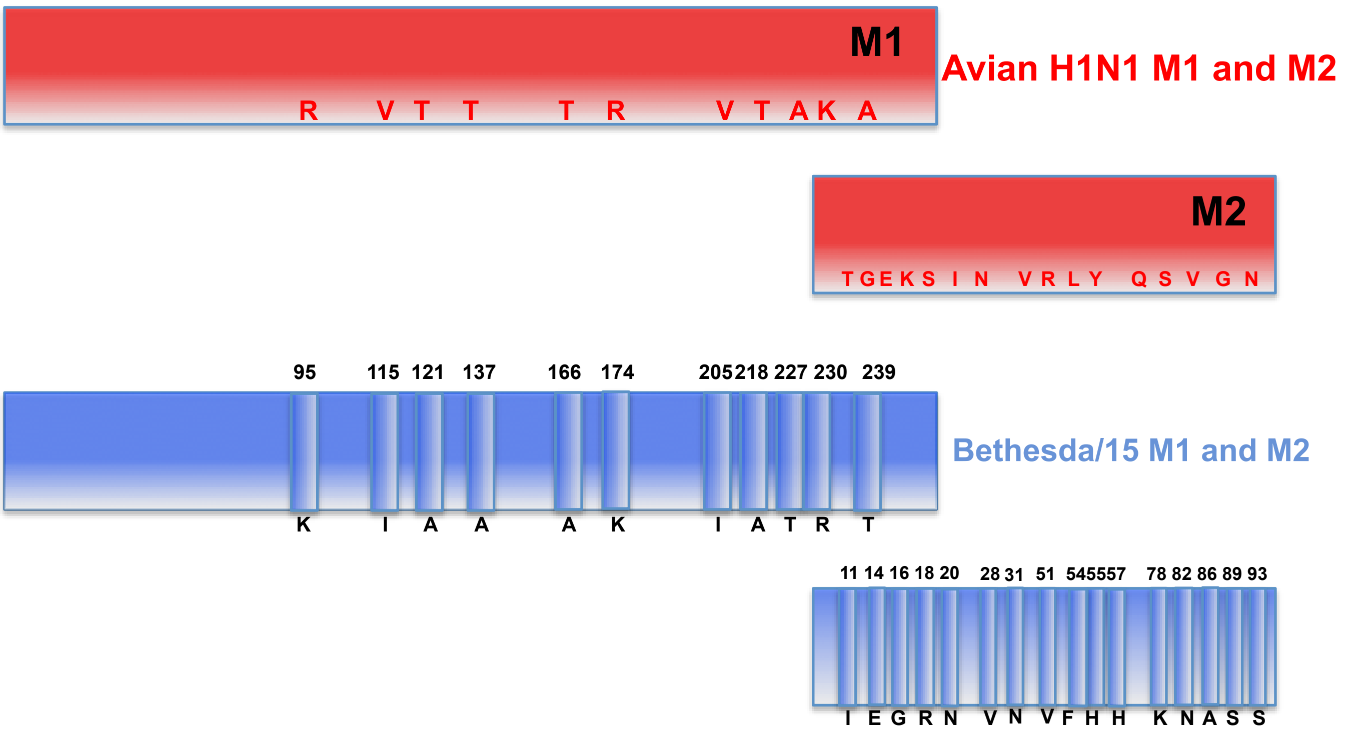


Supplementary Figure 1

**Supplementary Figure 1. Schematic of M segment mRNAs and gene products**

**A.** The M segment of influenza virus is template for synthesis of $mRNA_7$ (encoding M1), mRNA $M_{10}$ (encoding M2), and $mRNA_{11}$ (which encodes a putative but unconfirmed 10 amino acid peptide from a short open reading frame). **B.** Pandemic H1N1 influenza virus M1 and M2 proteins differ from the avian consensus sequences by 9 residues in M1 and 7 residues in M2. The M segments differ by 8.3% at the nucleotide level. **C.** Seasonal H3N2 influenza virus strain A/Panama/2007/99 M1 and M2 proteins differ from the avian consensus sequences by 11 residues in M1 and 14 residues in M2. The M segments differ by 9% at the nucleotide level. **D.** Seasonal H3N2 influenza virus strain A/Bethesda/55/15 M1 and M2 proteins differ from the avian consensus sequences by 11 residues in M1 and 16 residues in M2. The M segments differ by 9.5% at the nucleotide level.

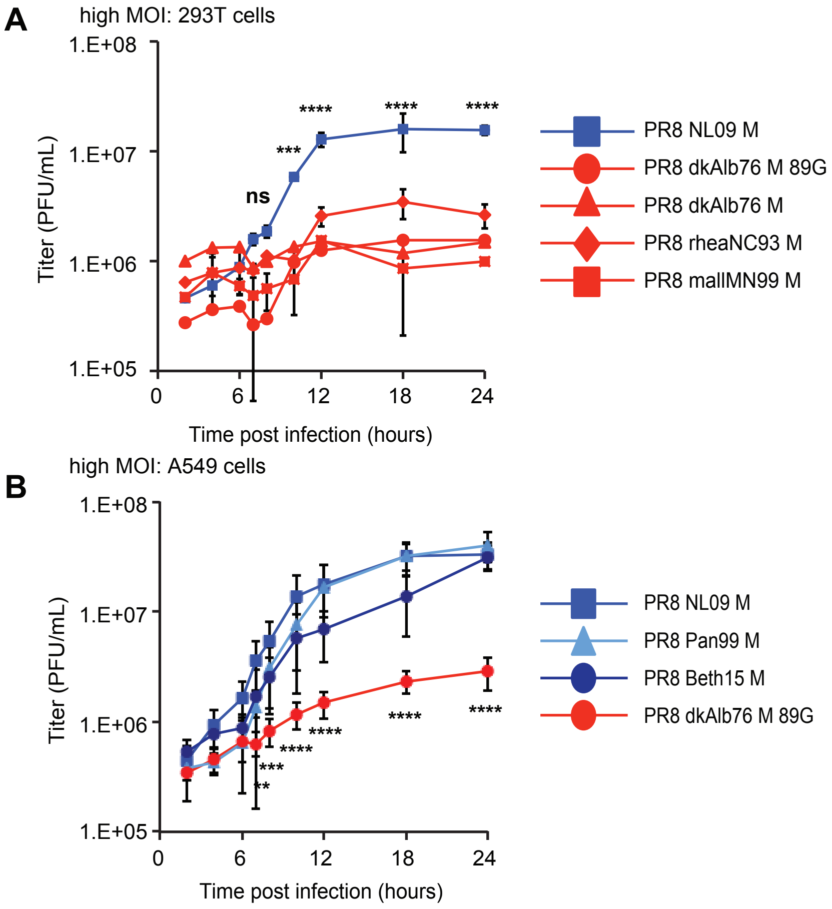


**Supplementary Figure 2. Human Host-derived M Segments Confer Higher Growth to PR8-Based Viruses than Avian Host-derived M Segments in Mammalian Cells at 37 ºC.**

PR8-based viruses were inoculated at an MOI of 5 onto human-derived 293T cells **(A)**, or A549 cells **(B)**. Cells were incubated at 37ºC for up to 24 hours. Virus released into supernatant was collected at the indicated time-points, and virus growth was measured by plaque titration. Data obtained from viruses possessing human M segments are represented with blue lines: A/NL/602/09 M (**A,B**), A/Panama/2007/99 M (**B**), and A/Bethesda/15 M (**B**), while data from viruses encoding avian M segments are represented with red lines. In each cell type, the human M segments conferred more rapid kinetics and higher peak titers of growth than any avian-origin M segment. Single-cycle growth was assessed in three independent experiments, with three technical sample replicates per experiment. Graphs show the means with SD for the three experiments. Statistical significance was determined using repeated measures, two way, multiple ANOVA on log transformed data, with Bonferroni correction applied as there were a limited no of means to compare.

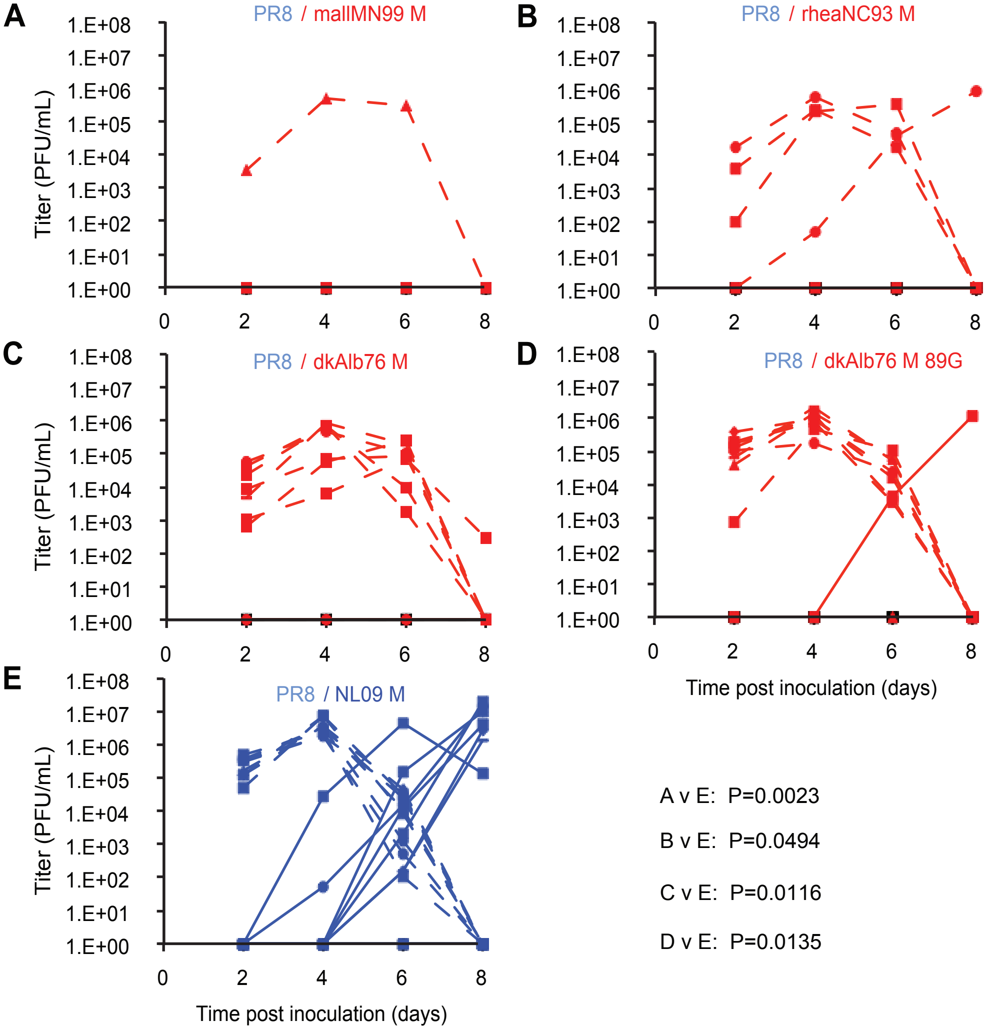


**Supplementary Figure 3. pH1N1 Influenza Virus M Segment Increases Kinetics of Replication of PR8-Based Viruses among Guinea Pigs.**

Groups of four guinea pigs were inoculated with 10 PFU of each avian M-encoding virus, or NL09 M-encoding virus, as indicated. Graphs show individual titers obtained from animals used in three independent experiments. (**A-E**) Virus replication in nasal wash of inoculated animals was measured by plaque titration at days 2, 4, 6, and 8 post-infection and the titers at each time point were plotted (dotted lines). The differences between PR8 NL09 M and each avian M-encoding virus were considered significant. Statistical significance in kinetics of growth was determined by assessing the interaction of time and virus using repeated measures, two way, multiple comparisons ANOVA on mean values, with Bonferroni correction applied to account for comparison of a limited no of means.

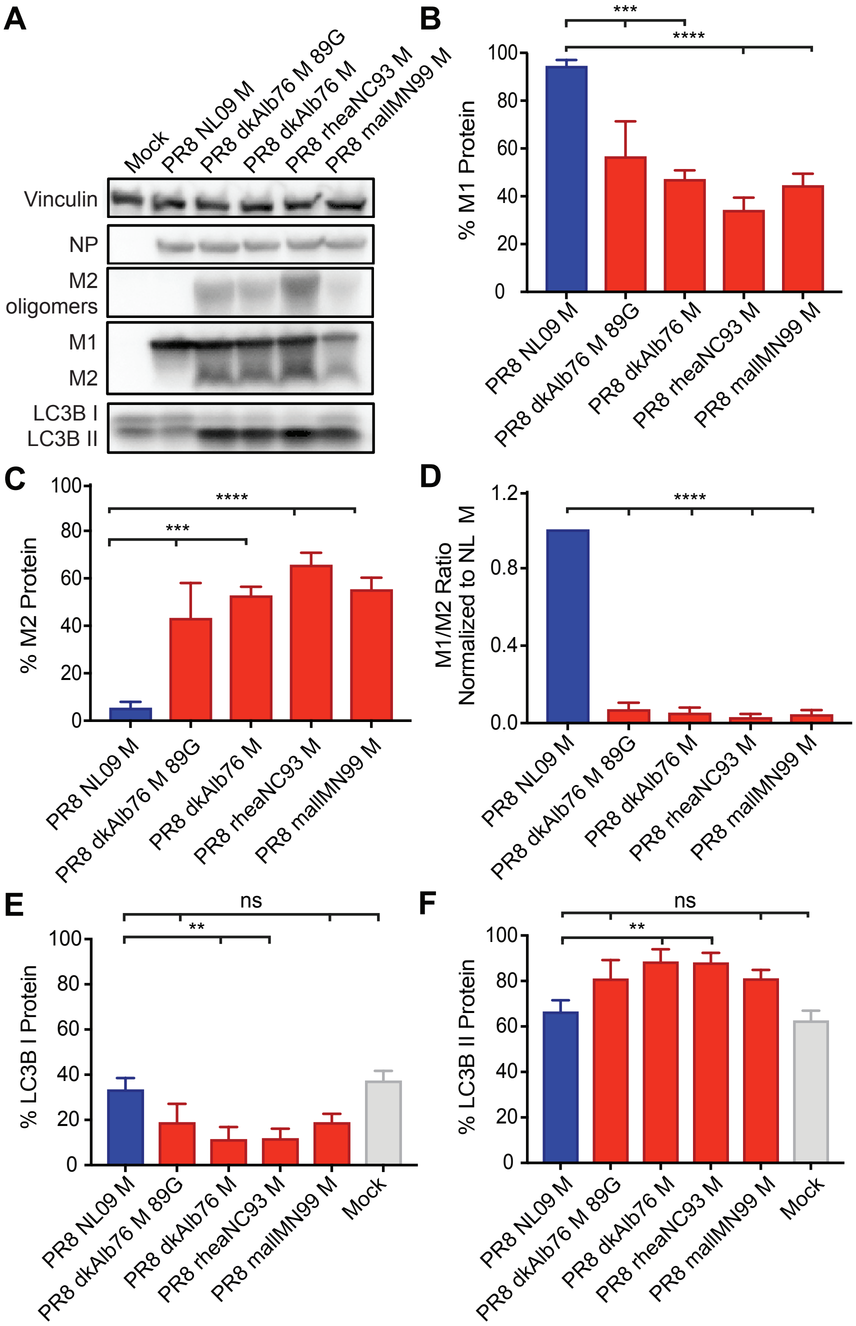

A
Mock
PR8 NL09 M
PR8 dkAlb76 M 89G
PR8 dkAlb76 M
PR8 rheaNC93 M
PR8 mallMN99 M
Vinculin
NP
M2 oligomers
M1
M2
LC3B I
LC3B II
B
% M1 Protein
***
****
C
% M2 Protein
***
****
D
M1/M2 Ratio Normalized to NL M
****
E
% LC3B I Protein
ns
**
Mock
F
% LC3B II Protein
ns
**

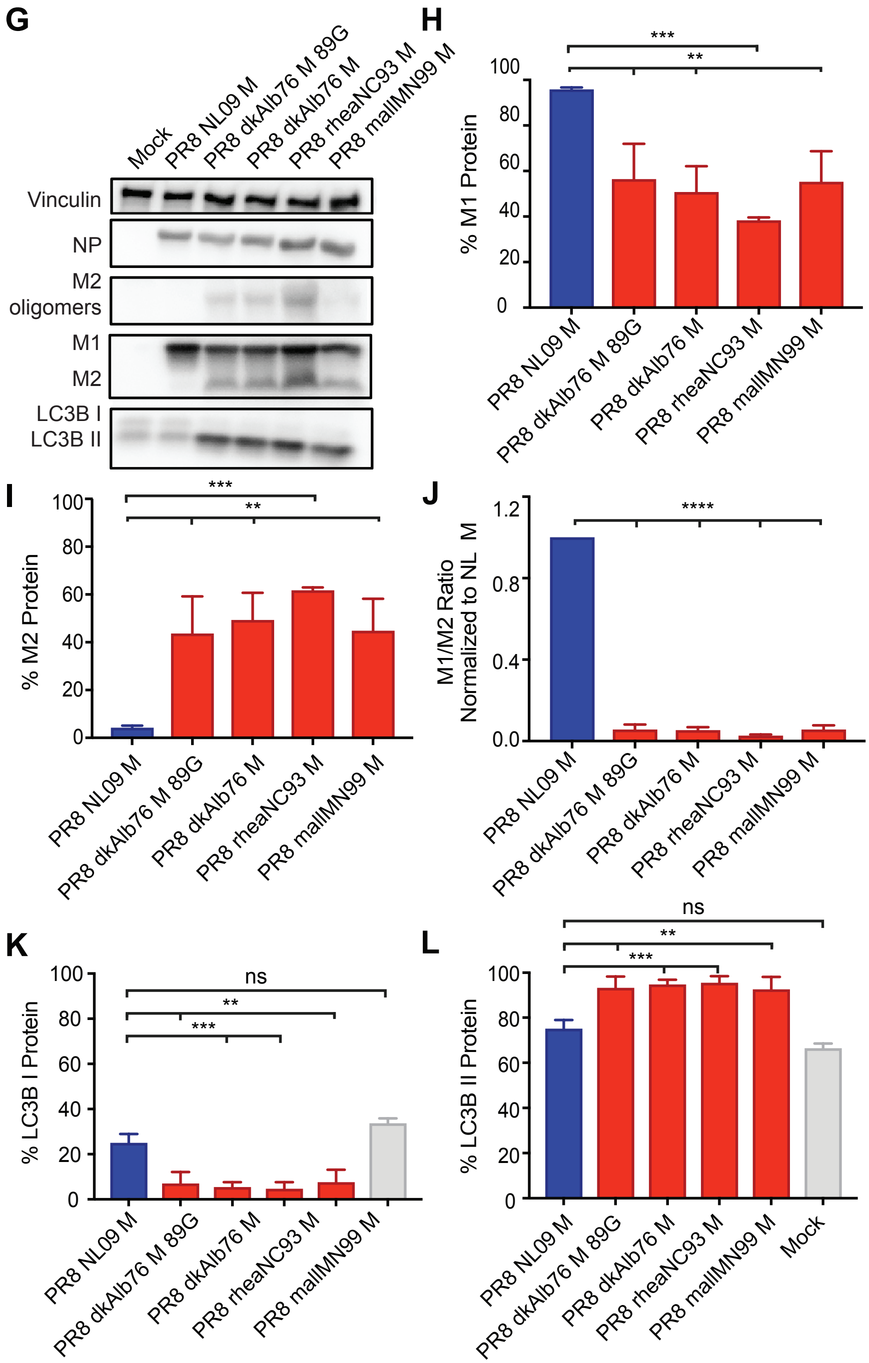


**Supplementary Figure 4. High Expression Ratio of M1 to M2 Protein in Human Cells is Dependent upon Viral M Segment Host Origin.**

293T and MDCK cells were inoculated at MOI=5 with PR8 viruses possessing M segments from human or avian host-derived strains and incubated at 37° C for 8 hours, then cells were lysed with whole cell lysis buffer. Western blot analysis of virus-infected 293T cells **(A)** and MDCK cells **(G)**. Vinculin expression was measured to allow normalization of viral protein levels. NP expression was measured to assess viral replication. Levels of M1 and M2 protein expression were assessed using an antibody (Mab E10) to a common epitope at the amino terminus of M1 and M2 proteins, allowing relative expression to be assessed. Levels of LC3B I and II were assessed using an antibody that detects both the precursor and activated forms of LC3B protein. M1 protein **(B, H)** and M2 protein **(C, I)** were normalized to vinculin, quantitated and displayed as a percentage of total protein expressed from the M gene. The ratio of M1:M2 protein expression was calculated and is plotted in **(D, J)**. LC3B I protein **(E, K)** and LC3B II protein **(F, L)** were normalized, quantitated and displayed as a percentage of total LC3B protein. Graphs in **B-F**, and **H-K** show the means with SD from three independent experiments. For each experiment, two replicate Western blots were performed and quantitated. Statistical significance was assessed using ordinary one-way ANOVA.

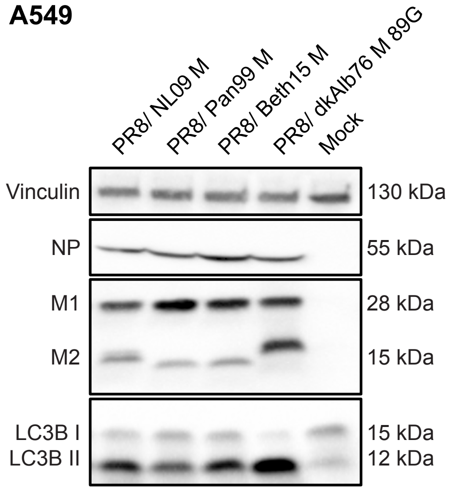


**Supplementary Figure 5. High Expression Ratio of M1 to M2 protein in Human Cells is Dependent upon Viral M Segment Host Origin.**

A549 cells were inoculated at MOI=5 with PR8 viruses possessing M segments from human or avian host-derived strains and incubated at 37º C for 8 hours, then cells were lysed with whole cell lysis buffer. Human M segments were derived from the following viruses: A/NL/602/09N (H1N1) (NL09); A/Panama/2007/99 (H3N2) (Pan99); and A/Bethes-da/15 (H3N2) (Beth15). Vinculin expression was measured to allow normalization of viral protein levels. NP expression was measured to assess viral replication. Levels of M1 and M2 protein expression were assessed using an antibody (Mab E10) to a common epitope at the amino terminus of M1 and M2 proteins, allowing relative expression to be assessed. Levels of LC3B I and II were assessed using an antibody that detects both the precursor and the activated forms of LC3B protein. Data presented are representative of Western blots from three independent experiments.

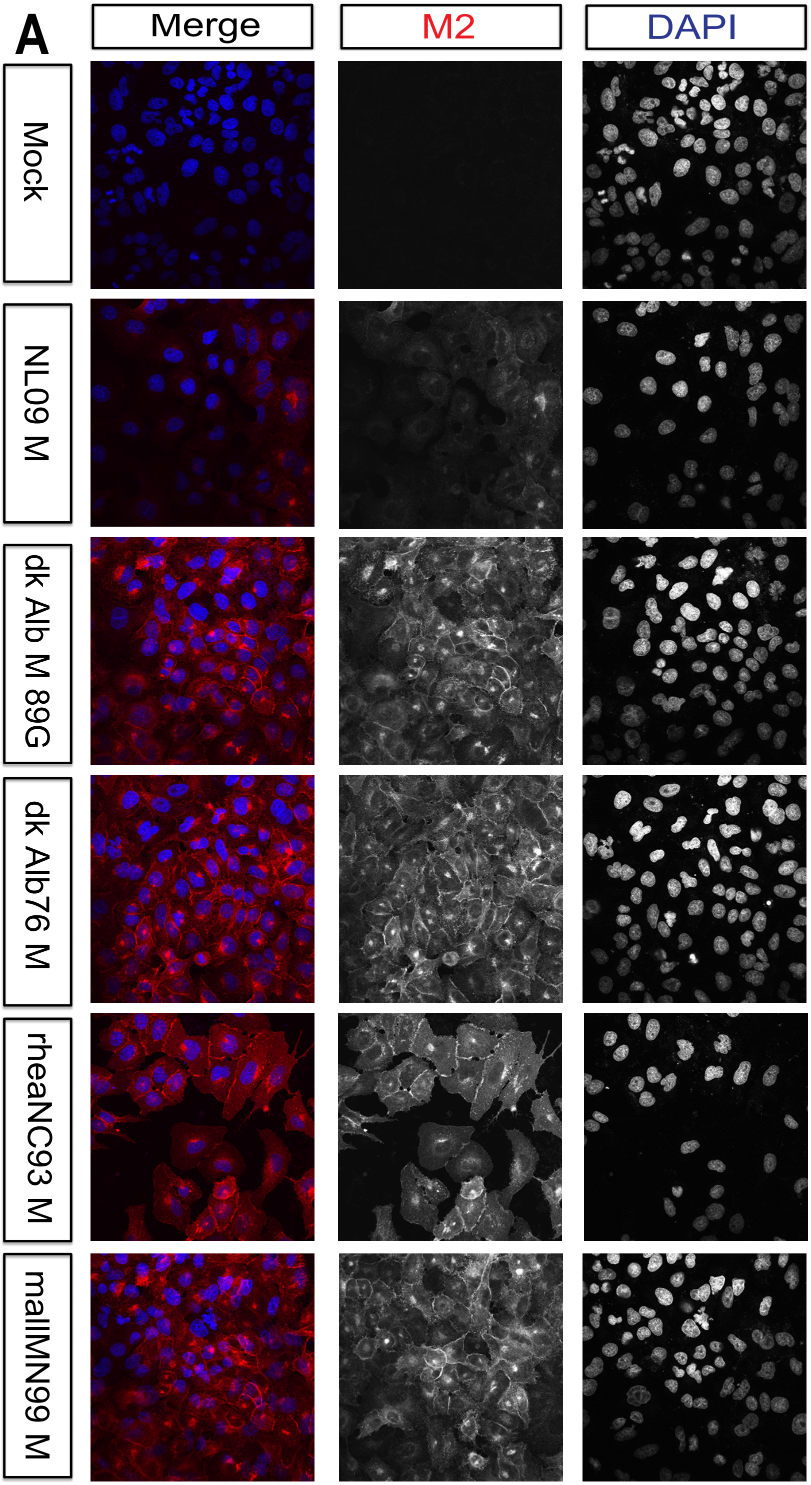
A
Merge
M2
DAPI
Mock
NL09 M
dk Alb M 89G
dk Alb76 M
rheaNC93 M
mallMN99 M

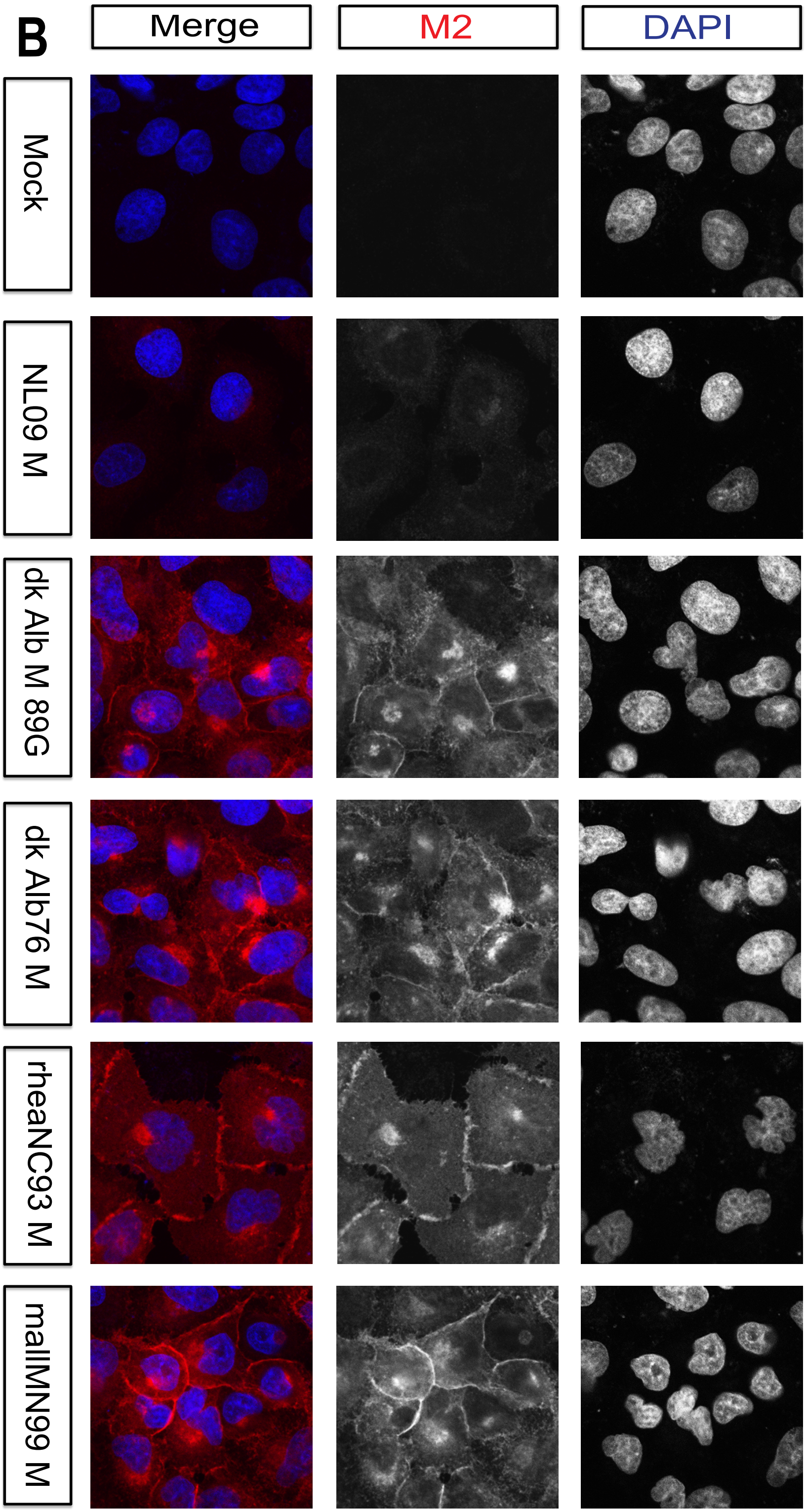
B
Merge
M2
DAPI
Mock
NL09 M
dk Alb M 89G
dk Alb76 M
rheaNC93 M
mallMN99 M

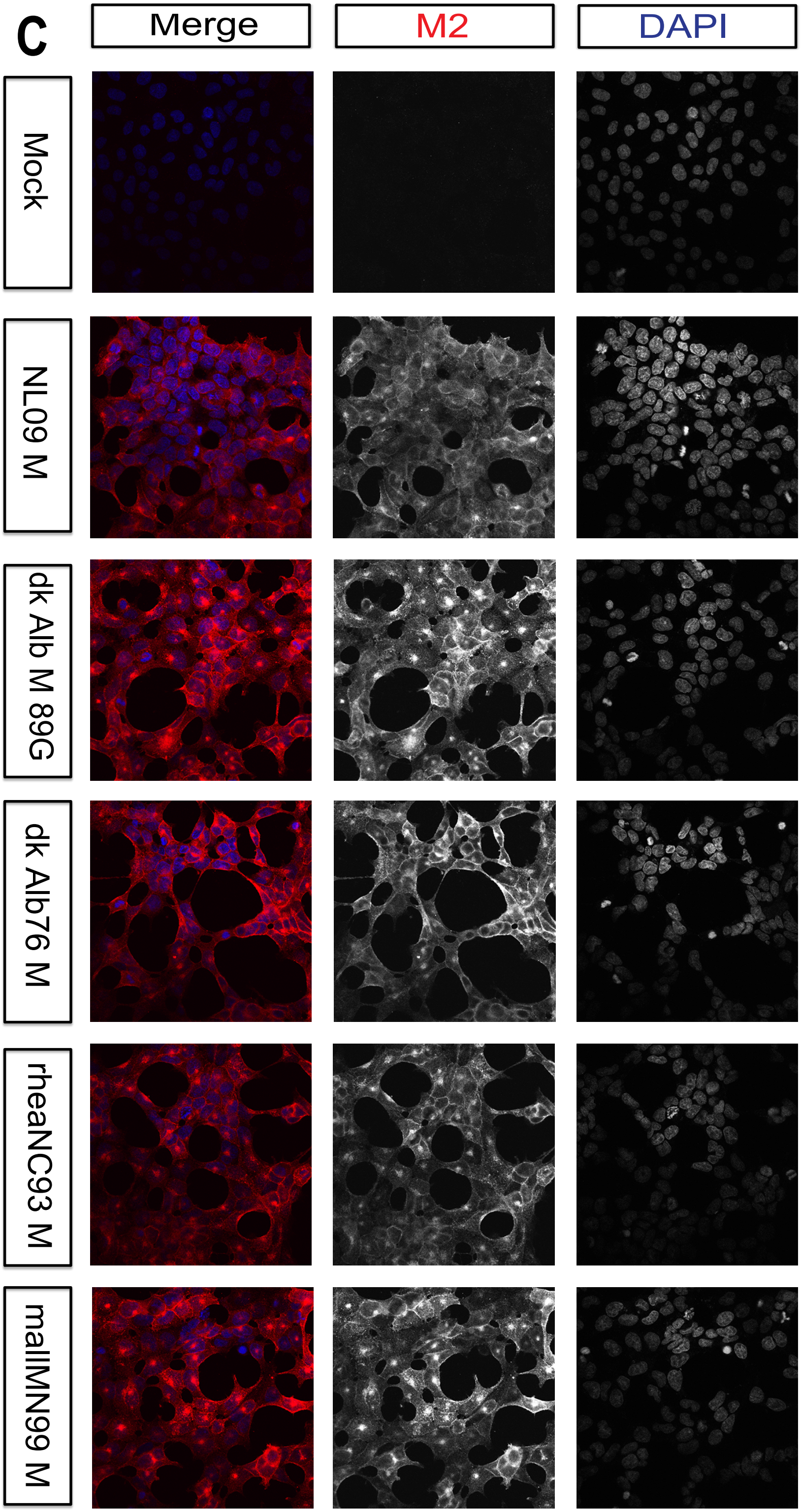
C
Merge
M2
DAPI
Mock
NL09 M
dk Alb M 89G
dk Alb76 M
rheaNC93 M
mallMN99 M

**Supplementary Figure 6. Visualization of M2 localization by immunofluorescence microscopy at 12 h post-infection in A549, and 293T cells.**

A549 (**A, B**), or 293T (**C, D**) cells were inoculated with the indicated viruses, encoding avian or human M segments, at an MOI of 5 PFU/cell. Cells were fixed at 12 h p.i., permeabilised, and stained with anti-M2 (Mab E10; red) and DAPI (blue) followed by imaging with confocal microscopy. Examples of optical sections are shown, either as merged 2-color images or the red and blue channels alone (in grey scale). **(A)** A549 cells with 63x magnification. **(B)** 3x magnification of the same images shown in **A**. **(C)** 293T cells with 63x magnification. **(D)** 3x magnification of the same images shown in **C**. **(D)** A549 cells with 63x magnification. **(E)** 3x magnification of the same images shown in **D**. Brightness was adjusted for optimal clarity, with all images treated equally.

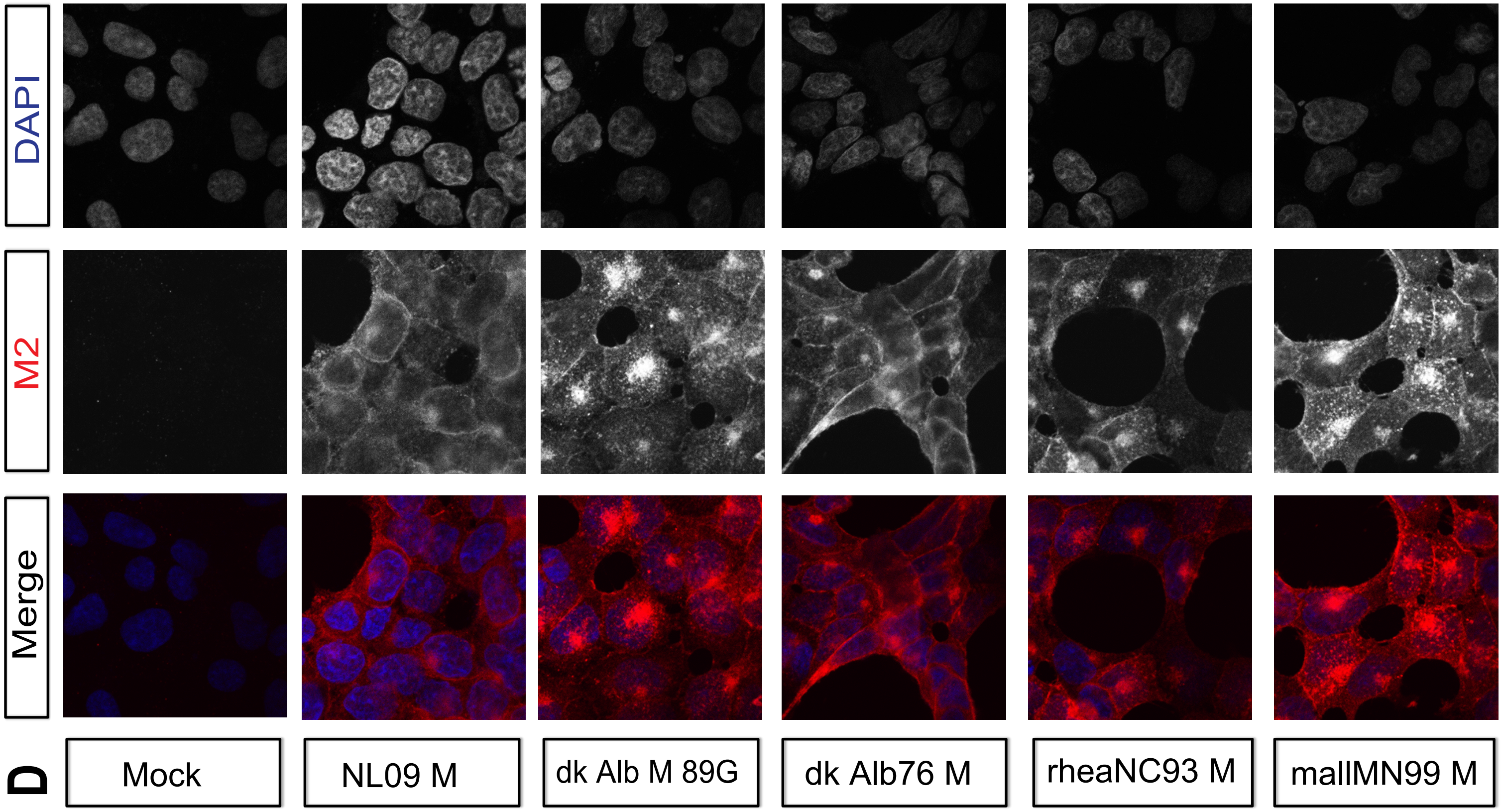

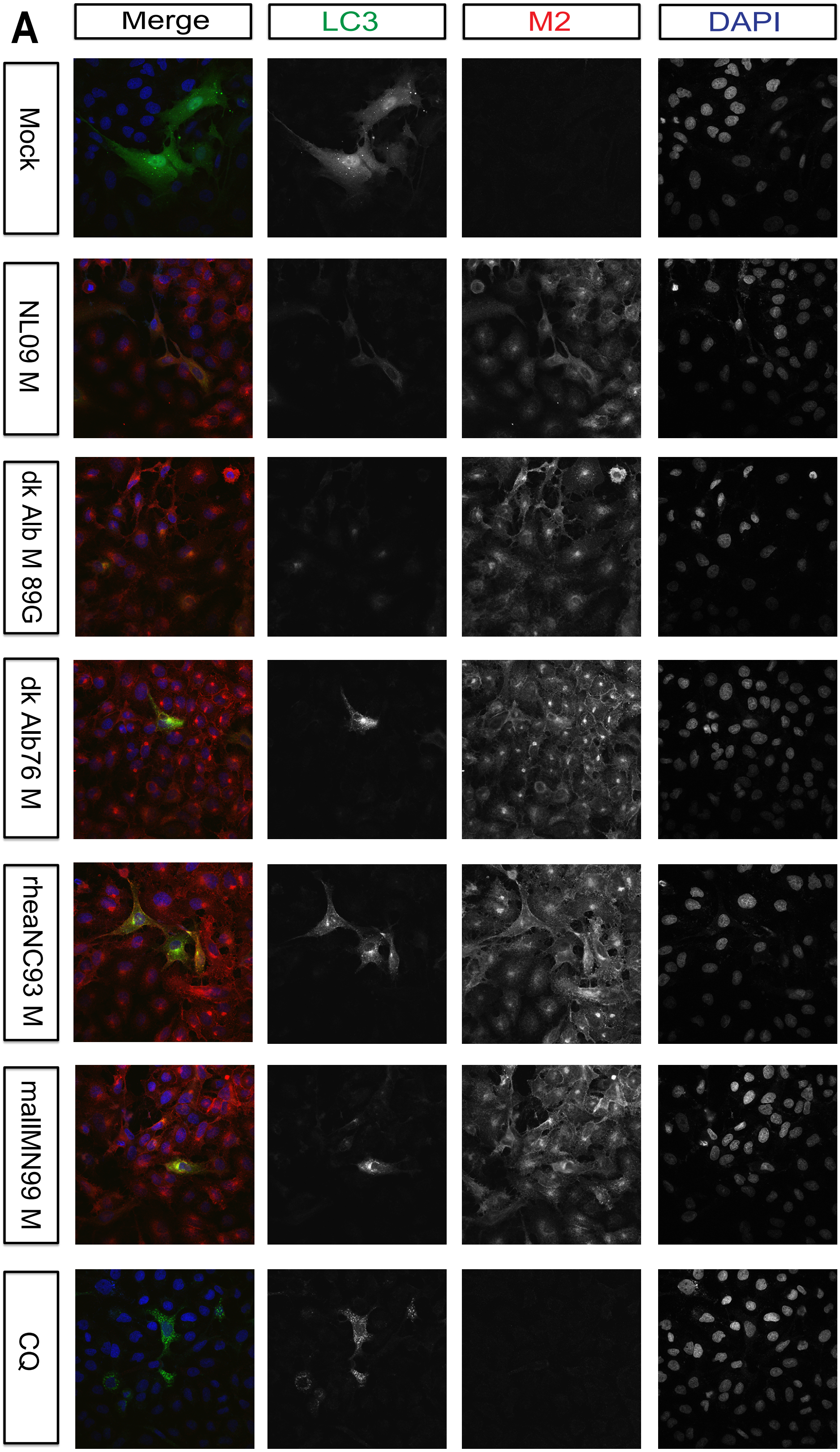
A
Merge
LC3
M2
DAPI
Mock
NL09 M
dk Alb M 89G
dk Alb76 M
rheaNC93 M
mallMN99 M
CQ

**Supplementary Figure 7. Visualization of LC3 and M2 co-localization by immunofluorescence microscopy.**

A549 cells were transduced with GFP-LC3 protein and inoculated 24 h later with the indicated IAVs, encoding avian or human derived M segments, at an MOI of 5 PFU/cell. Cells were fixed 8 h later and stained with anti-M2 (Mab E10; red) and DAPI (blue) followed by imaging with confocal microscopy. Examples of optical sections are shown, either as merged 3-color images or the red, green, and blue channels alone (in grey scale). **(A)** 63x magnification. **(B)** 3x magnification of the same images shown in **A**. Brightness was adjusted for optimal clarity, with all images treated equally.

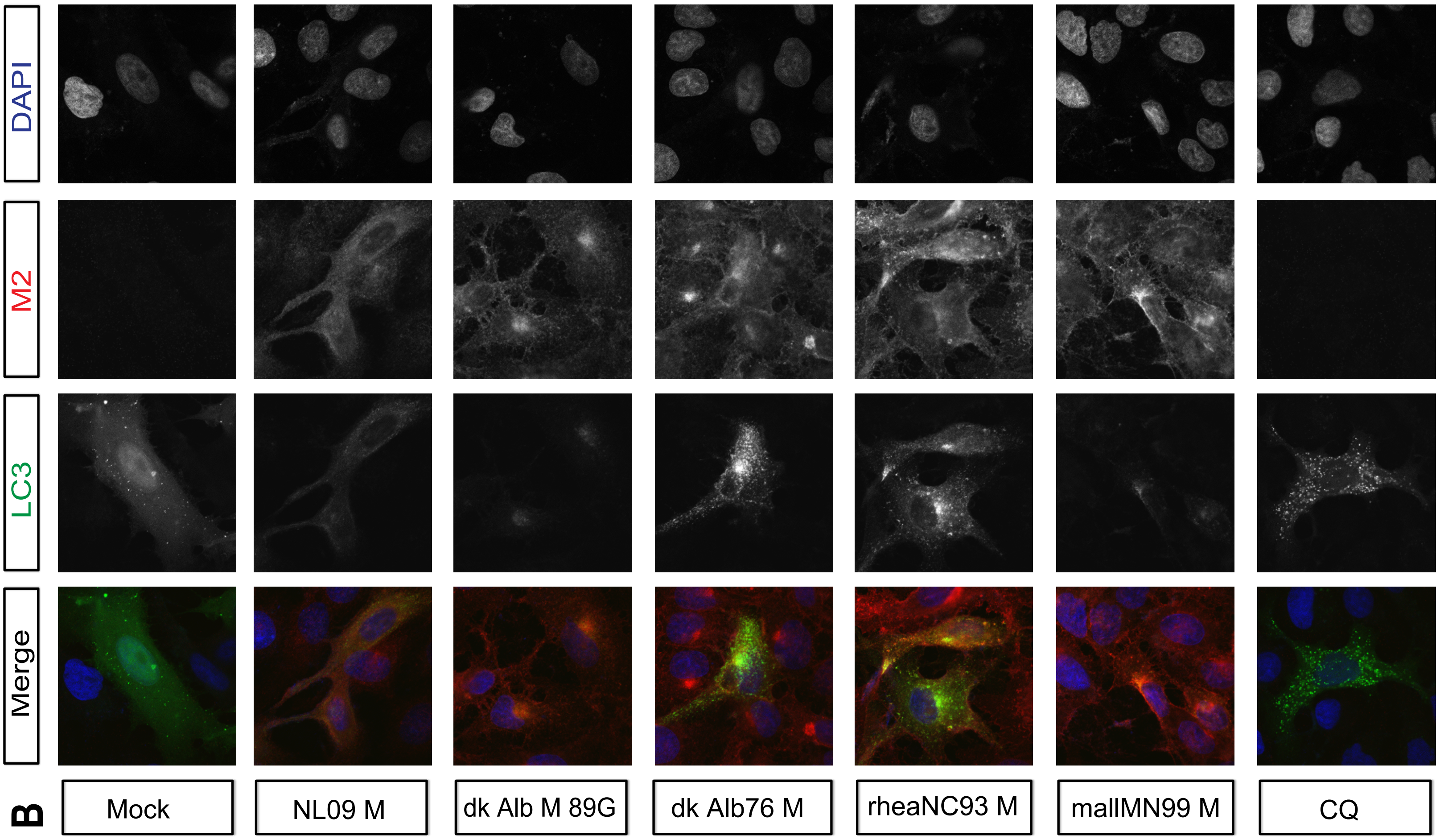

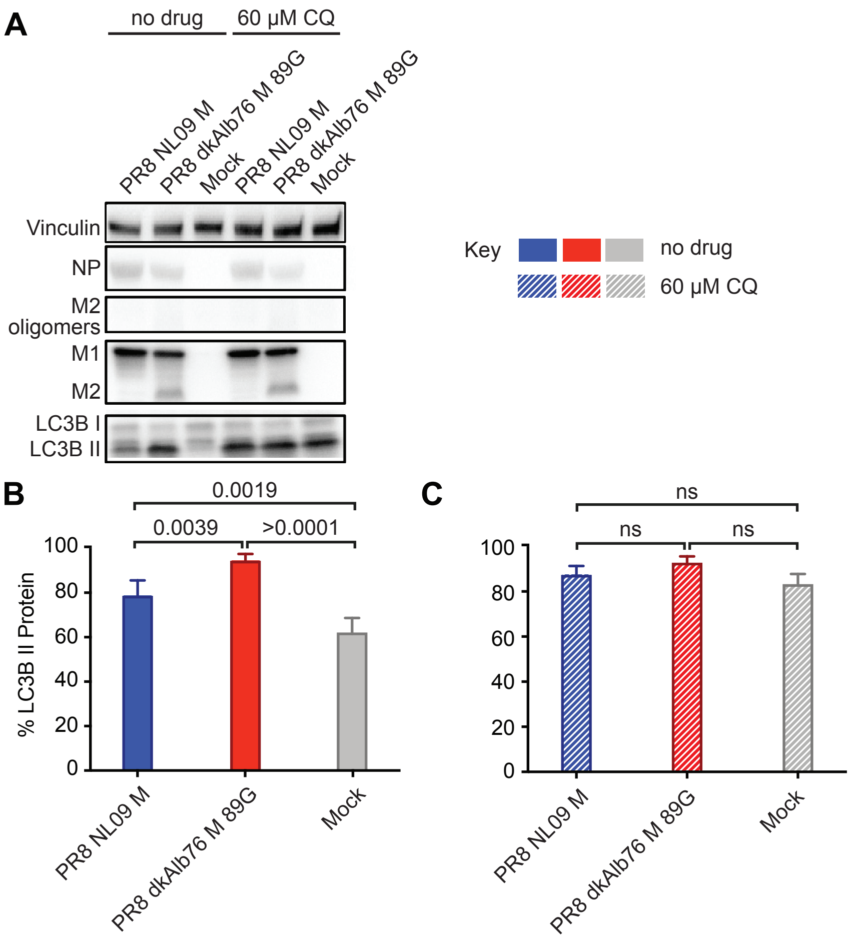


**Supplementary Figure 8. Chloroquine treatment results in loss of activation of LC3B.**

A549 cells were inoculated at MOI=5 with PR8 viruses possessing M segments from human or an avian host-derived strain and incubated in the presence or absences of 60   M chloroquine from 1 hpi. Cells were lysed with whole cell lysis buffer following 8 hours incubation at 37° C. Western blot of virus-infected A549 cells **(A)**. Vinculin expression was measured to allow normalization of viral protein levels. NP expression was measured to assess viral replication. Levels of M1 and M2 protein expression were assessed using an antibody (Mab E10) to a common epitope at the amino terminus of M1 and M2 proteins, allowing relative expression to be assessed. Levels of LC3B I and II were assessed using an antibody that detects both the precursor and the activated forms of LC3B protein. Data presented are representative Western blots from three independent experiments. LC3B I protein and LC3B II protein **(B, C)** were normalized, quantitated and displayed as a percentage of total LC3B protein in the absence **(B)** or presence **(C)** of 60 µM chloroquine. Data presented in **B-C**, show the means with SD from three independent experiments. For each experiment, two replicate radiograms were quantitated. Statistical significance was assessed using ordinary two-way ANOVA.

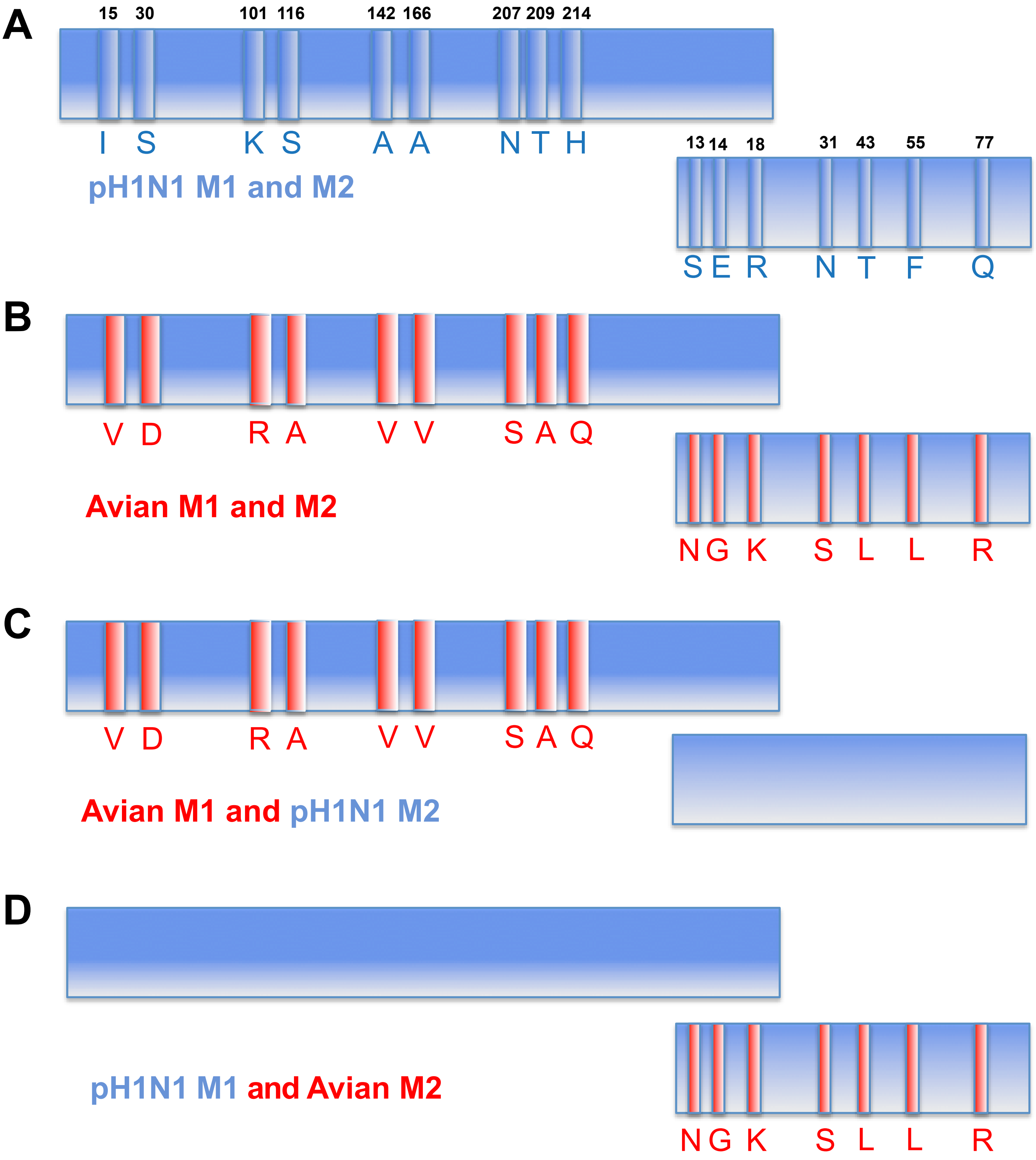
A
15 30 101 116 142 166 207 209 214
I S K S A A N T H
pH1N1 M1 and M2
13 14 18 31 43 55 77
S E R N T F Q
B
V D R A V V S A Q
Avian M1 and M2
N G K S L L R
C
V D R A V V S A Q
Avian M1 and pH1N1 M2
D
pH1N1 M1 and Avian M2
N G K S L L R

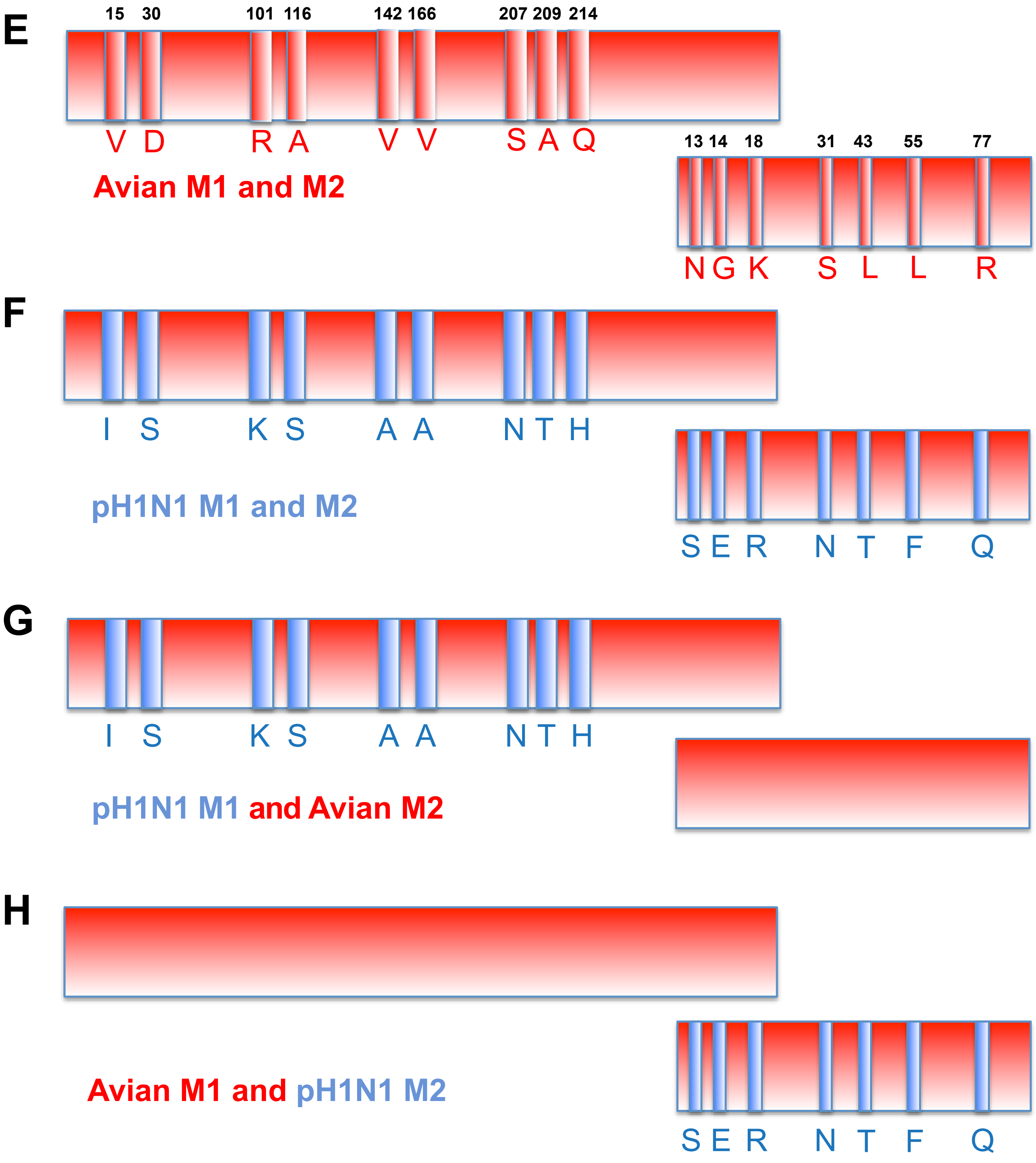


Supplementary Figure 9

**Supplementary Figure 9. Schematic depicting chimeric M Segment RNAs**

**(A)** Pandemic H1N1 influenza virus M1 and M2 proteins differ from the avian consensus sequences by 9 residues in M1 and 7 residues in M2. The pH1N1 amino acid identities are indicated in blue at their approximate positions within linear representations of M1 and M2 proteins. The pH1N1 M segment differs from the avian consensus by 8.3% at the nucleotide level. Here, pH1N1 RNA sequence is indicated by blue coloring. **(B-D)** Chimeric human-avian M segments were constructed in which only the non-synonymous changes in the avian consensus were introduced into the A/NL/602/09 (H1N1) M segment. **(E)** Avian consensus M1 and M2 proteins differ from the NL/09 sequences by 9 residues in M1 and 7 residues in M2. The avian amino acid identities are indicated in red and avian RNA sequence is indicated by red coloring. **(F-H)** A second set of chimeric M segments was constructed in which the NL/09 amino acid identities were introduced into the M segment of A/duck/Alberta/76 (H1N1) virus, yielding segments that encode avian consensus protein(s) but retain much of the nucleotide sequence of the NL/09 M segment.

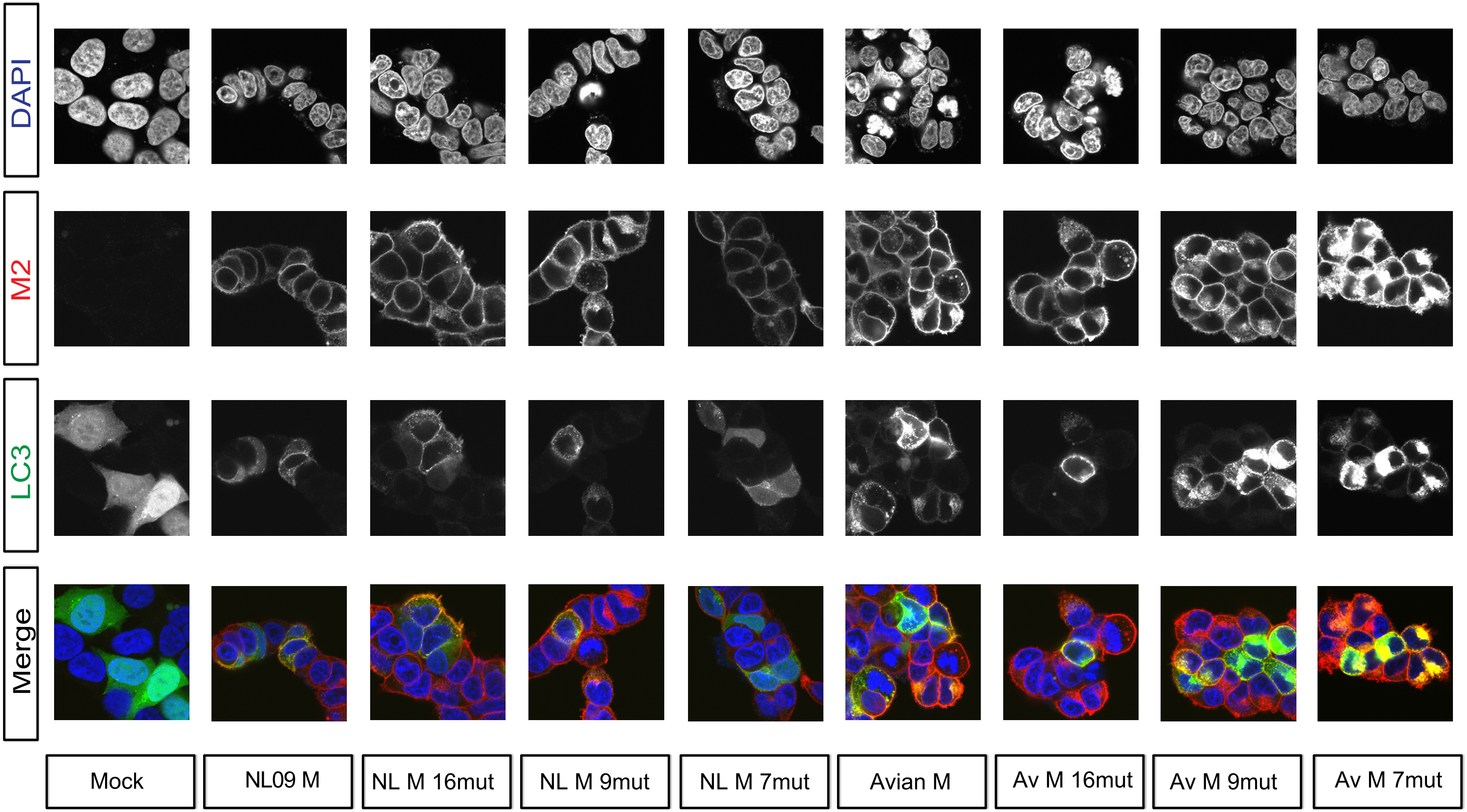


**Supplementary Figure 10. Visualization of LC3 and M2 co-localization in 293T cells infected with viruses carrying avian-human chimeric M segments.**

293T cells were transduced with GFP-LC3 protein and inoculated 24 h later with the indicated IAVs, encoding avian, human or chimeric M segments, at an MOI of 5 PFU/cell. Cells were fixed 12 h later and stained with anti-M2 (Mab E10; red) and DAPI (blue) followed by imaging with confocal microscopy. Examples of optical sections are shown, either as merged 3-color images or the red, green, and blue channels alone (in grey scale). 3x magnification of 63x images are shown. Brightness was adjusted for optimal clarity, with all images treated equally.

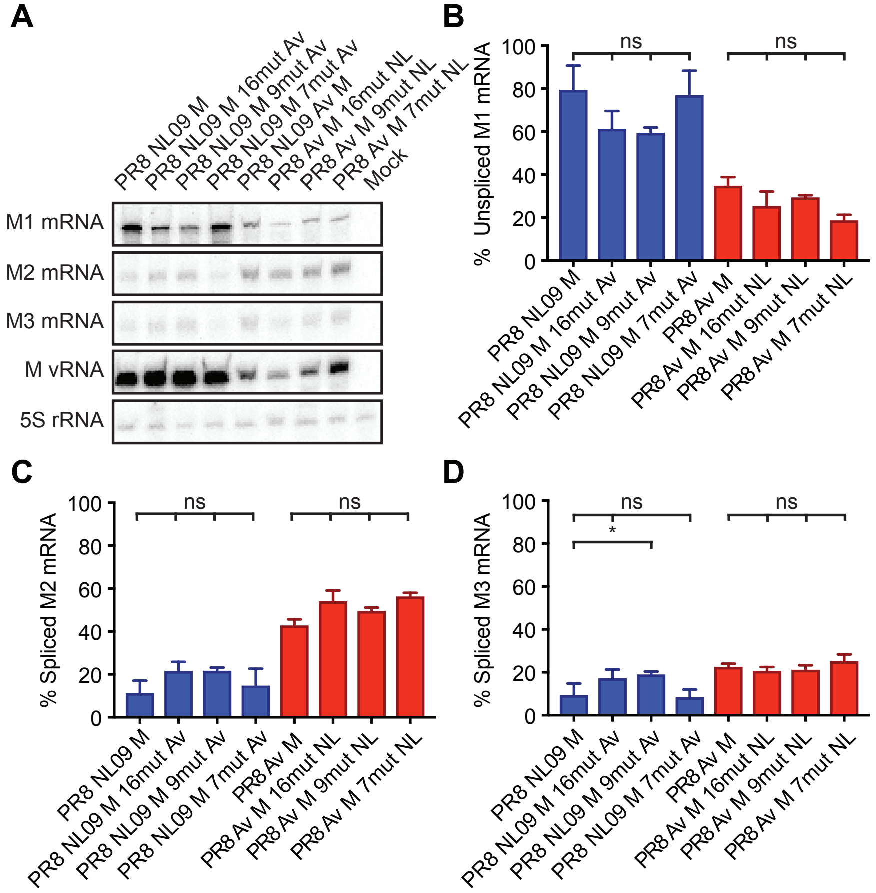


**Supplementary Figure 11. Levels of mRNA7 (encoding M1) and mRNA10 (encoding M2) transcripts in A549 cells infected with viruses carrying human, avian or chimeric M segments.**

A549 cells were inoculated at an MOI=5 with PR8 NL09 M virus and PR8 Av M virus, along with six PR8-based viruses with chimeric M segments. Cells were incubated at 37° C for 8 hours and then lysed with an RNeasy Kit. **(A)** RT primer extension radiogram of virus-infected A549 cells. 5S rRNA levels were measured to allow normalization of viral RNA. Segment 7 vRNA expression was measured to assess viral replication. Levels of mRNA7 **(B)**, mRNA10 **(C)**, and mRNA11 **(D)** were quantitated and displayed as a percentage of total M gene expressed mRNA. Graphs in B-D show the means with SD from three independent experiments. For each experiment, two replicate radiograms were quantitated. Statistical significance was assessed using ordinary one-way ANOVA.

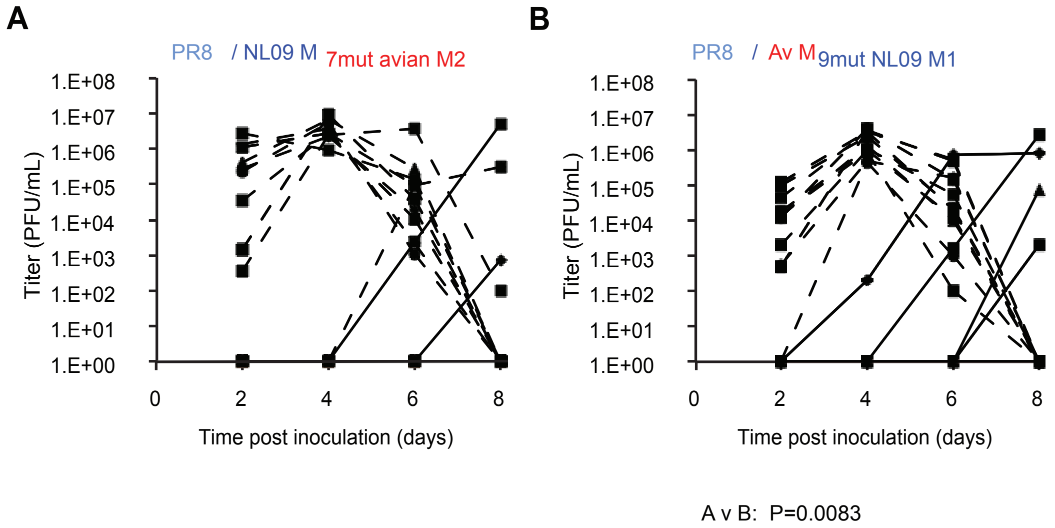


**Supplementary Figure 12. High Expression of M2 Protein Reduces Kinetics of Replication of PR8-Based Viruses in Guinea Pigs.**

Groups of four guinea pigs were inoculated with 10 PFU of each chimeric M-encoding virus, as indicated. Graphs show individual titers obtained from animals used in three independent experiments. Virus replication in nasal wash of inoculated animals was measured by plaque titration at days 2, 4, 6, and 8 post-infection and the titers at each time point were plotted (dotted lines). The differences between PR8 NL09 M 7 mut Av and avian M 9 mut NL encoding viruses were considered significant. Statistical significance in kinetics of growth was determined by assessing the interaction of time and virus using repeated measures, two way, multiple comparisons ANOVA on mean values, with Bonferroni correction applied to account for comparison of a limited no of means.

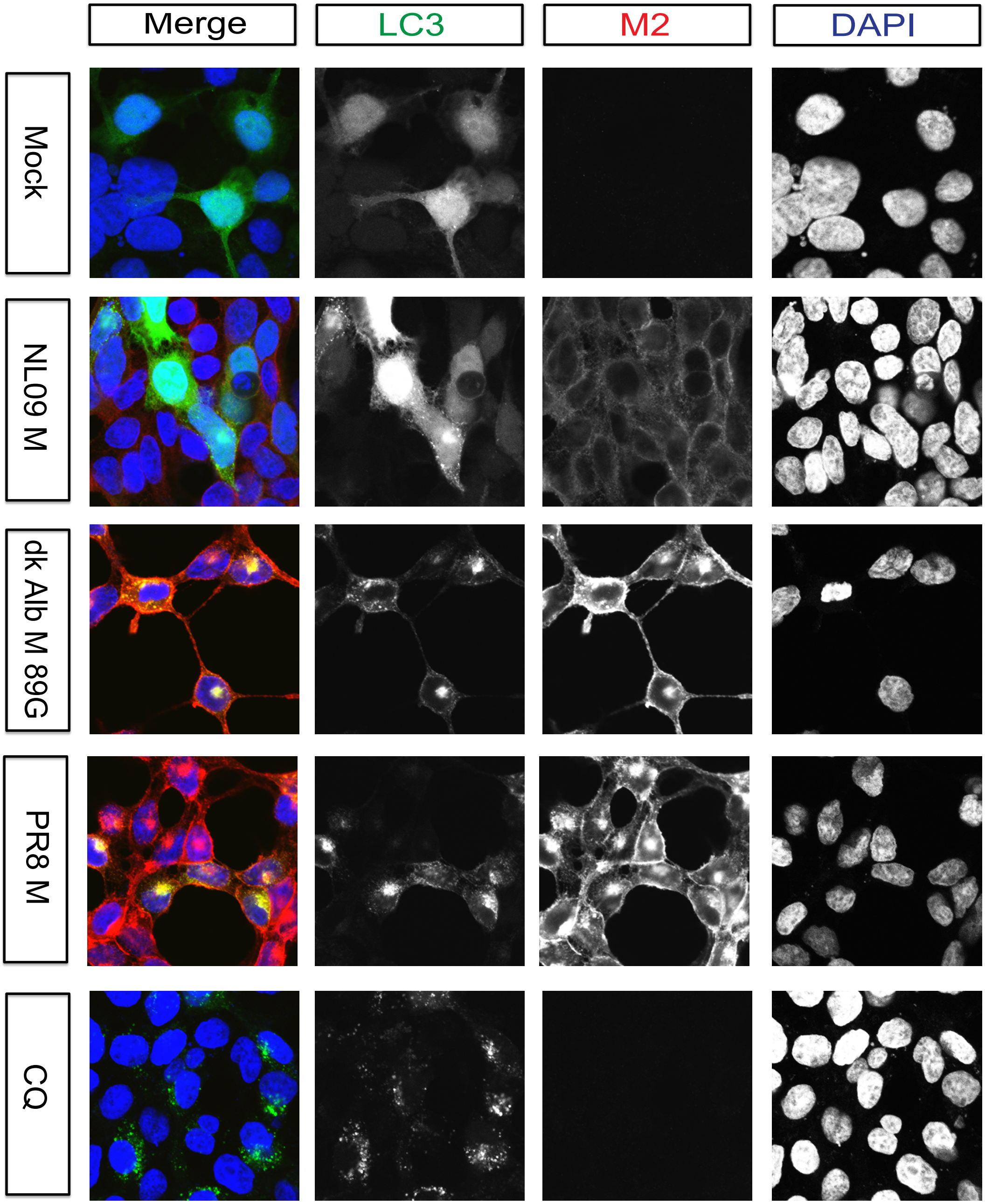


**Supplementary Figure 13. PR8 M2 protein  is over-expressed and partly localized in perinuclear vesicles in 293T cells**

293T cells were inoculated with the indicated IAVs, encoding avian, human or PR8 derived M segments, at an MOI of 5 PFU/cell, or treated with 60uM chloroquine. Cells were fixed 8 h later and stained with anti-M2 (Mab E10; red) and DAPI (blue) followed by imaging with confocal microscopy. Examples of optical sections are shown, either as merged 2-color images or the red and blue channels alone (in grey scale). 63x magnification with 3x optical zoom. Brightness was adjusted for optimal clarity, with all images treated equally.
